# Designed proteins assemble antibodies into modular nanocages

**DOI:** 10.1101/2020.12.01.406611

**Authors:** Robby Divine, Ha V. Dang, George Ueda, Jorge A. Fallas, Ivan Vulovic, William Sheffler, Shally Saini, Yan Ting Zhao, Infencia Xavier Raj, Peter A. Morawski, Madeleine F. Jennewein, Leah J. Homad, Yu-Hsin Wan, Marti R. Tooley, Franzika Seeger, Ali Etemadi, Mitchell L. Fahning, James Lazarovits, Alex Roederer, Alexandra C. Walls, Lance Stewart, Mohammadali Mazloomi, Neil P. King, Daniel J. Campbell, Andrew T. McGuire, Leonidas Stamatatos, Hannele Ruohola-Baker, Julie Mathieu, David Veesler, David Baker

**Affiliations:** Department of Biochemistry, University of Washington, Seattle, WA 98195, USA; Institute for Protein Design, University of Washington, Seattle, WA 98195, USA; Institute for Stem Cell and Regenerative Medicine, University of Washington, Seattle, WA 98109, USA; Oral Health Sciences, School of Dentistry, University of Washington, Seattle, WA 98195, USA; Benaroya Research Institute, Seattle, WA 98101, USA; Fred Hutchinson Cancer Research Center, Vaccines and Infectious Diseases Division, Seattle, WA, USA; Medical Biotechnology Department, School of Advanced Technologies in Medicine, Tehran University of Medical Sciences (TUMS), Tehran, Iran; University of Washington, Department of Global Health, Seattle, WA, USA; Department of Comparative Medicine, University of Washington, Seattle, WA 98195, USA; Howard Hughes Medical Institute, University of Washington, Seattle, WA 98195, USA

## Abstract

Antibodies are widely used in biology and medicine, and there has been considerable interest in multivalent antibody formats to increase binding avidity and enhance signaling pathway agonism. However, there are currently no general approaches for forming precisely oriented antibody assemblies with controlled valency. We describe the computational design of two-component nanocages that overcome this limitation by uniting form and function. One structural component is any antibody or Fc fusion and the second is a designed Fc-binding homo-oligomer that drives nanocage assembly. Structures of 8 antibody nanocages determined by electron microscopy spanning dihedral, tetrahedral, octahedral, and icosahedral architectures with 2, 6, 12, and 30 antibodies per nanocage match the corresponding computational models. Antibody nanocages targeting cell-surface receptors enhance signaling compared to free antibodies or Fc-fusions in DR5-mediated apoptosis, Tie2-mediated angiogenesis, CD40 activation, and T cell proliferation; nanocage assembly also increases SARS-CoV-2 pseudovirus neutralization by α-SARS-CoV-2 monoclonal antibodies and Fc-ACE2 fusion proteins. We anticipate that the ability to assemble arbitrary antibodies without need for covalent modification into highly ordered assemblies with different geometries and valencies will have broad impact in biology and medicine.

## Introduction

Antibodies are widely used therapeutic and diagnostic protein tools that are central to modern biotechnology, with the market for antibody-based technologies reaching $150 billion in 2019 (*1*). To increase binding avidity, and to enhance agonism through receptor clustering, there has been considerable interest in high valency antibody formats that present more than two antigen-binding sites (*2*, *3*). Current techniques for creating multivalent antibody-presenting formats include chaining together multiple antigen-binding fragments (*4*, *5*), pentameric immunoglobulin M (IgM) or IgM derivatives such as fragment crystallizable (Fc) domain hexamers (*6*), inorganic materials fused to multiple dimeric immunoglobulin G (IgG) antibodies (*7*), or protein oligomers or nanoparticles to which immunoglobulin (Ig) or Ig-binding domains are linked (*8*–*13*). While these approaches are effective at multimerizing antibodies, they often require extensive engineering or multiple-step conjugation reactions for each new desired antibody oligomer. In the case of nanoparticles with flexibly linked Ig-binding domains, it is difficult to ensure full IgG occupancy on the particle surface and to prevent particle flocculation induced when multiple nanoparticles bind to dimeric IgGs. To our knowledge, no methods currently exist for creating antibody-based protein nanoparticles across multiple valencies with precisely-controlled geometry and composition that are applicable to the vast number of off-the-shelf IgG antibodies.

We set out to design proteins that drive the assembly of arbitrary antibodies into symmetric assemblies with well-defined structures. Previous design efforts have successfully built nanocages by computationally fusing (*14*, *15*) or docking together (*16*, *17*) protein building blocks with cyclic symmetry so that the symmetry axes of the building blocks align with a larger target architecture. For example, an I52 icosahedral assembly is built by bringing together a pentamer and a dimer that align to the icosahedral five- and two-fold symmetry axes, respectively. We reasoned that symmetric protein assemblies could also be built out of IgG antibodies, which are two-fold symmetric proteins, by placing the symmetry axes of the antibodies on the two-fold axes of the target architecture and designing a second protein to hold the antibodies in the correct orientation.

### A general computational method for antibody cage design

We set out to design an antibody-binding, nanocage-forming protein that precisely arranges IgG dimers along the two-fold symmetry axes of a target architecture. We sought to accomplish this by rigidly fusing together three types of “building block” proteins: antibody Fc-binding proteins, monomeric helical linkers, and cyclic oligomers; each building block plays a key role in the final fusion protein. The Fc-binder forms the first nanocage interface between the antibody and the nanocage-forming design, the cyclic homo-oligomer forms the second nanocage interface between designed protein chains, and the monomer links the two interfaces together in the correct orientation for nanocage formation. The designed cage-forming protein is thus a cyclic oligomer terminating in antibody-binding domains that bind IgG antibodies at the orientations required for the proper formation of antibody nanocages (hereafter AbCs, for <u>A</u>ntibody <u>C</u>ages).

Key to the success of this fusion approach is a sufficiently large set of building blocks to fuse, and possible fusion sites per building block, to meet the rather stringent geometric criteria (described below) required to form the desired symmetric architecture. We used protein A (*18*), which recognizes the Fc domain of the IgG constant region, as one of two antibody-binding building blocks, and designed a second Fc-binding building block by grafting the protein A interface residues onto a previously designed helical repeat protein (Fig. S1) (*18*, *19*). Our final library consisted of these 2 Fc-binding proteins (*18*), 42 *de novo* designed helical repeat protein monomers (*19*), and between 1-3 homo-oligomers depending on geometry (2 C2s, 3 C3s, 1 C4, and 1 C5) (*20*, *21*). An average of roughly 150 residues were available for fusion per protein building block, avoiding all positions at the Fc or homo-oligomer protein interface, leading to on the order of 10^7^ possible tripartite (i.e., Fc-binder/monomer/homo-oligomer) fusions. For each of these tripartite fusions, the rigid body transform between the internal homo-oligomeric interface and the Fc-binding interface is determined by the shapes of each of its three building blocks and the locations and geometry of the “junctions” that link them into a single subunit.

We used a computational protocol that rapidly samples all possible fusions from our building block library to identify those with the net rigid body transforms required to generate dihedral, tetrahedral, octahedral, and icosahedral AbCs (*14*, *15*). To describe the final nanocage architectures, we follow a naming convention which summarizes the point group symmetry and the cyclic symmetries of the building blocks (*16*). For example, a T32 assembly has tetrahedral point group symmetry and is built out of a C3 cyclic symmetric antibody-binding designed oligomer, and the C2 cyclic symmetric antibody Fc. While the antibody dimer aligns along the two-fold axis in all architectures, the designed component is a second homodimer in D2 dihedral structures; a homotrimer in T32 tetrahedral structures, O32 octahedral structures, and I32 icosahedral structures; a homotetramer in O42 octahedral structures; and a homopentamer in I52 icosahedral structures.

To make the fusions, the protocol first aligns the model of the Fc and Fc-binder protein along the C2 axis of the specified architecture (Fig. 1a-b). The Fc-binder is then fused to a monomer, which is in turn fused to a homo-oligomer. Rigid helical fusions are made by superimposing residues in alpha helical secondary structure from each building block; in the resulting fused structure one building block chain ends and the other begins at the fusion point, forming a new, continuous alpha helix (Fig. 1c). To drive formation of the desired nanocage architecture, fusions are made such that the antibody two-fold axis and the symmetry axis of the homo-oligomer intersect at specified angles at the center of the architecture (Fig. 1d). To generate D2 dihedral, T32 tetrahedral, O32 or O42 octahedral, and I32 or I52 icosahedral nanocages, the required intersection angles are 90.0°, 54.7°, 35.3°, 45.0°, 20.9°, and 31.7°, respectively (*22*). We allowed angular and distance deviations from the ideal architecture of at most 5.7° and 0.5 Å, respectively (see Methods). Candidate fusion models were further filtered based on the number of contacts around the fusion junction (to gauge structural rigidity) and clashes between backbone atoms. Next, the amino acid identities and conformations around the newly formed building block junction were optimized using the SymPackRotamersMover in Rosetta to maintain the rigid fusion geometry required for assembly (Fig. 1e). Following sequence design, we selected for experimental characterization six D2 dihedral, eleven T32 tetrahedral, four O32 octahedral, two O42 octahedral, fourteen I32 icosahedral, and eleven I52 icosahedral designs predicted to form AbCs (Fig. 1f).

**Figure 1.**
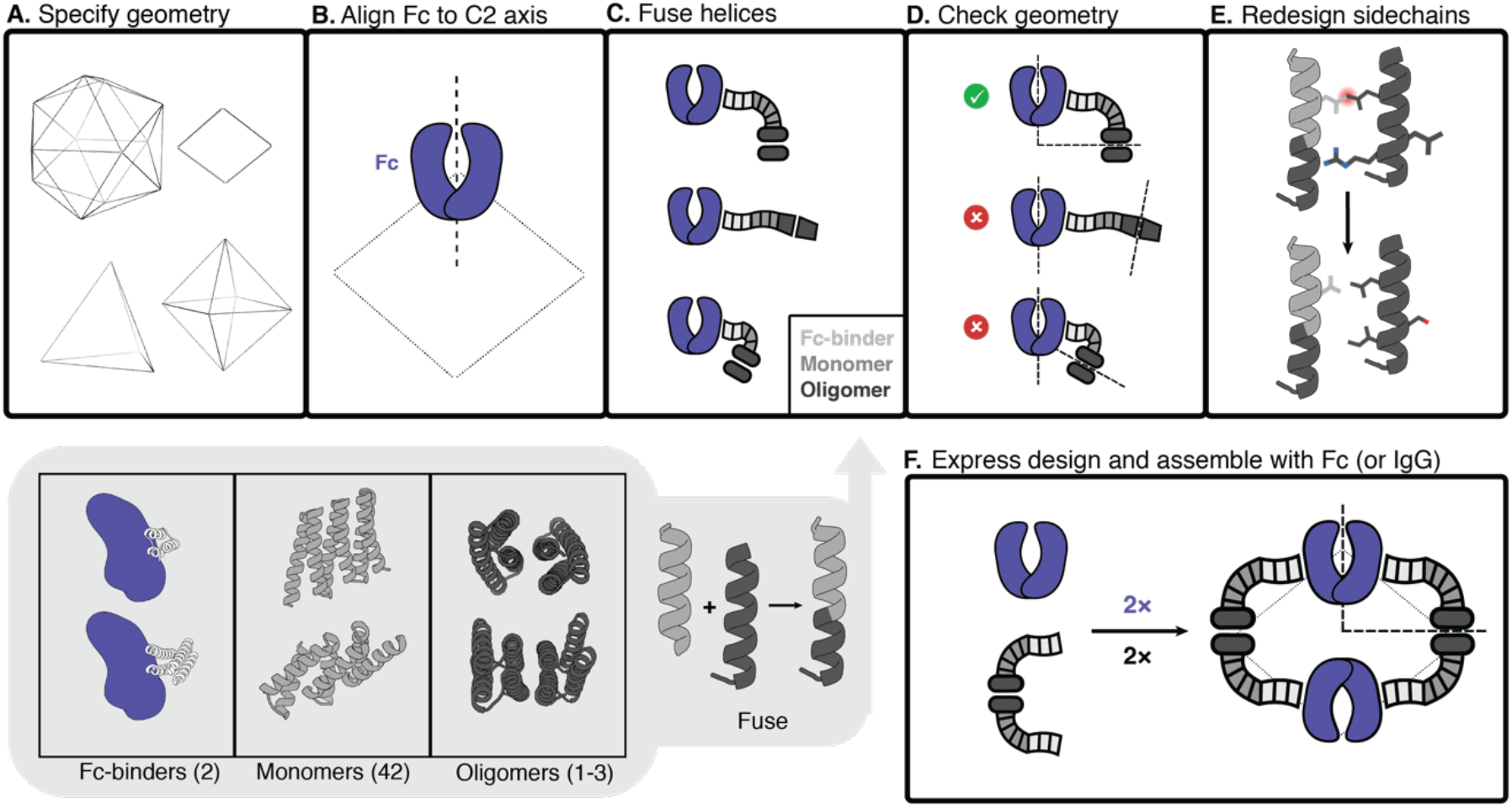
Antibody nanocage (AbC) design. **A,** Polyhedral geometry is specified. Clockwise from top left: icosahedral, dihedral, octahedral, and tetrahedral geometries are shown. **B,** An antibody Fc model from hIgG1 is aligned to one of the C2 axes (in this case, a D2 dihedron is shown). **C,** Antibody Fc-binders are fused to helical repeat proteins that are then fused to the monomeric subunit of helical cyclic oligomers. All combinations of building blocks and building block junctions are sampled (below inset, grey). **D,** Tripartite fusions are checked to ensure successful alignment of the C2 Fc symmetry axes with that of the polyhedral architecture (in the case of the D2 symmetry shown here, the C2 axes must intersect at a 90° angle). **E,** Fusions that pass the geometric criteria move forward with sidechain redesign, where e.g. amino acids are optimized to ensure that core-packing residues are nonpolar and solvent-exposed residues are polar. **F,** Designed AbC-forming oligomers are bacterially expressed, purified, and assembled with antibody Fc or IgG.

### Structural characterization

Synthetic genes encoding designed protein sequences appended with a C-terminal 6×histidine tag were expressed in *E. coli*. Designs were purified from clarified lysates using immobilized metal affinity chromatography (IMAC), and size exclusion chromatography (SEC) was used as a final purification step. Across all geometries, 34 out of 48 AbC-forming designs had a peak on SEC that roughly corresponded to the expected size of the design model (Fig. S2, Table S1). Designs were then combined with human IgG1 Fc, and the assemblies were purified via SEC.

Eight of these AbC-forming designs readily self-assembled after mixing with Fc into a species that eluted as a monodisperse peak at a volume consistent with the target nanoparticle molecular size (Fig. 2a-b; 3 D2 dihedral, 2 T32 tetrahedral, 1 O42 octahedral, and 2 I52 icosahedral AbCs). For the i52.6 design, adding 100 mM L-arginine to the assembly buffer prevented aggregation after combining with Fc (*23*); all other designs readily self-assembled in Tris-buffered saline. Of these eight AbC-forming designs, all designs expressed well, with SEC-purified protein yields between 50-100 mg protein/L of bacterial culture. After combining with Fc, at least 90% of the protein injected on SEC is recovered in the assembly (left-most) peak (Fig. 2b). SEC peaks for the T32 and O42 designs were somewhat broader than other designs, spanning 3-4 mL in retention volume, as observed in previous nanocage design efforts (*16*). The I52 designs eluted in the void volume, consistent with their predicted diameters. Most other designs still bound Fc, as evidenced by SEC or by visibly precipitating with Fc after combination, but did not form monodisperse nanoparticles by SEC (Table S1), perhaps because of deviations from the target fusion geometry.

**Figure 2.**
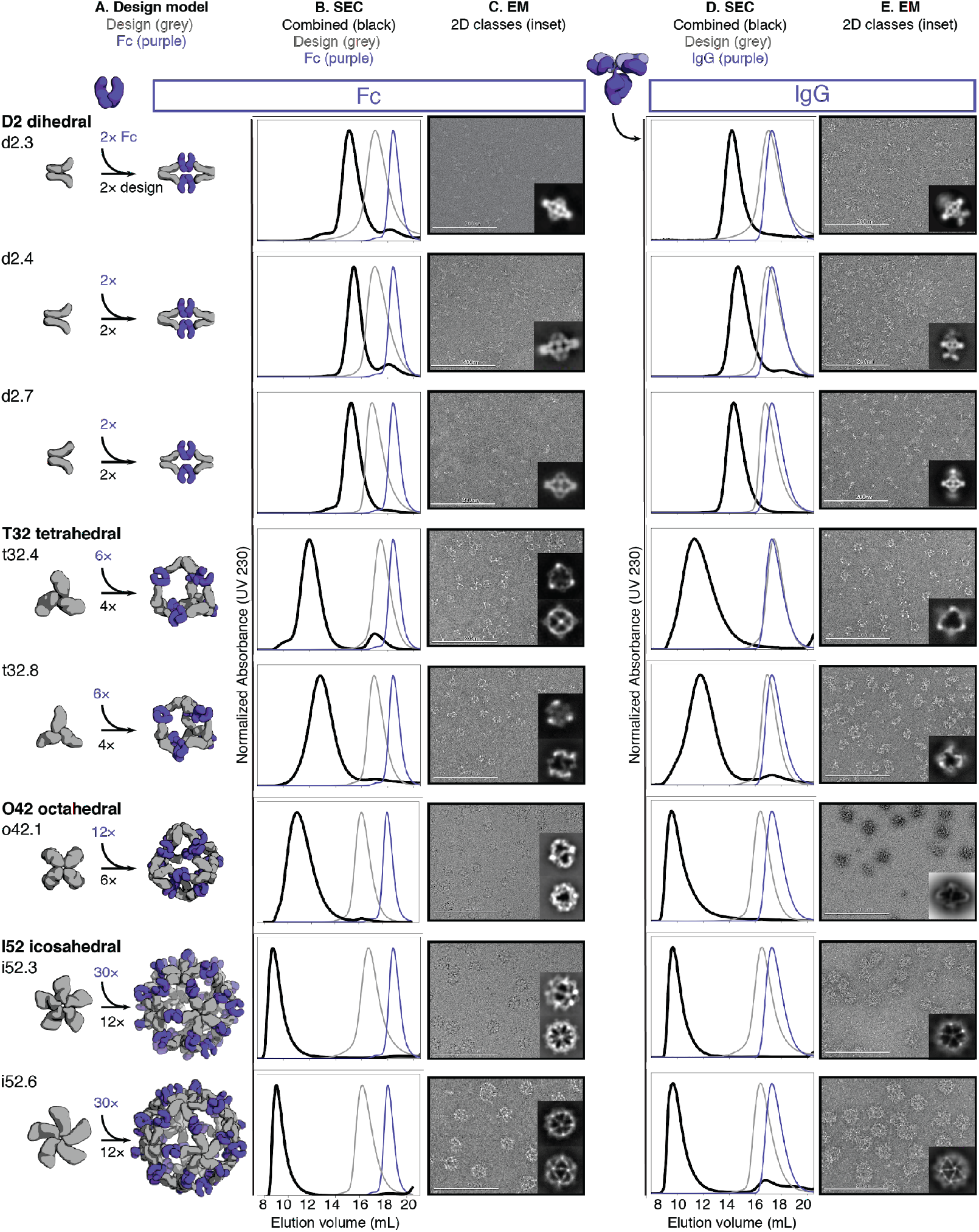
Structural characterization of AbCs. **A,** Design models, with antibody Fc (purple) and designed AbC-forming oligomers (grey). **B,** Overlay of representative SEC traces of assembly formed by mixing design and Fc (black) with those of the single components in grey (design) or purple (Fc). **C,** EM images with reference-free 2D class averages in inset; all data is from negative-stain EM with the exception of designs o42.1 and i52.3 (cryo-EM). **D-E,** SEC (**D)** and NS-EM representative micrographs with reference-free 2D class averages (**E**) of the same designed antibody cages assembled with full human IgG1 (with the 2 Fab regions intact). In all EM cases shown in **C** and **E**, assemblies were first purified via SEC, and the fractions corresponding to the left-most peak were pooled and used for imaging; this was mainly done to remove any excess of either design or Ig component.

We further characterized the eight Fc-AbCs with monodisperse SEC profiles by small-angle X-ray scattering (SAXS) and electron microscopy (EM). SAXS spectra, P(r) distributions, and radius of gyration (R_g_) values were close to design models for d2.4, d2.7, t32.4, o42.1, i52.3, and i52.6 Fc-AbCs (Fig. S3, Table S2) (*24*, *25*). The agreement to the SAXS data for the d2.3 and t32.8 design models was somewhat less good (higher R_g_ values, and deviations in the SAXS Guinier (low-q) region and P(r) distributions from those computed from the design model) potentially due to particle aggregation during data collection. Cryo-EM of o42.1 and i52.3 AbCs, and negative stain-EM (NS-EM) of the other six AbCs, showed monodisperse particle formation with individual cages, and 2D class averages resemble the design models (Fig. 2c; Table S3-S4).

AbCs also formed when assembled with full IgG antibodies (containing both Fc and Fab domains) again generating monodisperse nanocages as shown by SEC and NS-EM (Fig. 2d-e); here, the o42.1 design with IgG reproducibly elutes in the void due to the increased diameter from the added Fab domains. There is considerably more evidence of flexibility in the electron micrographs of the IgG-AbCs than the Fc-AbCs, as expected given the flexibility of the Fc-Fab hinge. In all cases, 2D class averages obtained from the NS-EM data of AbCs made with intact IgG resolved density corresponding to the Fc + design portion of the assembly (Fig. 2e).

Single-particle NS-EM and cryo-EM reconstructed 3D maps of the AbCs formed with Fc are in close agreement with the computational design models (Fig. 3). Negative-stain EM reconstructions for the dihedral (d2.3, d2.4, d2.7), tetrahedral (t32.4, t32.8), and one of the icosahedral (i52.6) nanocages clearly show dimeric “U”-shaped Fcs and longer designed protein regions that fit together as computationally designed. A single-particle cryo-EM reconstruction for the o42.1 design with Fc has clear density for the six designed tetramers sitting at the C4 vertices, which twist along the edges of the octahedral architecture to bind twelve dimeric Fcs, leaving the eight C3 faces unoccupied. The 3D density map for o42.1 with Fc suggests that the particle is flexing outwards compared to the design model, consistent with the SAXS R_g_ data. Cryo-EM density for i52.3 with Fc likewise recapitulates the 20-faced shape of a regular icosahedron, with 12 designed pentamers protruding at the C5 vertices (due to the longer length of the C5 building block compared to the monomer or Fc-binder), binding to 30 dimeric Fcs at the center of the edge, with 20 unoccupied C3 faces. Asymmetric cryoEM reconstructions of o42.1 with Fc and i52.3 with Fc had similar overall features to their respective symmetrized maps (Fig. S4).

**Figure 3.**
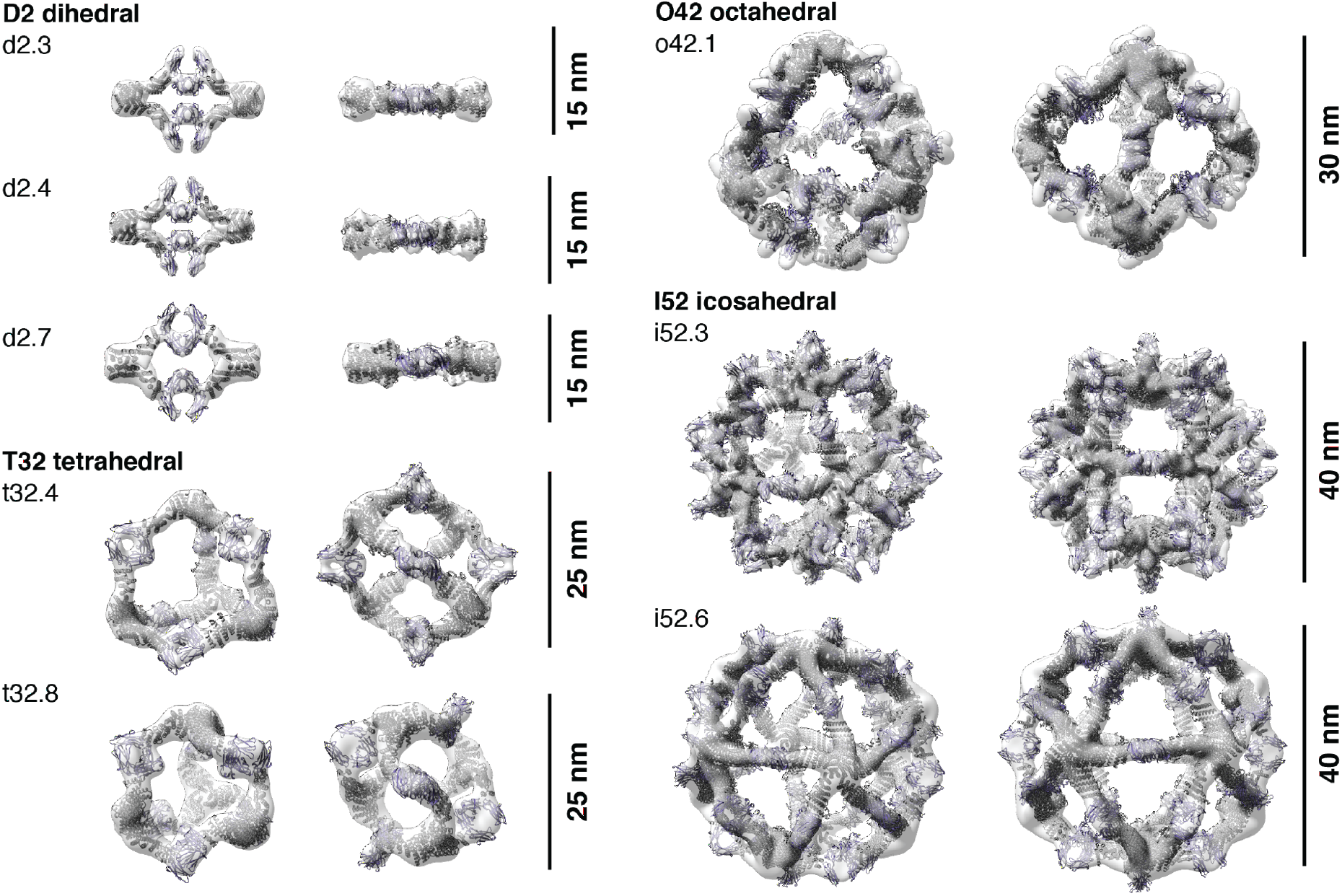
3D reconstructions of AbCs formed with Fc. Computational design models (cartoon representation) of each AbC are fit into the experimentally-determined 3D density from EM. Each nanocage is viewed along an unoccupied symmetry axis (left), and after rotation to look down one of the C2 axes of symmetry occupied by the Fc (right). 3D reconstructions from o42.1 and i52.3 are from cryo-EM analysis; all others, from NS-EM.

In a second design round, we sought to design both the homo-oligomeric building block and the nanocage in one step, and we obtained AbCs with D3 symmetry though the overall success rate was lower (Fig. S5, Table S5). SEC of assembled Fc-AbCs, SAXS analysis, and NS-EM micrographs, 2D class averages, and 3D reconstructed maps are all consistent with the shape and size of the corresponding design models.

We next assessed the stability of AbCs. Dynamic light scattering (DLS) readings remained constant over a period of 3-5 weeks for all designs when incubated at room temperature, with the exception of i52.6 which showed some broadening of the hydrodynamic radius after 2-3 weeks (Fig. S6, Table S6). Fc AbCs also did not show any degradation according to SDS gel electrophoresis. We investigated whether AbCs formed with one Fc-fusion or IgG would exchange when incubated with excess Fc; this is relevant to future in vivo applications of the AbCs where they would be in the presence of high concentration serum IgG. We formed o42.1 AbCs with GFP-Fc, then incubated these with 25-fold molar excess of RFP-Fc for up to 24 hours. SEC was used to separate the AbCs, and fluorescence readings were taken of the AbC peak fraction (Fig. S7a-c). Although the RFP-Fc signal increased slightly in the cage fraction (Fig. S7d-e, Table S7), the GFP-Fc signal was very close to that of the o42.1 GFP-Fc control, suggesting little exchange of the Fc-containing component out of the cage. Encouraged by these stability results, we moved forward with characterization of the biological impact of the AbCs.

### Enhancing cell signaling with AbCs

Our designed AbCs provide a general platform for investigating the effect of valency and geometry of receptor engagement on signaling pathway activation. Receptor dimerization, trimerization and higher order association have been implicated in transmembrane signaling in different receptor systems, but systematically probing the influence on geometry and valency on signaling has required considerable system-specific engineering (*26*). The combination of the wide range of receptor binding antibodies and natural ligands with the AbC methodology developed here in principle allow ready and systematic probing of the effect of geometry and valency of receptor subunit association on signaling for almost any pathway.

To explore the potential of this approach, we assembled antibodies and Fc-fusions targeting a variety of signaling pathways into nanocages and investigated their effects on signaling. Where possible, we attempted to use as many different cage geometries in each application. Typically, only one D2 dihedral design was used as the overall shapes of D2 dihedra were similar, and design i52.6 was avoided due to stability issues described above.

#### Induction of tumor cell apoptosis by α-DR5 nanocages

Death Receptor 5 (DR5) is a tumor necrosis factor receptor (TNFR) superfamily cell surface protein that initiates a caspase-mediated apoptotic signaling cascade terminating in cell death when cross-linked by its trimeric native ligand, TNF-related apoptosis-inducing ligand (TRAIL) (*5*, *6*, *27*–*31*). Like other members of the family, DR5 can also form alternative signaling complexes that activate non-apoptotic signaling pathways such as the NF-κB pro-inflammatory pathway and pathways promoting proliferation and migration upon ligand binding (*30*). Because DR5 is overexpressed in some tumors, multiple therapeutic candidates have been developed to activate DR5, such as α-DR5 IgG and recombinant TRAIL, but these have failed clinical trials due to low efficacy and the development of TRAIL resistance in tumor cell populations (*30*, *31*). Combining trimeric TRAIL with bivalent α-DR5 IgG leads to a much stronger apoptotic response than either component by itself, likely due to induction of larger-scale DR5 clustering via the formation of two-dimensional arrays on the cell surface (*28*).

We investigated whether α-DR5 AbCs formed with the same IgG (conatumumab) could have a similar anti-tumor effect without the formation of unbounded arrays. Five designs across four geometries were chosen (d2.4, t32.4, t32.8, o42.1, and i52.3) to represent the range of valencies and shapes (Fig. 4a). All α-DR5 AbCs were found to form single peaks on SEC with corresponding NS-EM micrographs that were consistent with the formation of assembled particles (Fig. 2d-e). All α-DR5 AbCs caused caspase 3/7-mediated apoptosis at similar levels to TRAIL in a colorectal tumor cell line (Colo205), whereas the antibody alone or AbCs formed with bare Fc did not lead to caspase-3/7 activity or cell death, even at the highest concentrations tested (comparing molarity at an antibody to antibody level; Fig. S8a, Table S8). On the TRAIL-resistant renal cell carcinoma line RCC4, we found that α-DR5 AbCs induced caspase-8 and caspase-3,7 activity (Fig. 4b, Fig. S8b-c) and designs t32.4, t32.8, and o42.1 greatly reduced cell viability at 150 nM concentration (Fig. 4c). Free α-DR5 antibody, Fc-only AbCs, or TRAIL neither activated caspase nor decreased cell viability after four days (Fig. 4b-d, Fig. S8b-d). Because designs t32.4 and o42.1 activated caspase-3,7 at 100-fold lower concentrations (1.5 nM, Fig. 4b), we tested prolonged 6 day treatment of these at 150 nM with RCC4 cells, which resulted in the killing of nearly all cells after six days, suggesting that RCC4 cells do not acquire resistance to the nanocages (Fig. 4e). We next investigated the downstream pathways activated by the α-DR5 AbCs by analyzing their effects on cleaved PARP, a measure of apoptotic activity. Consistent with the caspase and cell viability data, α-DR5 AbCs increased cleaved PARP in RCC4 cells, while free α-DR5 antibody, TRAIL or o42.1 Fc AbCs did not result in an increase in cleaved PARP over baseline (Fig. 4e-f shows o42.1 as a representative example; Fig. S8c,e). The α-DR5 AbCs did not significantly induce apoptosis in healthy primary kidney tubular cells (Fig. S8f-g).

**Figure 4.**
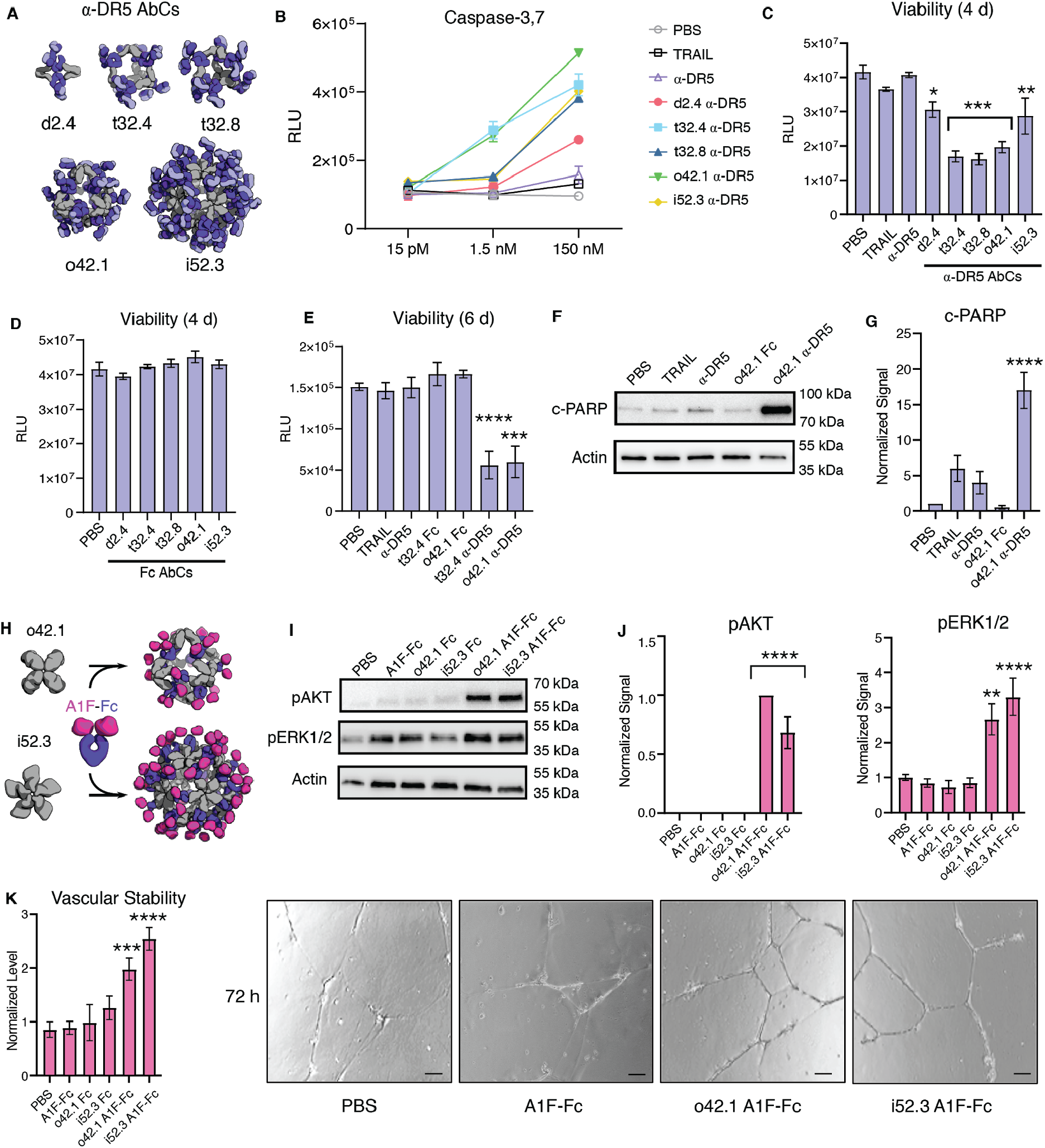
AbCs activate apoptosis and angiogenesis signaling pathways. Caspase-3,7 is activated by AbCs formed with α-DR5 antibody (**A**), but not the free antibody, in RCC4 renal cancer cells (**B**). **C-D**, α-DR5 AbCs (**C**), but not Fc AbC controls (**D**) reduce cell viability 4 days after treatment. **E**, α-DR5 AbCs reduce viability 6 days after treatment. **F-G**, o42.1 α-DR5 AbCs enhance PARP cleavage, a marker of apoptotic signaling; **G**, quantification of **F** relative to PBS control. **H**, The F-domain from Angiopoietin-1 was fused to Fc (A1F-Fc) and assembled into octahedral (o42.1) and icosahedral (i52.3) AbCs. **I**, Representative Western blots show that A1F-Fc AbCs, but not controls, increase pAKT and pERK1/2 signals. **J**, quantification of **I**: pAKT quantification is normalized to o42.1 A1F-Fc signaling (no pAKT signal in the PBS control); pERK1/2 is normalized to PBS. **K**, A1F-Fc AbCs increase vascular stability after 72 hours. Left: quantification of vascular stability compared to PBS. Right: representative images. All error bars represent means ± SEM; means were compared using ANOVA and Dunnett post-hoc tests (Tables S8, S9).

#### Tie-2 pathway activation by Fc-Angiopoietin 1 nanocages

Certain receptor tyrosine kinases (RTKs), such as the Angiopoietin-1 receptor (Tie2), activate downstream signaling cascades when clustered (*32*, *33*). Scaffolding the F-domain from angiopoietin-1 (A1F) onto nanoparticles induces phosphorylation of AKT and ERK1/2, enhances cell migration and tube formation *in vitro*, and improves wound healing after injury *in vivo (33)*. Therapeutics with these activities could be useful in treating conditions characterized by cell death and inflammation, such as sepsis and acute respiratory distress syndrome (ARDS). To determine whether the AbC platform could be used to generate such agonists, we assembled o42.1 and i52.3 AbCs with Fc fusions to A1F (Fig. 4g, Fig. S9a-b). The octahedral and icosahedral A1F-AbCs, but not Fc-only controls or free Fc-Ang1F, significantly increased AKT and ERK1/2 phosphorylation above baseline (Fig. 4h-i) and enhanced vascular stability (Fig. 4j, Fig. S9c-d, Table S9), comparable to a A1F-presenting octamer (Fig. S9c) (*33*). To further address particle stability upon AbC formation, o42.1 A1F-Fc AbCs were incubated for 24 hours with 100% human serum at 4°C or 37°C. If AbCs rapidly fell apart or exchanged with serum antibodies, we would likely see a decrease in signal; nonetheless, o42.1 A1F-Fc AbCs in serum remained equally active as control o42.1 A1F-Fc AbCs (Fig. S9e).

#### α-CD40 nanocages activate B cells

CD40, a TNFR superfamily member expressed on antigen presenting dendritic cells and B cells, is cross-linked by trimeric CD40 ligand (CD40L or CD154) on T cells, leading to signaling and cell proliferation (*34*, *35*). Non-agonistic α-CD40 antibodies can be converted to agonists by adding cross-linkers such as FcγRIIb-expressing Chinese Hamster Ovary (CHO) cells (*34*). We investigated whether assembling a non-agonist α-CD40 antibody (LOB7/6) into nanocages could substitute for the need for cell surface presentation; we focused on the o42.1 design given its promising data in the DR5 and A1F experiments. Octahedral AbCs were assembled with α-CD40 LOB7/6 IgG (Fig. 5a); SEC and dynamic light scattering (DLS) characterization showed these to be monodisperse and of the expected size (Fig. S10a-b). The octahedral α-CD40 LOB7/6 AbCs were found to induce robust CD40 activation in CD40-expressing reporter CHO cells (J215A, Promega), at concentrations around 20-fold less than a control activating α-CD40 antibody (Promega), while no activation was observed for the free LOB7/6 antibody or octahedral AbC formed with non-CD40 binding IgG (Fig. 5b, Table S10). Thus nanocage assembly converts the non-agonist α-CD40 mAb into a CD40 pathway agonist.

**Figure 5.**
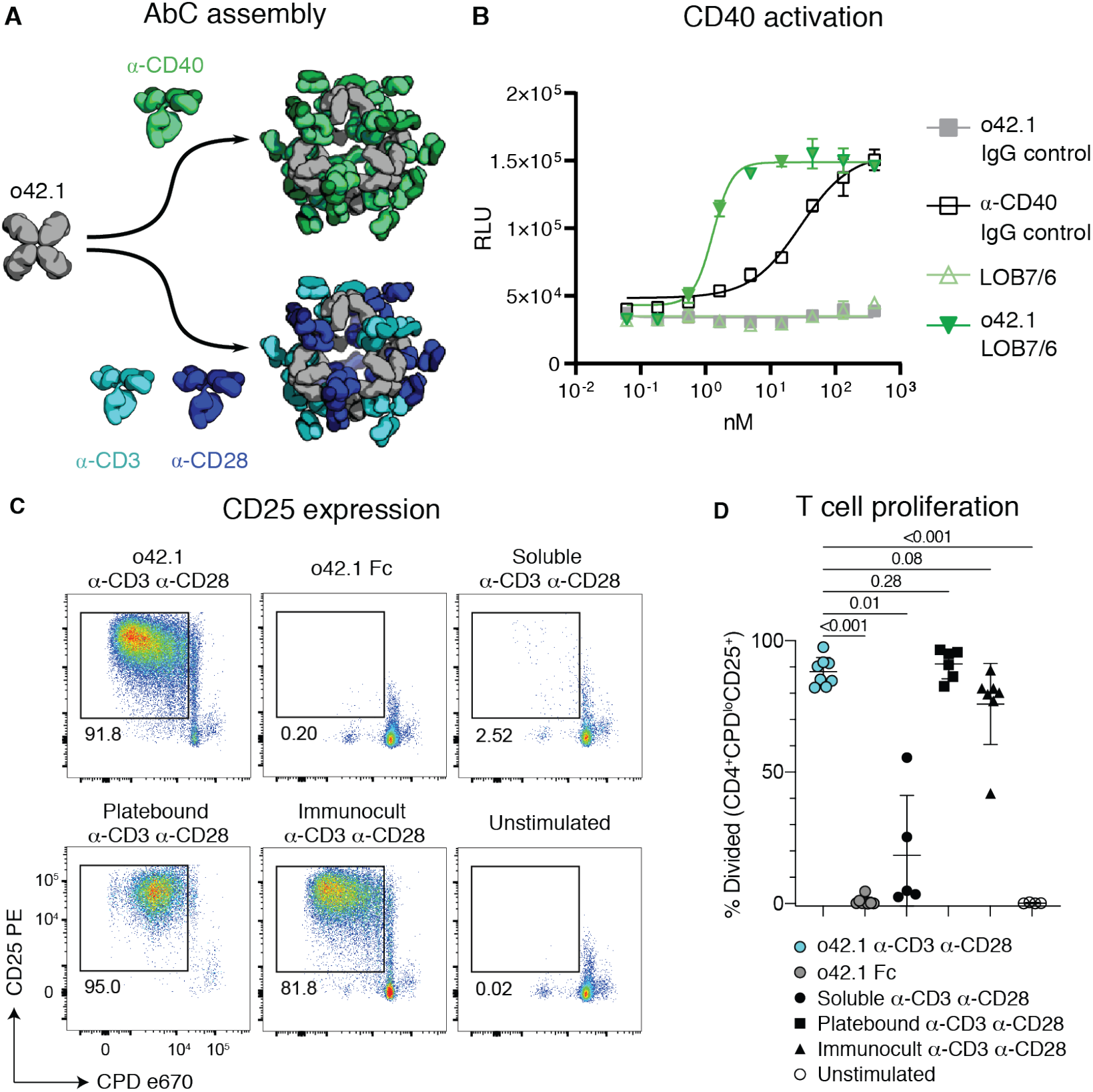
Activation of immune cells by α-CD40 and α-CD3/28 AbCs. **A,** Octahedral AbCs are produced with either α-CD40 or pre-mixed α-CD3 and α-CD28 antibodies. **B,** CD40 pathways are activated by α-CD40 LOB7/6 octahedral nanocages but not by free α-CD40 LOB7/6 or controls. Scale bars represent means ± SD, n=3; EC50s reported in Table S10. **C-D,** T cell proliferation and activation is strongly induced by α-CD3 α-CD28 mosaic AbCs compared to unassembled (soluble) α-CD3 α-CD28 antibodies. Representative plots (**C**) show the frequency of dividing, activated cells (CPD^lo^CD25^+^). Mosaic AbC-induced proliferation is comparable to traditional positive controls, platebound or tetrameric α-CD3 α-CD28 antibody bead complex (Immunocult). Gated on live CD4^+^ T cells. Summary graph (**D**) shows mean ± SD. Significance was determined by Kruskal-Wallis tests correcting for multiple comparisons using FDR two-stage method (n=4-8 per condition). Adjusted p values are reported. CPD, Cell Proliferation Dye.

#### α-CD3/28 mosaic nanocages cause T cell proliferation

T cell engineering technologies such as chimeric antigen receptor (CAR)-T cell therapy require the *ex vivo* expansion and activation of T cells, often carried out by presenting CD3- and CD28-binding ligands on the surface of beads or a plate (*36*, *37*). We sought to eliminate the need for the solid support for T cell activation by using AbCs formed with both α-CD3 and α-CD28 antibodies. Equimolar amounts of α-CD3 and α- CD28 were pre-mixed and then incubated with the o42.1 design to form “mosaic” octahedral cages (Fig. 5a, Fig. S10c-d). Octahedral α-CD3/CD28 AbC, but not free antibody or Fc nanocage, led to proliferation of naïve T cells sorted from healthy donor peripheral blood mononuclear cells (PBMCs) as read out by expression of the T cell activation marker CD25 (Fig. 5c) and proliferation assays (Fig. 5d); activation levels were similar to tetrameric or plate-bound α-CD3/CD28 stimulation controls. Together with the α-CD40 activation, these results demonstrate how readily specific immune cell pathways can be activated by simply swapping in different antibodies into the cage architecture.

### Enhancing viral neutralization with AbCs

There is considerable current effort directed at development of antibodies targeting the SARS-CoV-2 spike (S) protein for prophylaxis and post-exposure therapy for the current COVID-19 pandemic (*13*, *38*–*43*). We hypothesized that assembling α-SARS-CoV-2 antibodies into nanocages could potentially increase their neutralization potency by increasing avidity for viral particles, as multivalency was recently found to increase SARS-CoV-2 neutralization using apoferritin to scaffold binding domains (*13*). Octahedral AbCs (o42.1) formed with the SARS-CoV-2 S-binding antibodies CV1 or CV30 (*38*) were more effective at neutralizing pseudovirus entry into angiotensin-converting enzyme 2 (ACE2)-expressing cells than free CV1 or CV30, dropping the apparent IC_50_ over 200-fold and around 2.5-fold, respectively (comparing molarity at an antibody to antibody level; Fig. 6a-c, Fig. S11a-b, Table S11). The potency of a third non-neutralizing antibody, CV3, was unchanged by assembly into the nanocage format (Fig. S11c). We found that assembly into octahedral AbCs of Fc-ACE2, which directly engages the receptor binding domain of the spike protein (*43*), enhanced neutralization around 7-fold compared to free Fc-ACE2 fusion for SARS-CoV-2 pseudovirus and 2.5-fold for SARS-CoV-1 pseudovirus (Fig. 6d, Fig. S11d-e, Table S11).

**Figure 6.**
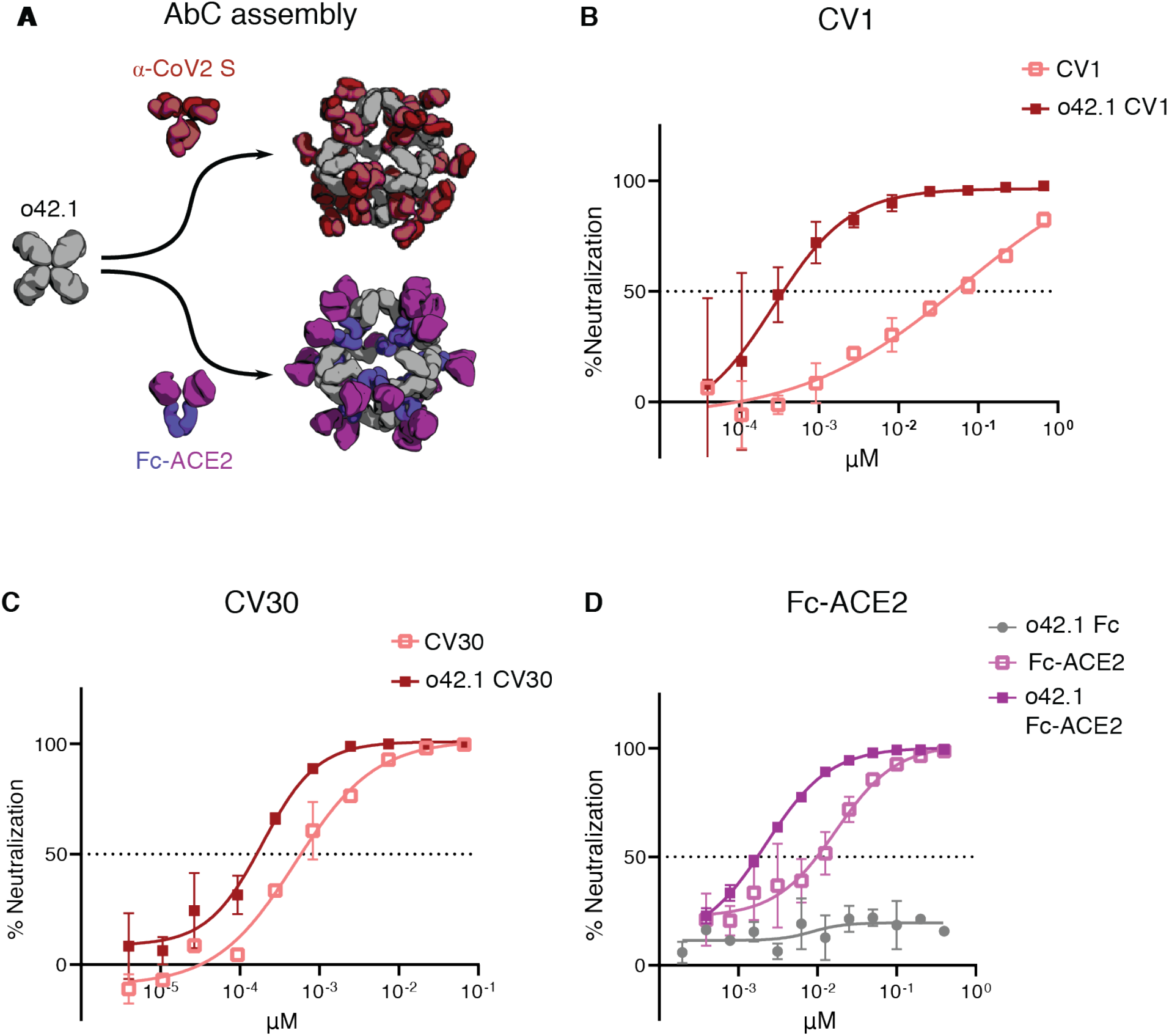
Nanocage assembly enhances SARS-CoV-2 pseudovirus neutralization. **A,** Octahedral AbCs are produced with either α-CoV-2 S IgGs or Fc-ACE2 fusion. **B-C,** SARS-CoV-2 S pseudovirus neutralization by octahedral AbC formed with α-CoV-2 S IgGs CV1 (**B)** or CV30 (**C**) compared to un-caged IgG. **D,** SARS-CoV-2 S pseudovirus neutralization by Fc-ACE2 octahedral AbC compared to un-caged Fc-ACE2. Error bars represent means ± SD, n=2; IC50s reported in Table S11.

## Discussion

Our approach goes beyond previous computational design efforts to create functional nanomaterials by integrating form and function; whereas previous work has fused functional domains onto assemblies constructed from separate structural components (*2*–*12*, *27*), our AbCs employ antibodies as both structural and functional components. By fashioning designed antibody-binding, cage-forming oligomers through rigid helical fusion, a wide range of geometries and orientations can be achieved. This design strategy can be generalized to incorporate other homo-oligomers of interest into cage-like architectures. For example, nanocages could be assembled with viral glycoprotein antigens using components terminating in helical antigen-binding proteins, or from symmetric enzymes with exposed helices available for fusion to maximize the proximity of active sites working on successive reactions. The AbCs offer considerable advantages in modularity compared to previous fusion of functional domain approaches; any of the thousands of known antibodies can be used “off-the-shelf” to form multivalent cages by mixing with the appropriate design to form the desired symmetric assembly, provided sufficient protein A/Fc affinity. EM and SEC demonstrate monodispersity comparable to IgM and control over binding domain valency and positioning that is not (to our knowledge) attained by other antibody-protein nanoparticle formulations (*44*).

AbCs show considerable promise as signaling pathway agonists. Assembly of antibodies against RTK- and TNFR-family cell-surface receptors into AbCs led to activation of diverse downstream signaling pathways involved in cell death, proliferation, and differentiation. While antibody-mediated clustering has been previously found to activate signaling pathways (*7*, *28*, *34*), our approach has the advantage of much higher structural homogeneity, allowing more precise tuning of phenotypic effects and more controlled formulation. Two or more different receptor engaging antibodies or Fc-fusions can be readily incorporated into the same cage by simple mixing, allowing exploration of the effects on downstream signaling of bringing together different receptors and comodulators in different valencies and geometries. There are exciting applications to targeted delivery, as the icosahedral AbCs have substantial internal volume (around 15,000 nm^3^, based on an estimated interior radius of 15.5 nm) that could be used to package nucleic acid or protein cargo, and achieving different target specificity in principle is as simple as swapping one antibody for another. An important next step towards the possibility of augmenting antibody therapeutics with our designed AbCs-forming oligomers will be investigating the pharmacokinetic and biodistribution properties of these molecules, their immunogenicity, and whether the Fc domains can still activate effector functions. We anticipate that the AbCs developed here, coupled with the very large repertoire of existing antibodies, will be broadly useful across a wide range of applications in biology and medicine.

## Supporting information

Supplement Antibody cages

## Acknowledgements

We thank Yang Hsia and Rubul Mout for help with computational design, Brooke Fiala, Natalie Brunette, Elizabeth Kepl, Michelle DeWitt, Lauren Carter, Michael Murphy, and Colin Correnti for providing protein materials used at various stages of the project. We thank Alexis Courbet for his help collecting micrographs for D3-08 and D3-36 designs, and Joel Quipse for his help in maintaining and operating the electron microscopes used. We thank Olivier Poncelet for technical help. We thank Dr. Akilesh for providing primary kidney tubular epithelial cells. We would like to thank Amgen for providing the AMG-655 antibody used herein. The SAXS work was conducted at the Advanced Light Source (ALS), a national user facility operated by Lawrence Berkeley National Laboratory on behalf of the Department of Energy, Office of Basic Energy Sciences, through DOE BER IDAT grant (DE-AC02-05CH11231) and NIGMS supported ALS-ENABLE (GM124169-01) and National Institute of Health project MINOS (R01GM105404). We thank the staff at the Advanced Light Source SIBYLS beamline at Lawrence Berkeley National Laboratory, including K. Burnett, G. Hura, M. Hammel, J. Tanamachi, and J. Tainer for the services provided through the mail-in SAXS program, which is supported by the DOE Office of Biological and Environmental Research Integrated Diffraction Analysis program DOE BER IDAT grant (DE-AC02-05CH11231) and NIGMS supported ALS-ENABLE (GM124169-01) and National Institute of Health project MINOS (R01GM105404).

This work was supported by NSF grant CHE 1629214 (N.P.K., D.B.), HHMI (W.S., D.B), the Audacious Project at the Institute for Protein Design (R.D., I.V., M.R.T., A.R.), The Washington Research Foundation (F.S., J.L.), The Nordstrom-Barrier Directors Fund at the Institute for Protein Design (A.E.), Washington State General Operating Fund for the Institute for Protein Design (L. Stewart), Wu Tsai Translational Investigator Fund (G.U.), and Nan Fung Life Sciences Translational Investigator Fund (J.A.F.). This work was supported by generous donations to Fred Hutch COVID-19 Research Fund (M.F.J., L.J.H., Y.W., A.T.M., L. Stamatatos). This work was also supported by NIH grant R01AI127726 (P.A.M., M.L.F., and D.J.C). This research was funded in part through the NIH/NCI Cancer Center Support Grant P30 CA015704 and ISCRM Pilot Award (J.M.). This work was also supported by the NIAID / NIH (DP1AI158186 and HHSN272201700059C to D.V.) NIGMS/NIH (R01GM120553 to D.V.), a Pew Biomedical Scholars Award to D.V., and a Burroughs Wellcome Investigators in the Pathogenesis of Infectious Diseases award to D.V., Fast Grants (D.V.), the University of Washington Arnold and Mabel Beckman cryoEM center.

## Author contributions

R.D., H.V.D., G.U., J.A.F., J.M., D.V., and D.B. designed the research. R.D. designed the AbC forming oligomers with contributions from I.V. and D.B. for d2.3, d2.4, d2.7, and t32.4. I.V., W.S., and D.B. wrote the computational fusion protocol used during design, based on I.V.’s original concept. R.D., H.V.D., G.U., J.A.F., and A.R. produced and characterized all designs. F.S. designed and characterized the Fc-binding repeat protein. R.D. and A.E. prepared samples for SAXS analysis, and R.D., G.U, J.A.F., and A.E. analyzed the SAXS results. R.D. and H.V.D. prepared the NS-EM grids. H.V.D, R.D. and A.E. collected NS-EM data. R.D. analyzed the NS-EM datasets. H.V.D. prepared samples and analyzed the cryo-EM datasets for o42.1 and i52.3 nanocages with Fc. A.E., G.U., J.A.F., M.M., and R.D. designed and characterized D3-08 and D3-36. R.D. and J.L. designed and conducted the stability and exchange experiments. G.U., J.A.F., S.S., J.L., H.R., and J.M. designed the DR5 experiments, with S.S. performing the assays. G.U., J.A.F., Y.T.Z., I.X.R., J.M., and H.R. designed the A1F-Fc experiments, with Y.T.Z. and I.X.R. performing the assays. M.R.T., R.D., N.P.K., and D.B. designed the CD40 agonist experiments, with M.R.T. conducting the cell culture assays. P.A.M., M.L.F., and D.J.C. designed and conducted the T cell proliferation assays. M.F.J., L.J.H., Y.W., A.T.M., and L.S. (L. Stamatatos) designed the pseudovirus neutralization studies with IgGs. H.V.D., A.C.W., L.S. (L. Stewart), D.V., and D.B. designed the pseudovirus neutralization studies with Fc-ACE2.

