## Supplement Antibody cages for "Designed proteins assemble antibodies into modular nanocages"

**Affiliations:**

**This document includes:**

1. Materials and methods
2. Figures S1-S11
3. Tables S1-S15
4. Supplementary text
5. Full reference list (*1-62*)

### Materials and Methods

#### Computational design and testing of Fc-binder helical repeat protein (DHR79-FcB)

The crystal structure of the B-domain from *S. aureus* protein A in complex with Fc fragment (PDB ID: 1L6X) was relaxed with structure factors using Phenix Rosetta (*45*, *46*). Briefly, the RosettaScripts MotifGraft mover was used to assess suitable solutions to insertions of the protein A binding motif extracted from 1L6X into a previously reported designed helical repeat protein (DHR79) (*19*). Specifically, a minimal protein A binding motif was manually defined and extracted and used as a template for full backbone alignment of DHR79 while retaining user-specified hotspot residues that interact with the Fc domain in the crystal structure at the Fc/DHR interface and retaining native DHR residues in all other positions. The MotifGraft alignment was followed by 5 iterations of FastDesign and 5 iterations of FastRelax in which the DHR side chain and backbone rotamers were allowed to move while the Fc context was completely fixed. The best designs were selected based on a list of heuristic filter values. Fig. S1a shows the design model of DHR79-FcB.

Designs were initially assessed via yeast surface display binding to biotinylated Fc protein; yeast display procedures followed previously-published protocols (*47*). Upon confirmation of a qualitative binding signal, the design was cloned into a pET29b expression vector with a C-terminal His-tag. The protein was expressed in BL21 DE3 in autoinduction medium (6 g tryptone, 12 g yeast extract, 10 mL 50×M, 10 mL 50x5052, 1 mL 1M $MgSO_4$, 100 μL Studier Trace metals, 50 μg kanamycin antibiotic, brought to a final volume of 500 mL using filtered water) for 20 hours at 27°C at 225 rpm; 50×M, 50×5052, and Studier trace metals were prepared according to previously-published recipes (*48*). Cells were resuspended in lysis buffer (20mM Tris, 300mM NaCl, 30mM imidazole, 1mM phenylmethylsulfonyl fluoride (PMSF), 5% glycerol (v/v), pH 8.0) and lysed using a microfluidizer at 18000 PSI. Soluble fractions were separated via centrifugation at 24,000×g. IMAC with Ni-NTA batch resin was used for initial purification; briefly, nickel-nitrilotriacetic acid (Ni-NTA) resin was equilibrated with binding buffer (20mM Tris, 300mM NaCl, 30mM imidazole, pH 8.0), soluble lysate was poured over the columns, columns were washed with 20 column volumes (CVs) of binding buffer, and eluted with 5 CVs of elution buffer (20mM Tris, 300mM NaCl, 500mM imidazole, pH 8.0). Size exclusion chromatography (SEC) with a Superdex 200 column was used as the polishing step (Fig. S1b). SEC buffer was 20 mM Tris/HCl pH 7.4, 150 mM NaCl.

Affinity of DHR79-FcB to biotinylated IgG1 and biotinylated Fc protein bound to streptavidin plates was assessed using Octet Biolayer Interferometry (BLI). Data was fit using a 1:1 binding mode. Both 1:1 and 2:1 binding stoichiometries were assessed, and it was determined that the 1:1 binding mode better accounted for the noise in the experiment. This was tested on the Fc binder monomer before any Fc binder-oligomer fusions were made. DHR79-FcB exhibits a 71.7 nM affinity to IgG1 (full antibody) and a 113 nM affinity to the IgG1 Fc protein (Fig S1c).

**Computational Design of Antibody Nanocages**

Input .pdb files were compiled to use as building blocks for the generation of antibody cages. For the protein A binder model, the Domain D from *Staphylococcus aureus* Protein A (PDB ID 1DEE) was aligned to the B-domain of protein A bound to Fc (PDB ID: 1L6X) (*18*, *45*). The other Fc-binding design structure, where protein A was grafted onto a helical repeat protein, was also modeled with Fc from 1L6X. PDB file models for monomeric helical repeat protein linkers (42) and cyclic oligomers (2 C2s, 3 C3s, 1 C4, and 2 C5s) that had at least been validated via SAXS were compiled from previous work from our lab (*19*–*21*). Building block models were manually inspected to determine which amino acids were suitable for making fusions without disrupting existing protein-protein interfaces.

These building blocks were used as inputs, along with the specified geometry and fusion orientation, into the alpha helical fusion software ("WORMS," ran using instructions provided at https://github.com/willsheffler/worms; also see Supplementary Text for a description on how to operate WORMS) (*14*, *15*). Fusions were made by overlapping helical segments at all possible allowed amino acid sites. Fusions are then evaluated for deviation for which the cyclic symmetry axes intersect according to the geometric criteria: D2, T32, O32, O42, I32, and I52 intersection angles are 45.0°, 54.7°, 35.3°, 45.0°, 20.9°, and 31.7°, respectively (*22*) with angular and distance tolerances of at most 5.7° and 0.5 Å respectively. Post-fusion .pdb files were manually filtered to ensure that the N-termini of the Fc domains are facing outwards from the cage, so that the Fabs of an IgG would be external to the cage surface. Sequence design was performed using Rosetta symmetric sequence design (SymPackRotamersMover in RosettaScripts) on residues at and around the fusion junctions (*49*), with a focus on maintaining as many of the native residues as possible. Residues were redesigned if they clashed with other residues, or if their chemical environment was changed after fusion (e.g. previously-core facing residues were now solvent-exposed). Index residue selectors were used to prevent design at Fc residue positions. See Supplemental Materials 4 for an example .xml file used in post-fusion design.

**Protein expression for AbC-forming designs and Fc constructs**

*Bacterial expression of AbC-forming designs*

Genes were codon optimized for bacterial expression of each designed AbC forming oligomer, with a C-terminal glycine/serine linker and 6× C-terminal histidine tag appended. Synthetic genes were cloned into pet29b+ vectors between NdeI and XhoI restriction sites; the plasmid contains a kanamycin-resistant gene and T7 promoter for protein expression. Plasmids were transformed into chemically competent Lemo21(DE3) *E. coli* bacteria using a 15-second heat shock procedure as described by the manufacturer (New England Biolabs). Transformed cells were added to auto-induction expression media, as described above, and incubated for 16 hours at 37°C and 200 rpm shaking (*48*). Cells were pelleted by centrifugation at 4000×g and resuspended in lysis buffer (150 mM NaCl, 25 mM Tris-HCl, pH 8.0, added protease inhibitor and DNAse). Sonication was used to lyse the cells at 85% amplitude, with 15 second on/off cycles for a total of 2 minutes of sonication time. Soluble material was separated by centrifugation at 16000×g. IMAC was used to separate out the His-tagged protein in the soluble fraction as described above. IMAC elutions were concentrated to approximately 1 mL using 10K MWCO spin concentrators, filtered through a 0.22 uM spin filter, and run over SEC as a final polishing step (SEC running buffer: 150 mM NaCl, 25 mM Tris-HCl, pH 8.0).

*Production of Fc and Fc-fusions*

Synthetic genes were optimized for mammalian expression and subcloned into the CMV/R vector (VRC 8400) (*50*). XbaI and AvrII restriction sites were used for insertion of the target gene (Fc, GFP-Fc, RFP-Fc, or A1F-Fc). Gene synthesis and cloning was performed by Genscript. Expi293F cells were grown in suspension using Expi293 Expression Medium (Thermo Fisher Scientific) at 150 RPM, 5% $CO_2$, 70% humidity, 37°C. At confluency of ~$2.5\times10^6$ cells/mL, the cells were transfected with the vector encoding the Fc or Fc-fusion (1000 µg per 1 L of cells) using PEI MAX (Polysciences) as a transfection reagent. Cells were incubated for 96 hours, after which they were spun down by centrifugation (4,000xg, 10 min, 4 °C) and the protein-containing supernatant was further clarified by vacuum-filtration (0.45 µm, Millipore Sigma). In preparation of nickel-affinity chromatography steps, 50 mM Tris, 350 mM NaCl, pH 8.0 was added to the clarified supernatant. For each liter of supernatant, 4 mL of Ni Sepharose excel resin (GE) was added to the supernatant, followed by overnight shaking at 4 °C. After 16-24 h, resin was collected and separated from the mixture and washed twice with 50 mM Tris, 500 mM NaCl, 30 mM imidazole, pH 8.0 prior to elution of desired protein with 50 mM Tris, 500 mM NaCl, 300 mM imidazole, pH 8.0. Eluates were purified by SEC using a Superdex 200 Increase column.

**Structural evaluation of AbC-forming designs**

Designs that produced monodisperse SEC peaks around their expected retention volume were combined with Fc from human IgG1. Cage components were incubated at 4°C for at minimum 30 minutes. 100 mM L-arginine was added during the assembly to AbCs formed with the i52.6 design, as this was observed to maximize the formation of the designed AbC I52 and prevent the formation of visible “crashed out” aggregates (*23*). Fc-binding and cage formation were confirmed via SEC; earlier shifts in retention time (compared to either component run alone) show the formation of a larger structure. NS-EM was used as described below to confirm the structures of designs that passed these steps.

For confirming AbC structures with intact IgGs, human IgG1 (hIgG1) was combined with AbC-forming designs following the same protocol for making Fc cages. This assembly procedure was also followed for all IgG or Fc-fusion AbCs reported hereafter. The data in Figure 2d-e shows AbCs formed with the $\alpha$-DR5 antibody AMG-655 (*23*) for the following designs: d2.3, d2.4, d2.7, t32.4, t32.8, o42.1, and i52.3. The data for the i52.6 design shown in Figure 2d-e is from AbCs formed with the hIgG1 antibody mpe8 (*51*); this was simply due to limited AMG-655 availability at the time of the experiment and not a reflection on i52.6/AMG-655 assembly competency. Tables S12 and S13 show the list of IgGs and Fc fusions that have been formed into AbCs. Table S14 lists the amino acid sequences of all successful AbC-forming designs; Table

S15 lists the amino acid sequences of Fc and Fc-fusions used in the following experiments.

*SAXS*

Samples were prepared for small-angle X-ray scattering (SAXS) analysis by first expressing and purifying AbCs with Fc as described above. Fc AbCs were sized into 150 mM NaCl and 25 mM Tris-HCl at pH 8.0. Fractions corresponding to the Fc AbC peak off SEC were combined and glycerol was added at 2% final concentration. Samples were concentrated to between 1-3 mg/mL using a 10K molecular weight cut-off (MWCO) benchtop spin concentrator. The flow-through was used as a blank for buffer subtraction during SAXS analysis. Proteins were then passed through a 0.22 μm syringe filter (Millipore). These proteins and buffer blanks were shipped to the SIBYLS High Throughput SAXS Advanced Light Source in Berkeley, California to obtain scattering data (*25*). Scattering traces were fit to theoretical models using the FOXS server (https://modbase.compbio.ucsf.edu/foxs/) (*24*).

*NS-EM specimen preparation and data collection of Fc and IgG AbCs*

For all samples except o42.1 Fc and i52.3 Fc, 3.0 μL of each SEC-purified sample between 0.008- 0.014 mg/mL in TBS pH 8.0 was applied onto a 400-mesh or 200-mesh Cu grid glow-discharged carbon-coated copper grids for 20s, followed by 2× application of 3.0 μL 2% nano-W or UF stain.

For 14 samples (d2.3 Fc, d2.4 Fc, d2.7 Fc, t32.4 Fc, t32.8 Fc, i52.6 Fc,  d2.3 Fc, d2.4 IgG, d2.7 IgG, t32.4 IgG, t32.8 IgG, o42.1 IgG, i52.3 IgG and i52.6 IgG samples), micrographs were recorded using Leginon software (*52*) on a 120 kV FEI Tecnai G2 Spirit with a Gatan Ultrascan 4000 4k × 4k CCD camera at 67,000 nominal magnification (pixel size 1.6 Å/pixel) or 52,000 nominal magnification (pixel size 2.07 Å) at a defocus range of 1.0 – 2.5 μm (Table S3).

For D3-08 Fc and D3-36 Fc samples, micrographs were recorded via manual acquisition on a 120 kV FEI L120C Talos TEM with a 4K × 4K Gatan OneView camera at 57,000 nominal magnification (pixel size 2.516 Å/pixel) at a defocus range of 1.0 – 2.5 μm.

*NS-EM data analysis of Fc and IgG AbCs*

Particles were picked either with DoGPicker within the Appion interface (*53*) or cisTEM (*54*); both are reference-free pickers. Contrast-transfer function (CTF) was estimated using GCTF (*55*) or cisTEM. 2D class averages were generated in cryoSPARC (*56*) or in cisTEM. Reference-free *ab initio* 3D reconstruction using particles selected from 2D class averages from each dataset was performed in cryoSPARC or in cisTEM (Table S4).

*Cryo-EM specimen preparation and data collection of o42.1 and i52.3 AbCs*

3.0 μL of o42.1 Fc sample at 0.8 mg/mL in TBS pH 8.0 with 100mM Arginine was applied onto glow-discharged 1.2μm C-flat copper grids. 3.0 μL of i52.3 Fc sample at 0.1 mg/mL in TBS pH 8.0 was applied onto glow-discharged 1.2 μm C-flat copper grids coated with a thin layer of continuous homemade carbon. Grids were then plunge-frozen in liquid ethane, cooled with liquid nitrogen using an FEI MK4 Vitrobot with a 6 second blotting time and 0 force for o42.1 Fc, and 2.5 second blotting time and -1 force for i52.3 Fc. The blotting process took place inside the Vitrobot chamber at 20°C and 100% humidity. Data acquisition was performed with the Leginon data collection software on an FEI Glacios electron microscope at 200 kV and a Gatan K2 Summit camera. The nominal magnification was 36,000 with a pixel size of 1.16 Å/pixel. The dose rate was adjusted to 8 counts/pixel/s. Each movie was acquired in counting mode fractionated in 50 frames of 200 ms/frame.

*Cryo-EM data analysis of o42.1 and i52.3 AbCs*

For both o42.1 Fc and i52.3 Fc datasets, micrographs were motion-corrected using Warp (*57*) and exported to cryoSPARC for CTF estimation with CTFFIND4. A manually picked set of particles was used to generate 2D class averages that were subsequently used (after low-pass filtering to 20 Å resolution) for Template Picker in cryoSPARC on the whole dataset. Particles were extracted with a box size of 648 pixels and subjected to reference-free 2D classification in cryoSPARC.

For the o421.1Fc dataset, particles from selected 2D classes were classified using *ab initio* reconstruction in cryoSPARC with default parameters, 4 classes, and no symmetry imposed. Micrographs containing particles from 2 classes out of 4 resulting *ab initio* classes were subjected to Manually Curate Exposures function in cryoSPARC to remove bad micrographs. This set of particles after manual curation in cryoSPARC underwent another round of *ab initio* reconstruction in cryoSPARC with default parameters, 4 classes, and no symmetry imposed. One class (4032 particles) from these 4 resulting *ab initio* classes was selected for Non-uniform refinement (NUR) in cryoSPARC with no symmetry applied or with O symmetry applied. The NUR map with no symmetry has a 17.7 Å resolution and the NUR map with O symmetry applied has a 11.14 Å resolution; both maps were similar, justifying imposing O symmetry for the final reconstruction.

For the i52.3 Fc dataset, after 1 round of 2D classification in cryoSPARC, the micrographs containing particles in a set of selected 2D classes were subjected to Manually Curate Exposures function in cryoSPARC to remove bad micrographs. This set of particles after manual curation in cryoSPARC were subjected to another round of 2D classification in cryoSPARC. 3,918 particles from selected 2D classes were reconstructed into one 3D class using *ab initio* reconstruction in cryoSPARC with no symmetry imposed, maximum and initial resolutions set to 6 Å and 12 Å respectively, initial and final minibatch sizes set to 1000 images. The resulting C1 *ab initio* map and particles then underwent NUR in cryoSPARC with no symmetry applied or with I symmetry applied. The NUR map with no symmetry has a 18.44 Å resolution and the NUR map with I symmetry applied has a 12.18 Å resolution; both maps were similar, justifying imposing I symmetry for the final reconstruction.

All resolutions are reported based on the gold-standard Fourier shell correlation FSC (GSFSC) = 0.143 criterion (*58*, *59*) and FSC curves were corrected for the effects of soft masking by high-resolution noise substitution (*60*). A summary of EM data acquisition and processing is provided in Tables S3-S4.

**Computational design of AbCs concurrent with oligomer design**

Given the success in designing AbCs when using only previously-validated oligomers, we were curious whether we could design structures with newly-designed cyclic oligomers. This has the advantage of creating oligomer building blocks for future applications as well as additional antibody nanocages. First, C3s were generated by docking helical proteins into cyclic symmetries and designing a low-energy de novo interface (*20*). Those C3s were used to design 48 antibody nanocages across D3 dihedral (14), T32 tetrahedral (11), O32 octahedral (15), and I32 icosahedral (8) symmetries following the same fusion and design approach described above. From these designs, 36 were soluble, and two D3 dihedra (Fig. S5a) formed with Fc into structures very similar to the designed models according to SEC, SAXS, and NS-EM (Fig. S5b-e).

**Stability experiments**

Samples were prepared for stability analysis by mixing equimolar amounts of each AbC-forming design component with hIgG1 Fc domain. These were purified using SEC using a Superose 6 column, following similar techniques as described above, into tris buffered saline (150 mM NaCl, 25 mM Tris-HCl pH 8.0) with 50 mM L-arginine (from a 1 M L-arginine pH 8.0); L-arginine was added to all designs as it had been observed to reduce hydrophobic association for the i52.6 AbCs. After SEC, the fractions corresponding to the AbC (left-most peak) were pooled. These were incubated at room temperature and analyzed once per week for up to five weeks post-SEC via DLS and SDS-PAGE. Designs d2.3 and d2.4 experiments were started three weeks later than the other six designs.

*Dynamic light scattering*

Dynamic light scattering measurements (DLS) were performed using the default Sizing and Polydispersity method on the UNcle (Unchained Labs). 8.8 μL of AbCs were pipetted into the provided glass cuvettes. DLS measurements were run in triplicate at 25°C with an incubation time of 1 second; results were averaged across runs and plotted using Graphpad Prism. Table S6 provides DLS summary data.

*SDS-PAGE*

10 uL of Fc AbCs were diluted to approximately 0.1 mg/mL and prepared for SDS by mixing with 2 μL of 6× loading dye (197 mM Trs-HCl, pH 6.8; 70% glycerol; 6.3% SDS; 0.03% bromephenol blue); these were then heated for 5-10 minutes at 95°C and loaded into the wells of a Tris-Glycine gel (Bio-Rad, catalogue #5678125). SDS running buffer was prepared to a final concentration of 5 mM Tris, 192 mM glycine, 0.1% SDS, pH 8.3. 2-5 uL of ladder was also added (BioRad 161-0377 or 161-0374). The gel was run for 25-30 minutes at 180-200 V or until the dye reached near the bottom of the gel. Gels were stained with Coomassie Brilliant Blue dye using the Genscript eStain protein staining system.

*Exchange experiments*

GFP-Fc and RFP-Fc were produced in Expi293F cells and purified as described above. GFP-Fc was mixed with o42.1 tetramer; a pre-mixed ratio of RFP-Fc and GFP-Fc (at a 25:1 molar ratio) was separately combined with o42.1 tetramer as a positive control meant to mimic 100% exchange (as the GFP-Fc o42.1 AbC would be mixed with 25-fold excess RFP-Fc). Fc-GFP o42.1 and 25:1 Fc-RFP:GFP o42.1 were purified via a Superose 6 SEC column into TBS (150 mM NaCl, 25 mM Tris-HCl pH 8.0) with 50 mM L-arginine. Fc-GFP o42.1 was then incubated with 25-fold excess Fc-RFP at a final volume of 2 mL and separated using an autosampler set to inject 470 µL; the autosampler was necessary to control injection volume (Cytiva ALIAS autosampler). Time points were taken at 5 minutes, 2 hours, 4 hours, and 24 hours after mixing and incubation at 25°C. Controls were GFP-Fc o42.1 without added Fc-RFP, Fc-RFP without AbC, and the “pre-exchanged” control normalized to the GFP-Fc o42.1 molarity. 100 µL from each peak fraction were then added to a 96-well fluorescence plate (Corning, black polystyrene). To measure GFP signal, excitation and emission wavelengths were set to 485/510 (respectively); for RFP signal, excitation and emission wavelengths were set to 558/605; fluorescence readings were taken with the Neo2 Microplate Reader (BioTek).

**DR5 and A1F-Fc experiments**

*Cell culture*

Colorectal adenocarcinoma cell line-Colo205, and renal cell carcinoma cell line RCC4 were obtained from ATCC. Primary kidney tubular epithelial cells RAM009 were a gift from Dr. Akilesh (University of Washington). Colo205 cells were grown in RPMI1640 medium with 10% Fetal Bovine Serum (FBS) and penicillin/streptomyocin. RCC4 cells were grown in Dulbecco’s Modified Eagle’s Medium with 10% FBS and penicillin/streptomyocin. RAM009 were grown in RPMI with 10% FBS, ITS-supplement, penicillin/streptomyocin and Non Essential Amino Acids (NEAA). All cell lines were maintained at 37°C in a humidified atmosphere containing 5% CO2.

Human Umbilical Vein Endothelial Cells (HUVECs, Lonza, Germany, catalog # C2519AS) were grown on 0.1% gelatin-coated 35 mm cell culture dish in EGM2 media. Briefly, EGM2 consist of 20% Fetal Bovine Serum, 1% penicillin-streptomycin, 1% Glutamax (Gibco, catalog #35050061), 1% endothelial cell growth factor (*32*), 1mM sodium pyruvate, 7.5mM HEPES, 0.08mg/mL heparin, 0.01% amphotericin B, a mixture of 1× RPMI 1640 with and without glucose to reach 5.6 mM glucose concentration in the final volume. Media was filtered through a 0.45-μm filter. HUVECs at passage 7 were utilized in Tie2 signaling experiments. HUVECs at passage 6 were used in the tube formation assay.

*Caspase-Glo 3/7 and Caspase-Glo 8 assays*

Cells were passaged using trypsin and 40,000 cells/well were plated onto a 96-well white tissue culture plate and grown in appropriate media. Medium was changed the next day (100 μL/well) and cells were treated with either uncaged α-DR5 AMG655 antibody (150nM), recombinant human TNF Related Apoptosis Inducing Ligand (TRAIL; 150nM), Fc-only AbCs or α-DR5 AbCs (150nM, 1.5nM, 15pM for caspase-3/7; only 150 nM and 1.5 nM were tested for caspase-8) and incubated at 37°C for 24 hours (caspase-3/7) or 12 hours (caspase-8). In all cases here and throughout, the antibody or AbC concentration refers to the protein's asymmetric unit (e.g, the molar unit for the antibody is 1 heavy chain and 1 light chain). The following day, 100 μL/well of Caspase-Glo 3/7 reagent or Caspase-Glo 8 reagent (Promega, USA) was added into the media and incubated for 1 hour (caspase-3/7) or 2 hours (caspase-8) at 37°C. Luminescence was then recorded using Perkin EnVision microplate reader (Perkin Elmer). Statistical comparisons were performed using Graphpad Prism (see Table S8 for full detail).

*Titer Glo cell viability assay (4 d viability)*

Cells were plated onto a 96-well plate at 20,000 cells/well. The next day, cells were treated with 150nM of α-DR5 AbCs, TRAIL and α-DR5 antibody for 4 d. At day 4, 100 μL of CellTiter-Glo reagent (Promega Corp. USA, #G7570) was added to the 100 μL of media per well, incubated for 10 min at 37°C and luminescence was measured using a Perkin-Elmer Envision plate reader.

*Alamar Blue cell viability assay (6 d viability)*

Cells were seeded onto a 12-well tissue culture plate at 50,000 cells/well. The next day, cells were treated with α-DR5 AbCs, TRAIL, or α-DR5 antibodies at 150 nM concentration. Three days later cells were passaged at 30,000 cells/well and treated with 150 nM of α-DR5 cages, TRAIL and α-DR5 antibody for 3 days. At 6 days, the media was replaced with 450 μL/well of fresh media and 50 μL of Alamar blue reagent (Thermofisher Scientific, USA, #DAL1025) was then added. After 4 hours of incubation at 37°C, 50 μL of media were transferred into a 96-well opaque white plate and fluorescence intensity was measured using plate reader according to manufacturer's instructions.

*Protein isolation for western blot analysis*

Cells were passaged onto a 12-well plate at 80,000 cells/well and were grown until 80% confluency is reached. Before treatment the media was replaced with 500 µL of fresh media. For DR5 experiments, AMG-655 antibody and TRAIL were added at 150 nM concentration and Fc-only nanocages or α-DR5 nanocages were added at 150 nM, 1.5 nM and 15 pM concentration onto the media and incubated for 24 hours at 37°C prior protein isolation; as above, concentrations are calculated based on the asymmetric unit. For the caspase inhibition experiment, RCC4 cells were pre-treated for 30 minutes with 10 µM of zVAD followed by treatment with 150nM o42.1 α-DR5 Ab for additional 24 hours, and total protein isolation.

Media containing dead cells was transferred to a 1.5 mL Eppendorf tube, and the cells were gently rinsed with 1× phosphate buffered saline. 1× trypsin was added to the cells for 3 minutes. All the cells were collected into the 1.5 mL Eppendorf containing the medium with dead cells. Cells were washed once in PBS 1× and lysed with 70 µL of lysis buffer containing 20 mM Tris-HCl (pH 7.5), 150 mM NaCl, 15% Glycerol, 1% Triton, 3% SDS, 25 mM β-glycerophosphate, 50mM NaF, 10mM Sodium Pyrophosphate, 0.5% Orthovanadate, 1% PMSF (all chemicals were from Sigma-Aldrich, St. Louis, MO), 25 U Benzonase Nuclease (EMD Chemicals, Gibbstown, NJ), protease inhibitor cocktail (PierceTM Protease Inhibitor Mini Tablets, Thermo Scientific, USA), and phosphatase inhibitor cocktail 2 (catalog#P5726), in their respective tubes). Total protein samples were then treated with 1 µL of Benzonase (Novagen, USA) and incubated at 37°C for 10 minutes. 21.6 µL of 4x Laemmli Sample buffer (Bio-Rad, USA) containing 10% beta-mercaptoethanol was added to the cell lysate and then heated at 95°C for 10 minutes. The boiled samples were either used for western blot analysis or stored at -80 °C.

*Western blotting*

The protein samples were thawed and heated at 95°C for 10 minutes. 10 μL of protein sample per well was loaded and separated on a 4-10% SDS-PAGE gel for 30 minutes at 250 V. The proteins were then transferred onto a Nitrocellulose membrane for 12 minutes using the semi-dry turbo transfer western blot apparatus (Bio-Rad, USA). Post-transfer, the membrane was blocked in 5% nonfat dry milk for 1 hour. After 1 hour, the membrane was probed with the respective antibodies: cleaved-PARP (Cell Signaling #9541, USA) at 1:2000 dilution; Cleaved-caspase 8 (Cell signaling #9496, USA) at 1:2000 dilution; pERK1/2 (Cell Signaling) at 1:5000 dilution; pFAK (Cell Signaling) at 1:1000 dilution; p-AKT(S473) (Cell Signaling) at 1:2000 dilution; and actin (Cell Signaling, USA) at 1:10,000 dilution. Separately, for p-AKT(S473) the membrane was blocked in 5% BSA for 3 hours followed by primary antibody addition. Membranes with primary antibodies were incubated on a rocker at 4°C, overnight. Next day, the membranes were washed with 1× TBST (3 times, 10 minutes interval) and the respective HRP-conjugated secondary antibody (Bio-Rad, USA) (1:10,000) was added and incubated at RT for 1 hour. For p-AKT(S473), following washes, the membrane was blocked in 5% milk at room temperature for 1 hour and then incubated in the respective HRP-conjugated secondary antibody (1:2000) prepared in 5% milk for 2 hours. After secondary antibody incubation, all the membranes were washed with 1× TBST (3 times, 10 minutes interval). Western blots were developed using Luminol reagent (Immobilon Western Chemiluminescent HRP Substrate, Millipore) for 3-15 seconds and imaged using Bio-Rad ChemiDoc Imager. Data were quantified using the ImageJ software to analyze band intensity.

Quantifications were done by calculating the peak area for each band. Each signal was normalized to the actin quantification from that lane of the same gel, to allow for cross-gel comparisons. Fold-changes were then calculated compared to PBS for all samples except for the pAKT reported for the A1F-Fc western blot (there was not enough pAKT signal for comparison, so o42.1 A1F-Fc was used for normalization). Statistical comparisons were performed using Graphpad Prism (see Tables S8, S9 for full detail). For all statistical analyses, means were compared to the PBS condition.

*Tube formation assay (vascular stability)*

Vascular stability and tube formation were assessed using a protocol modified from a previous report (*61*). Briefly, passage 6 HUVECs were seeded onto 24-well plates precoated with 150 µL of 100% cold Matrigel (Corning, USA) at 150,000 cells/well density, along with scaffolds at 89 nM F-domain concentrations or PBS in low glucose DMEM medium supplemented with 0.5% FBS for 24 hours. At the 24 hour time point, old media was aspirated and replaced with fresh media without added AbCs or controls. The cells were incubated up to 72 hours. Cells were imaged at 48 hour and 72 hour time points using Leica Microscope at 10× magnification under phase contrast. Thereafter, the tubular formations were quantified by calculating the number of nodes, meshes and tubes using the Angiogenesis Analyzer plugin in Image J software. Vascular stability was calculated by averaging the number of nodes, meshes, and tubes, and then normalizing to PBS. Statistical comparisons were performed using

Graphpad Prism (see Table S9 for full detail).

*Human serum A1F-Fc AbC incubation experiment*

HUVECs (C2519AS, Lonza) were grown to at least 80% confluence in 24-well plate format pre-treated with attachment factor (S006100, ThermoFisher) and cultured in EGM-2 growth medium (CC-3162, Lonza) according to manufacturer's instructions. The cells were then starved in DMEM low glucose serum-free media (11885084, Gibco) for 24 hrs. In parallel, proteins were incubated in 100% human serum (Sigma, H4522-100ML) at 1.5 μM for 24 hours at 4°C or 37°C; dilutions of AbC into serum were approximately 1:4 (AbC to final, v/v). After starvation and protein incubation, cell media was replaced, and proteins were added to the cells at a final concentration of 150 nM for 30 minutes at 37°C. Conditions with human serum were all normalized to a final concentration of 10% upon addition to the cell media. After treatment, the media was aspirated and cells were washed once with PBS before lysis. Cells were lysed with 60 μL of lysis buffer containing 50 mM HEPES, 150 mM NaCl, 10% Glycerol, 1% Triton X-100, 3% SDS, 25 mM β-glycerophosphate, 100 mM NaF, 10 mM Sodium Pyrophosphate, 1 mM EGTA, 1.5 mM MgCl2, 1% Sodium Orthovanadate, 300 μM PMSF, 25 U DNase, 1% phosphatase inhibitor cocktail 2 (all chemicals were from Sigma-Aldrich), and protease inhibitor cocktail (PierceTM Protease Inhibitor Mini Tablets, Thermo Scientific, USA). Cell lysate was collected in a fresh Eppendorf tube. Lysate samples were prepared using the Anti-Rabbit Detection Module for the Jess instrument (ProteinSimple) and boiled for 10 minutes at 98°C. A 12-230 kDa 25-capillary cartridge and microplate were utilized for the Jess instrument, using the antiphospho-Akt (S473) (D9E) XP rabbit mAb (4060, Cell Signaling) with a 30 minute incubation time. Replicate chemiluminescent peak values corresponding to phospho-Akt (~56 kDa) are reported.

**Immune cell activation materials and methods**

*CD40 luminescence assay*

A non-agonistic antibody (clone LOB7/6, product code MCA1590T, BioRad), was combined with the octahedral o42.1 AbC-forming design as described above and further characterized by DLS and NS-EM (Fig. S10). Negative control o42.1 AbC was made using a non-CD40 binding IgG (mpe8), which binds to RSV spike protein (*44*). These two AbCs, along with uncaged LOB7/6 and a positive control CD40-activating IgG (Promega, catalog #K118A) were diluted to make a 10-point, threefold dilution series for triplicate technical repeats starting at 1.2 μM; as described above, concentrations are calculated based on the asymmetric unit. The positive control CD40-activating IgG (K118A) is a murine IgG1a antibody, it was not compatible for assembly with the o42.1 design, likely due to the low binding interface between protein A and mIgG1a (data not shown). Particles were filtered using a 0.22 μm syringe filter (Millipore) and AbC formation was assessed using SEC and DLS using procedures described above. SEC was used as an analytical technique to show absence of unassembled components; due to the expense of commercial antibodies and the typical loss of yield using SEC as would be expected for any filtration technique, we did not use SEC as a separation technique here prior to DLS measurements or in vitro assays. SEC and DLS confirm the presence of the designed assemblies and absence of off-target or unassembled species; the o42.1 AbCs eluted in the SEC void of the Superose 6 column as expected given their designed and verified radii (~40 nm when formed with IgGs). Post-filtration concentration readings were taken and confirmed that there was no sample loss when using the syringe filter.

To assay CD40 activation, we followed manufacturer's instructions for a bioluminescent cell-based assay that measures the potency of CD40 response to external stimuli such as IgGs (Promega, JA2151). Briefly, CD40 effector Chinese Hamster Ovary (CHO) cells were cultured and reagents were prepared according to the assay protocol. The antibodies and AbCs were incubated with the CD40 effector CHO cells for 8 hours at 37ºC, 5% CO2. Bio-Glo™ Luciferase Assay System (G7941) included in the assay kit is used to visualize the activation of CD40 from luminescence readout from a plate reader. The Bio-Glo™ Reagent is applied to the cells and luminescence was detected by a Synergy Neo2 plate reader every min for 30 minutes. Data were analyzed by averaging luminescence between replicates and subtracting plate background. The fold induction of CD40-binding response was determined by RLU of sample normalized to RLU of no antibody controls. Data curves were plotted and EC50 was calculated using GraphPad Prism using the log(agonist) vs. response -- Variable slope (four parameters); see Table S10 for EC50 values and 95% CI values.

*T cell proliferation and flow cytometry*

Mosaic AbCs were formed by mixing α-CD3 (clone name: OKT3, BioLegend) and α-CD28 (CD28.6, catalog #16-0288-85, ThermoFisher) antibodies together first, and then combining with excess o42.1 AbC-forming design. Mosaic α-CD3/28 o42.1 cages were purified via SEC into PBS as described above. SEC and DLS confirmed the assembly of o42.1 AbCs, which eluted as expected into the void volume in SEC given the particle's size.

Primary human peripheral blood mononuclear cells (PBMC) were obtained upon written informed consent from the Virginia Mason Medical Center in Seattle, WA, USA. All studies were approved by the Institutional Review Board of the Benaroya Research Institute (Seattle, WA). Naive $CD4^+$ conventional human T cells ($CCR7^+CD45RA^+CD127^{hi}CD25^{neg}$) were isolated from PBMC by cell sorting to >99% purity. PBMC were first labeled with 2.5 μM Cell Proliferation Dye e670 (ThermoFisher) according to manufacturer instructions, then rested for 1h at 37C 5% $CO_2$. CPD-labeled cells were harvested, incubated with viability dye ef780 (ThermoFisher) and stained in buffer containing HBBS + 0.3% BSA with indicated fluorescently labeled surface markers. Cell sorting and analysis were performed on a FACSAria Fusion (BD Biosciences) using an 85 μM nozzle at 45 psi. Sorted T cells (1e6/mL) were incubated in the presence of indicated stimulation conditions (0.01μM) in ImmunoCult-XF T Cell Expansion Medium (Stem Cell). After 4-5 days, cells were harvested and re-stained with fluorescent antibodies. Data were analyzed using FlowJo software (Tree Star, Inc.)

**Viral neutralization**

*CV1, CV3, CV30*

α-CoV-2 S cages using CV IgGs were prepared by mixing α-CoV-2 S IgGs with a 1:1 molar ratio of o42.1 design component and purifying via SEC into TBS, following similar protocols to those as described above for AbC assembly.

HIV-1 derived viral particles were pseudotyped with full length wildtype SARS CoV-2 S (*62*). Briefly, plasmids expressing the HIV-1 Gag and pol (pHDM-Hgpm2, BEI resources Cat# NR-52517), HIV-1Rev (pRC-CMV-rev1b, BEI resources Cat# NR-52519), HIV-1 Tat (pHDM-tat1b, BEI resources Cat# NR-52518), the SARS CoV2 spike (pHDM-SARS-CoV-2 Spike, BEI resources Cat# NR-52514) and a luciferase/GFP reporter (pHAGE-CMV-Luc2-IRES-ZsGreen-W, BEI resources Cat# NR-52516) were co-transfected into 293T cells at a 1:1:1:1.6:4.6 ratio using 293 Free transfection reagent (EMD Millipore Cat# 72181) according to the manufacturer's instructions. Transfected cells were incubated at 32ºC for 72 hours after which the culture supernatant was harvested, clarified by centrifugation and frozen at -80˚C.

293 cells stably expressing ACE2 (HEK-293T-hACE2, BEI resources Cat# NR-5251) were seeded at a density of $4\times10^3$ cells/well in a 100 µL volume in 96 well flat bottom clear bottomed, black walled plates (Greiner Bio-One Cat # 655090) (*62*). The next day, IgG alone, or in complex with cage components were serially diluted in 30µl of cDMEM in 96 well round bottom plates in triplicate; as described above, concentrations are calculated based on the asymmetric unit.

An equal volume of viral supernatant was added to each well and incubated for 60 min at 37ºC. Meanwhile 50 µL of cDMEM containing 6µg/ml polybrene was added to each well of 293T-ACE2 cells (2µg/ml final concentration) and incubated for 30 min. The media was aspirated from 293T-ACE2 cells and 100µl of the virus-antibody mixture was added. The plates were incubated at 37˚C for 72 hours. The supernatant was aspirated and replaced with 100 µL of Steadyglo luciferase reagent (Promega) and the plate was read on a Fluorskan Ascent Fluorimeter. Control wells containing virus but no antibody (cells + virus) and no virus or antibody (cells only) were included on each plate.

Percent neutralization for each well was calculated as the RLU of the average of the cells + virus wells, minus test wells (cells + mAb + virus), and dividing this result difference by the average RLU between virus control (cells + virus) and average RLU between wells containing cells alone, multiplied by 100. The antibody concentration that neutralized 50% of infectivity (IC50) was interpolated from the neutralization curves determined using the log(inhibitor) vs. response -- Variable slope (four parameters) fit using Graphpad Prism Software. Experiments were performed in duplicate. See Table S4 for IC50 values and 95% CI values.

*Fc-ACE2*

Murine leukemia virus (MLV)-based SARS-CoV-2 S-pseudotyped viruses were prepared as previously described (*43*). Briefly, HEK293T cells were co-transfected with a SARS-CoV-2 S encoding-plasmid, an MLV Gag-Pol packaging plasmid and the MLV transfer vector encoding a luciferase reporter using the Lipofectamine 2000 transfection reagent (Life Technologies) according to the manufacturer's protocols. Transfection mixture was added dropwise to HEK293T cells. Cells were then incubated in the transfection mixture and OPTI-MEM for 5 hours at 37°C with 8% CO2 before the medium was exchanged into DMEM containing 10% FBS. After 72 hours, the pseudovirus-containing supernatant was collected, centrifuged for 10 minutes at 3000×g to clear cell debris and filtered using a 0.45um filter with PES-membrane (MilliporeSigma). The pseudoviruses were concentrated using 30 kDa cut-off concentrators (Amicon) and stored at -80°C until further use.

HEK-293T-hACE2 (BEI resources Cat# NR-5251) were cultured in DMEM containing 10% FBS and 1% PenStrep (*62*). 16-24 hours before infection, cells were plated into white sided clear bottom 96-well plates coated with Poly-L-Lysine solution (Sigma Aldrich, Cat #: P4707). Briefly, 25 μL Poly-L-Lysine solution was added to each well. The plate was incubated at RT for 10 minutes before removal of Poly-L-Lysine and washing with tissue culture grade water. The Poly-L-lysine coated plate was dried for 10 minutes before the cell plating step. Prior to transfection the HEK-293T-hACE2 96 well plates were washed 3 times with DMEM. Fc-ACE2 (Sinobiologicals, Cat #: 10108-H02H), o42.1 Fc, and o42.1 Fc-ACE2 were purified via SEC as described above, and serially diluted 2× in DMEM starting from 800nM; all concentrations are calculated based on the asymmetric unit. Equal volumes of concentrated pseudovirus and serial dilution of treatments (Fc-ACE2, o42.1 Fc particles or o42.1 Fc-ACE2 particles or DMEM) were combined and incubated for 30 minutes and then added to the cells. After 2-3 hours, DMEM containing 20% FBS and 2% PenStrep was added to the cells. 48 hours post infection, One-Glo-EX (Promega) was added to the cells and incubated in the dark for 5-10 minutes prior to reading on a Varioskan LUX plate reader (ThermoFisher). As above, the antibody concentration that neutralized 50% of infectivity (IC50) was interpolated from the neutralization curves determined using the log(inhibitor) vs. response -- Variable slope (four parameters) using Graphpad Prism Software. The difference in IC50 was compared using the extra sum-of-squares F-test function in Prism with a P-value cutoff at 0.05. Experiments were performed in technical duplicate. See Table S11 for IC50 values and 95% CI values.

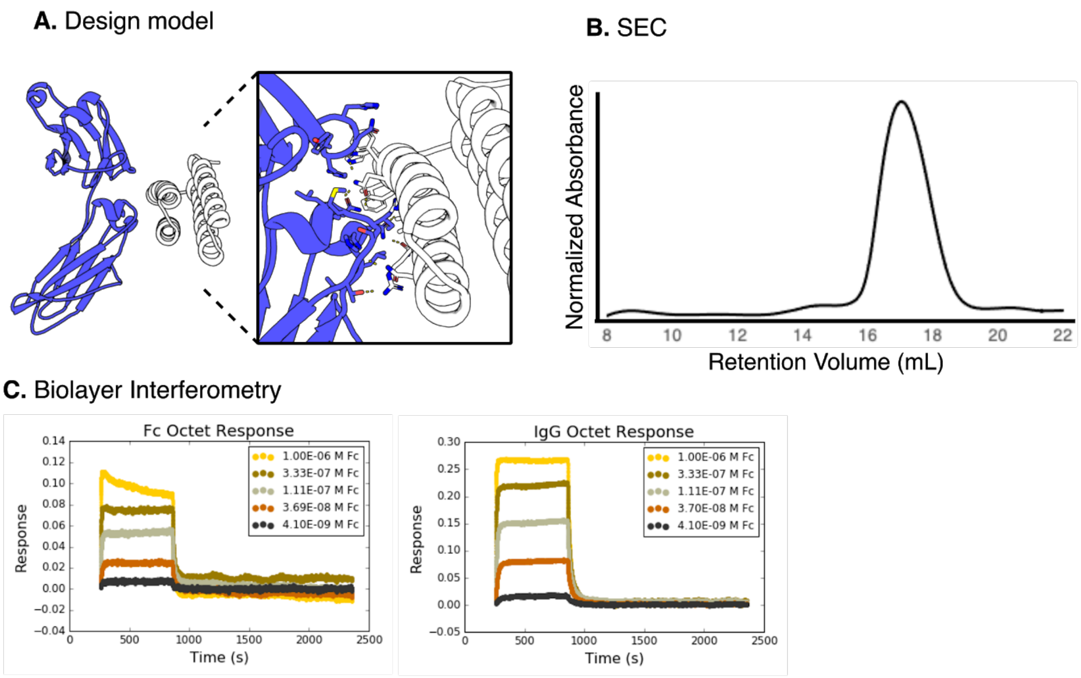


| Target | $K_D$ (M) | $K_D$ Error | $k_{on}$ (1/Ms) | $k_{on}$ Error | $k_{dis}$ (1/s) | $k_{dis}$ Error |
|---|---|---|---|---|---|---|
| Fc | 1.13E-07 | 7.69E-09 | 7.30E+05 | 4.74E+04 | 8.25E-02 | 1.68E-03 |
| IgG1 | 7.17E-08 | 1.24E-09 | 5.44E+05 | 9.03E+03 | 3.90E-02 | 1.92E-04 |

**Figure S1. Designed Fc-binding designed helical repeat. A,** Model of the helical repeat protein DHR79 docked against antibody Fc (PDB ID: 1DEE). Residues from protein A (PDB ID: 1L6X) are grafted at the interface between the Fc and the helical repeat protein. **B,** SEC trace of the Fc-binding helical repeat monomer. **C,** Biolayer interferometry (BLI) of the Fc-binding helical repeat design with Fc (left) or with hIgG1 (right), with summary statistics (below).

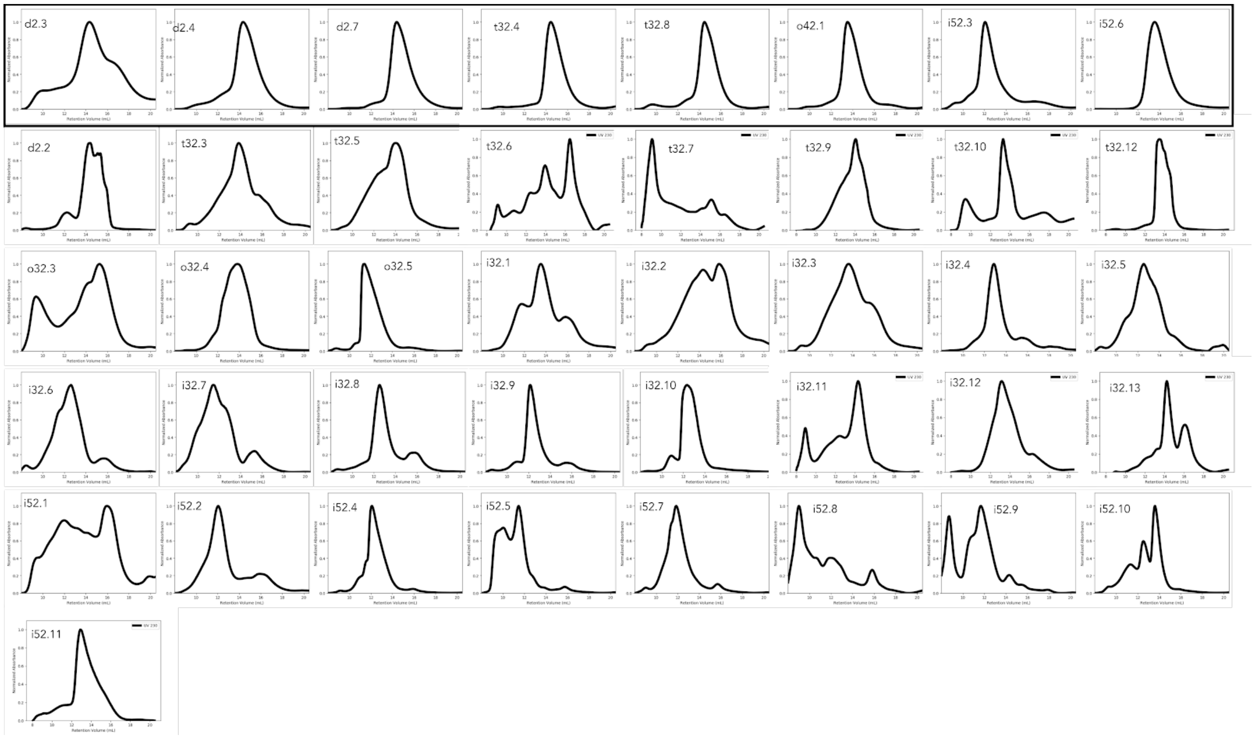


**Figure S2. SEC profiles for all soluble designs.** First row: SEC profiles of all soluble antibody nanocage-forming designs run over a Superose S200 column. Bottom rows: all other designs that did not form nanocages when mixed with antibodies. Several designs appear to still form oligomers at the expected size, but these may not have formed at the right orientation to lead to successful nanocage formation. X-axis for each is retention volume, and the Y-axis is normalized A230 absorbance.

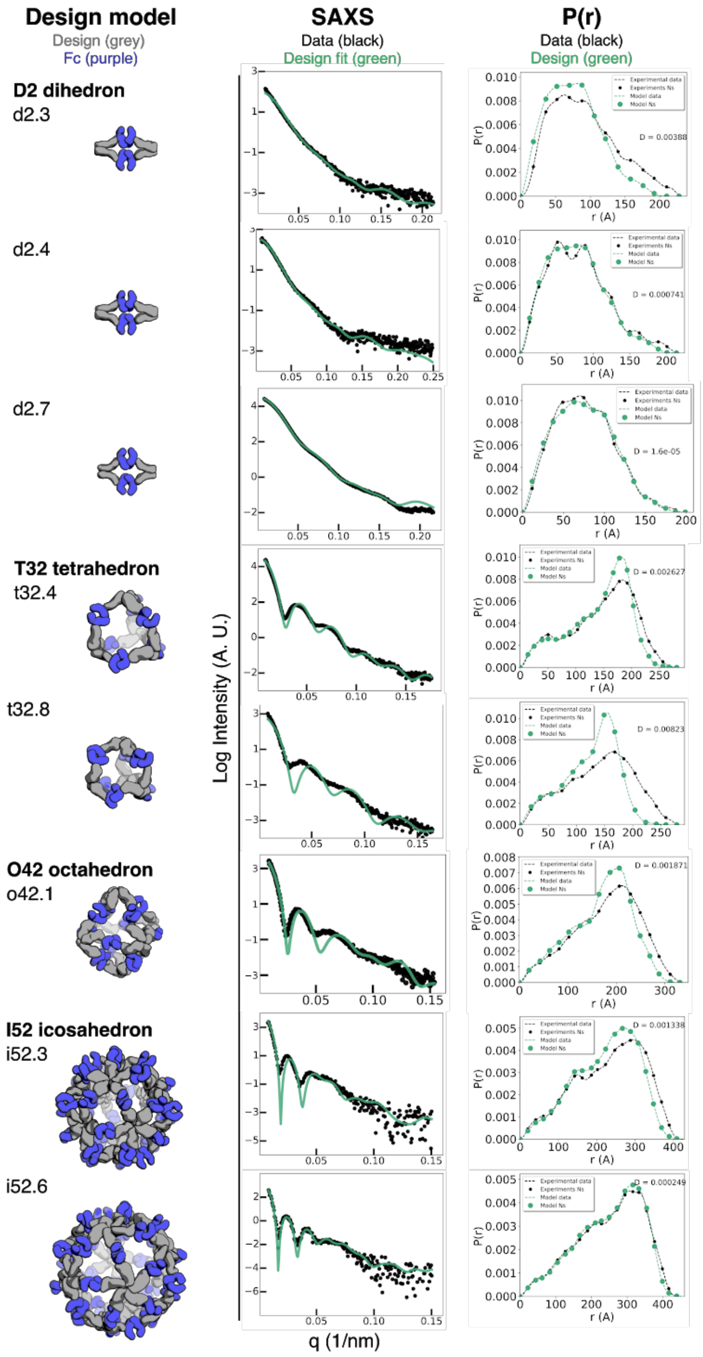


**Figure S3. SAXS profiles for AbCs formed with Fc.** Left: design models show designed Fc-binding oligomers in grey and antibody Fc in purple. Middle: small angle X-ray scattering (SAXS) curve fit for all designs with Fc; Black dots represent experimentally-determined data, and the green lines are calculated from the design models, with the FOXS server (*18*) used to perform the curve fitting. Right: Distance distribution functions (P(r) curves) for experimental data (black) compared to theoretical distribution functions from the design models (green); Scatter3 was used to perform the P(r) analyses.

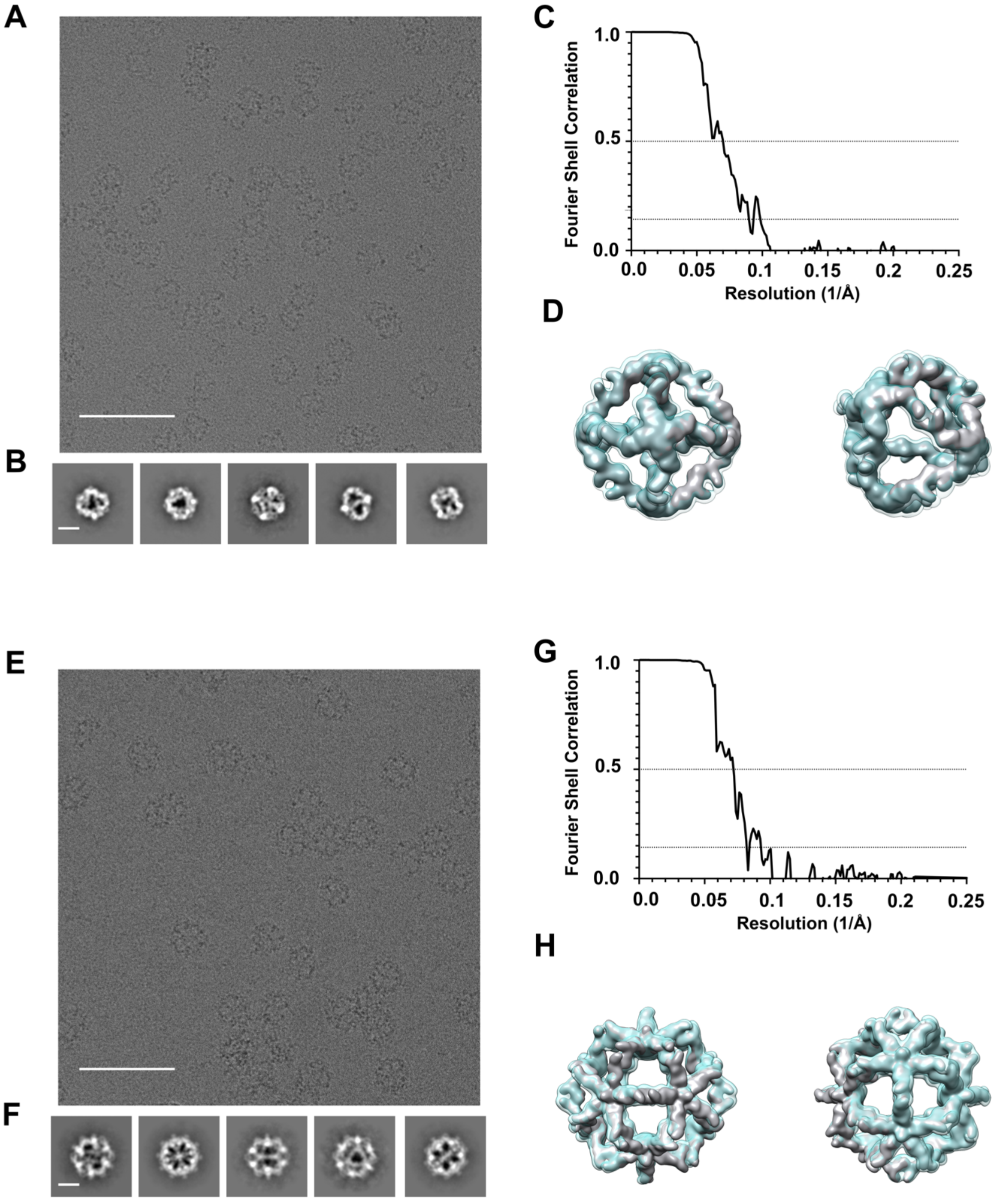


**Figure S4. Cryo-EM characterization of o42.1 Fc (A-D) and i52.3 Fc AbCs (E-H). A,** Representative cryo EM micrograph of o42.1 Fc. Scale bar: 100Å. **B,** Reference-free 2D class averages of o42.1-Fc. Scale bar, 200 Å. **C,** Gold-standard Fourier shell correlation curves for the o42.1 Fc map with O symmetry applied. Dotted line indicates the 0.143 and 0.5 thresholds. **D,** Two views of the o42.1 Fc cryo-EM map reconstructed with no symmetry (transparent cyan) superimposed on the o42.1 Fc cryo-EM map with O symmetry applied (solid gray). **E,** Representative micrograph of i52.3 Fc. Scale bar: 100Å. **F,** Reference-free 2D class averages of i52.3 Fc. Scale bar, 200 Å. **G,** Gold-standard Fourier shell correlation curves for the i52.3 Fc map with I symmetry applied. Dotted line indicates the 0.143 and 0.5 thresholds. **H,** Two views of the i52.3 Fc cryo-EM map reconstructed with no symmetry (transparent cyan) superimposed on the i52.3 Fc cryo-EM map with I symmetry applied (solid gray).

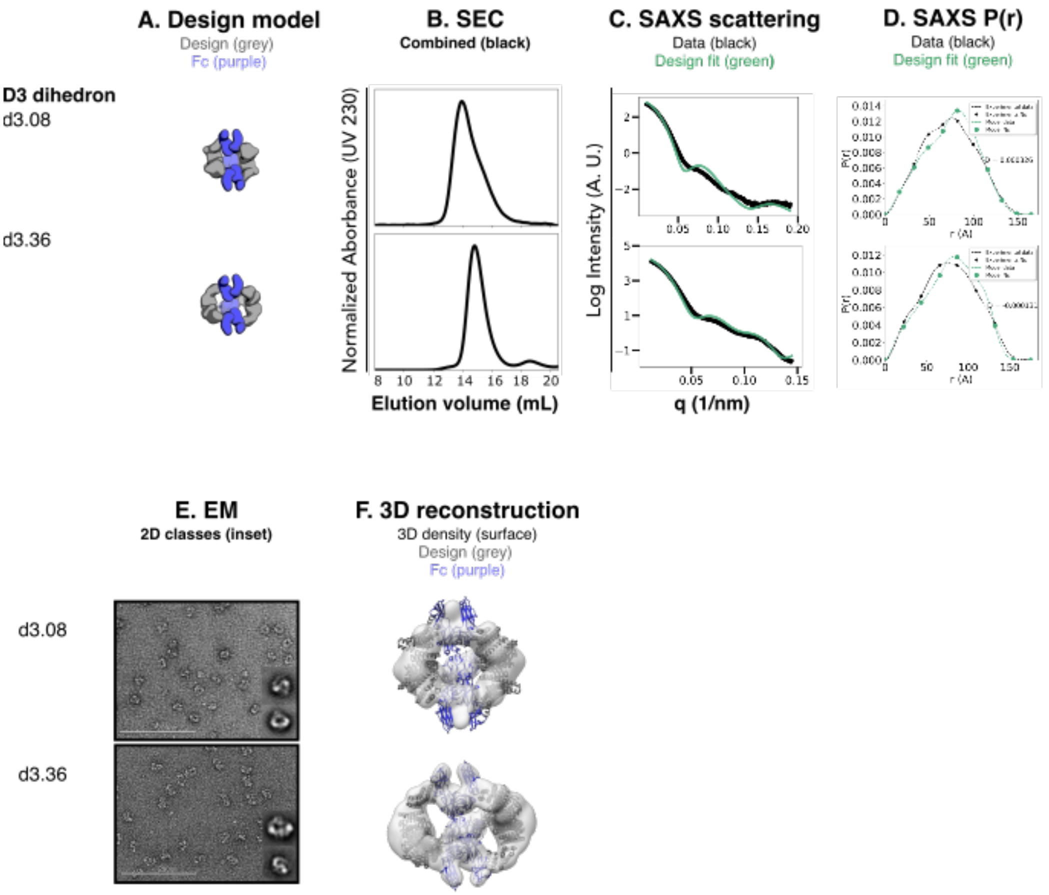


**Figure S5. Structural characterization of D3 dihedral AbCs with newly designed oligomers**. **A,** Design models, with antibody Fc (purple) and designed particle-forming oligomers (grey). **B,** SEC of the assembled AbC with Fc. **C.** SAXS curve fits for all designs with Fc; Black dots represent experimentally-determined data, and the green lines are calculated from the design models, with the FOXS server (*18*) used to perform the curve fitting. **D,** Distance distribution functions (P(r) curves) for experimental data (black) compared to theoretical distribution functions from the design models (green); Scatter3 was used to perform the P(r) analyses (reference). **E,** EM images with 2D averages in inset. **F,** 3D reconstructions from NS-EM data.

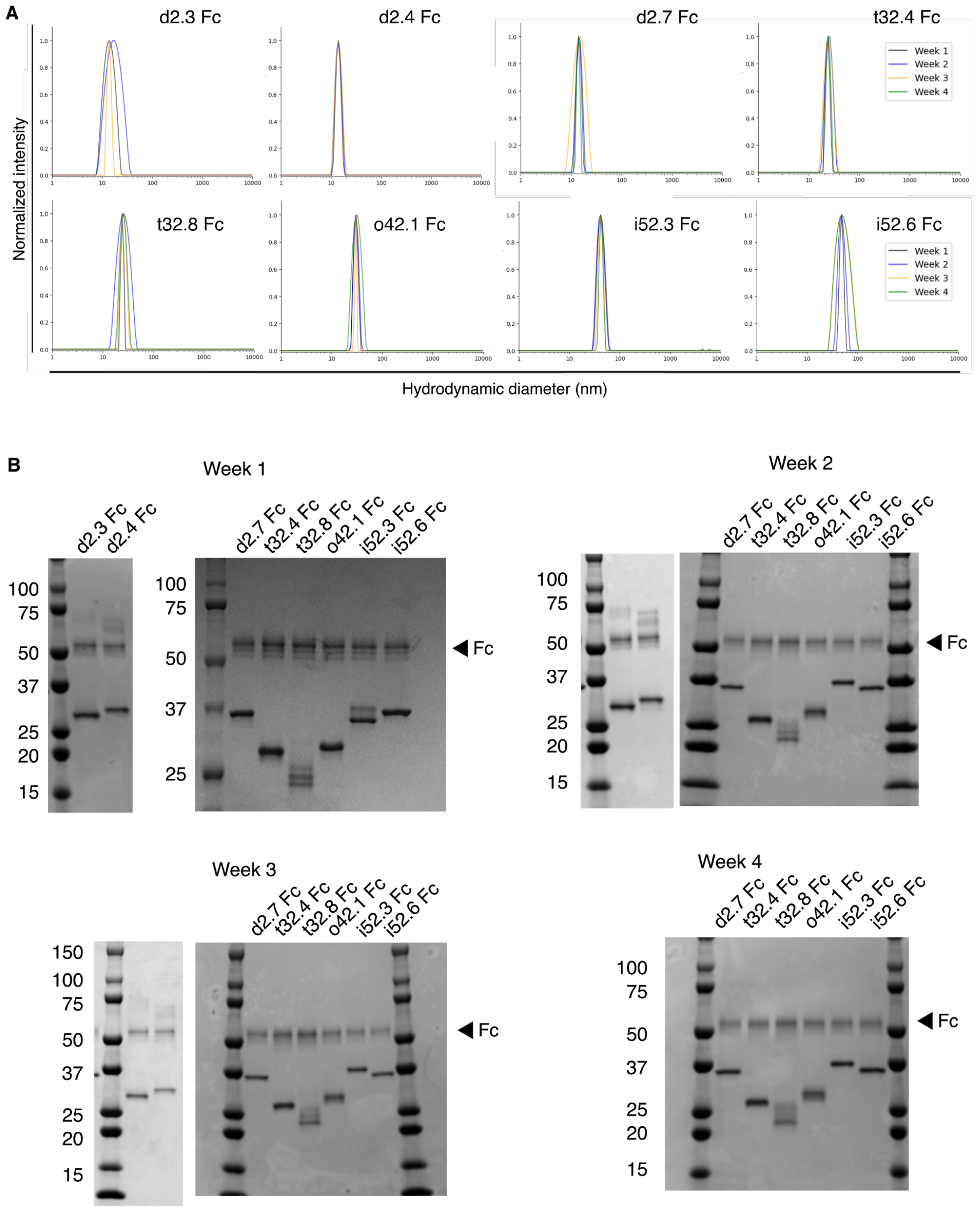


**Figure S6. Fc AbC particle stability over time. A,** Dynamic light scattering (DLS) of Fc AbCs, incubated at 25°C, and measured once per week. Traces are an average of 4 measurements each. **B,** SDS-PAGE analysis of Fc AbCs (without reducing agent), incubated at 25°C, and measured once per week. Molecular mass standards were run on outer lanes with masses (KDa) labeled.

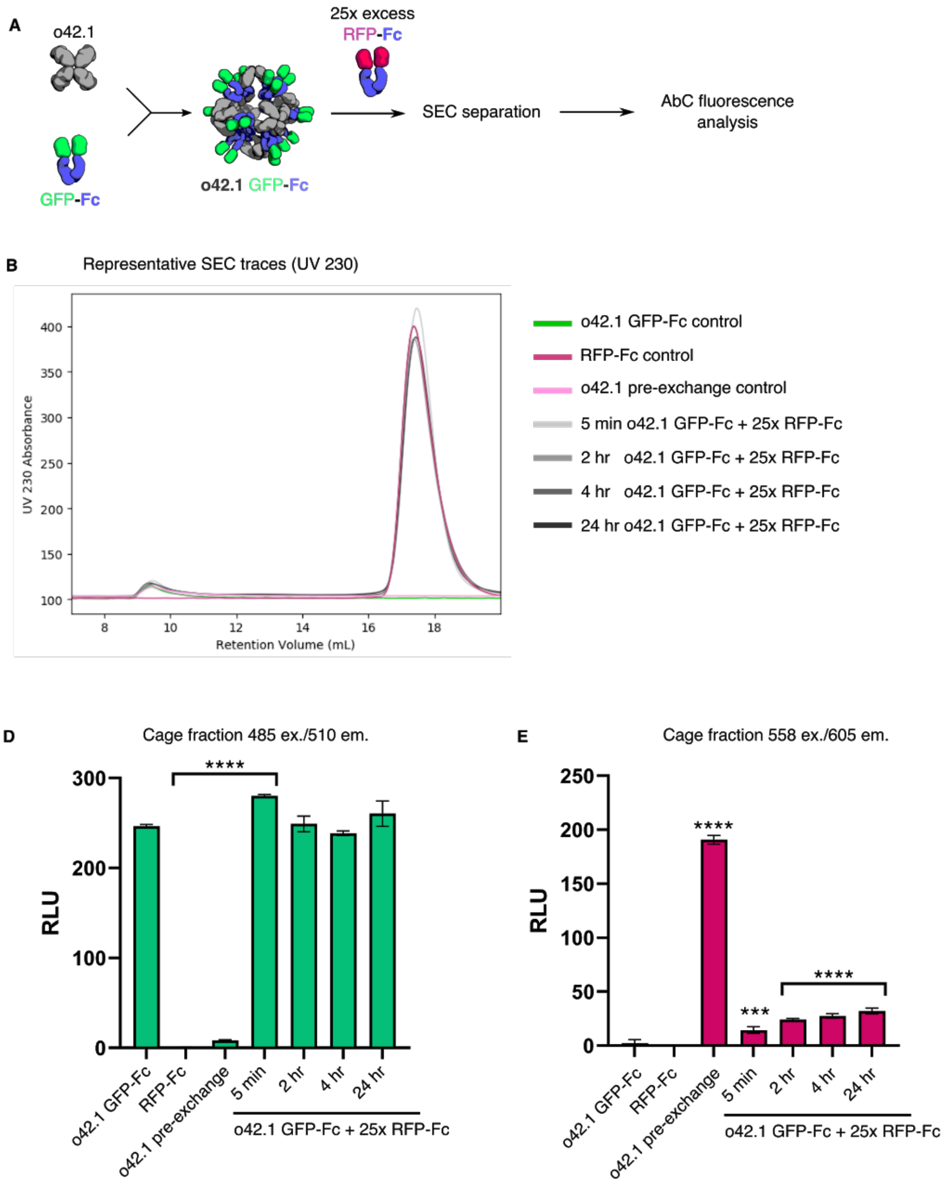


**Figure S7. o42.1 AbC formed with GFP-Fc minimally exchanges with free 25-fold excess RFP-Fc. A,** o42.1 AbCs are formed with GFP-Fc, purified, and incubated with 25-fold molar excess of RFP-Fc for up to 24 hours at room temperature. SEC is used to purify cage away from excess Fc-fusion; if AbCs exchange with free Fc-fusions, GFP signal would decrease in the cage fraction as RFP signal increases. Controls include: o42.1 GFP-Fc alone; RFP-Fc alone; and "pre-exchanged" o42.1 AbC prepared by pre-mixing RFP-Fc and GFP-Fc at a 25:1 molar ration prior to AbC formation. **B,** representative SEC traces showing UV 230 absorbance. **C,** GFP signal briefly increases in the cage fraction for the o42.1 GFP-Fc AbCs incubated with 25-fold excess RFP-Fc, but drops to control o42.1 GFP-Fc levels, which is maintained for 24 hours at room temperature. **D,** RFP signal is increased in the cage fraction of o42.1 GFP-Fc AbCs incubated with Fc-RFP by less than 20% over 24 hours.

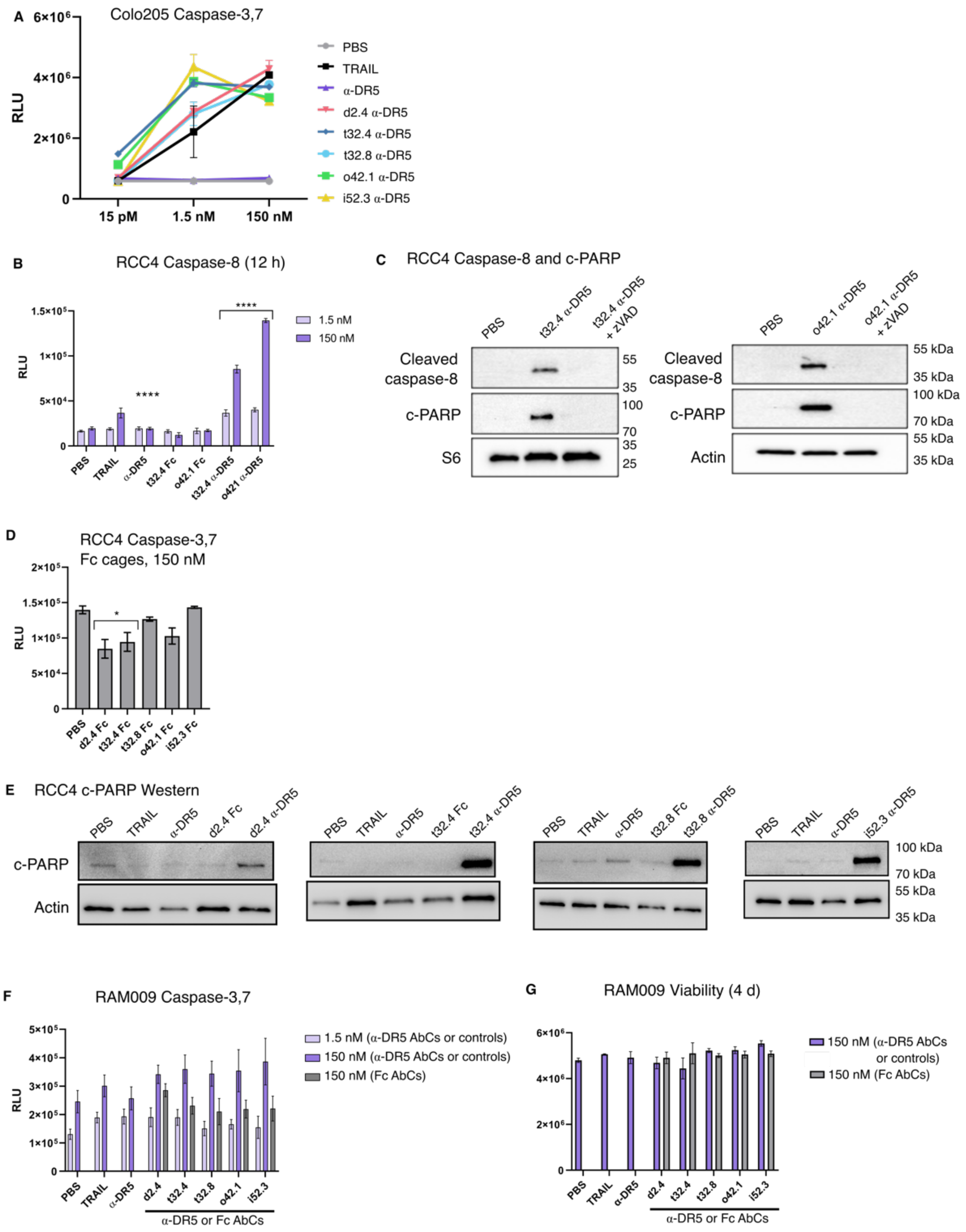


**Figure S8. Additional α-DR5 AbC experiments. A,** α-DR5 AbCs and TRAIL activate caspase-3,7 in Colo205 colorectal cancer cell lines. **B,** α-DR5 o42.1 AbCs activate caspase-8 after 12 hour incubation in t32.4 and o42.1 α-DR5 AbCs **C**, Cleaved-caspase 8 and cleaved-PARP inhibition after 24 hour incubation with t32.4 and o42.1 α-DR5 AbCs, and 10 μM zVAD, a caspase inhibitor (**C**). **D,** AbCs formed with Fc from hIgG1 do not activate caspase-3,7 at 150 nM in RCC4 cells. **E,** PARP is cleaved by α-DR5 AbCs in RCC4 cells, but not by TRAIL, α-DR5, or Fc AbCs. **F-G,** α-DR5 AbCs do not greatly activate caspase-3,7 after 2 d **(F)** or reduce viability (**G**) in a primary tubular kidney cell line (RAM009). Statistical analyses are reported in Table S8.

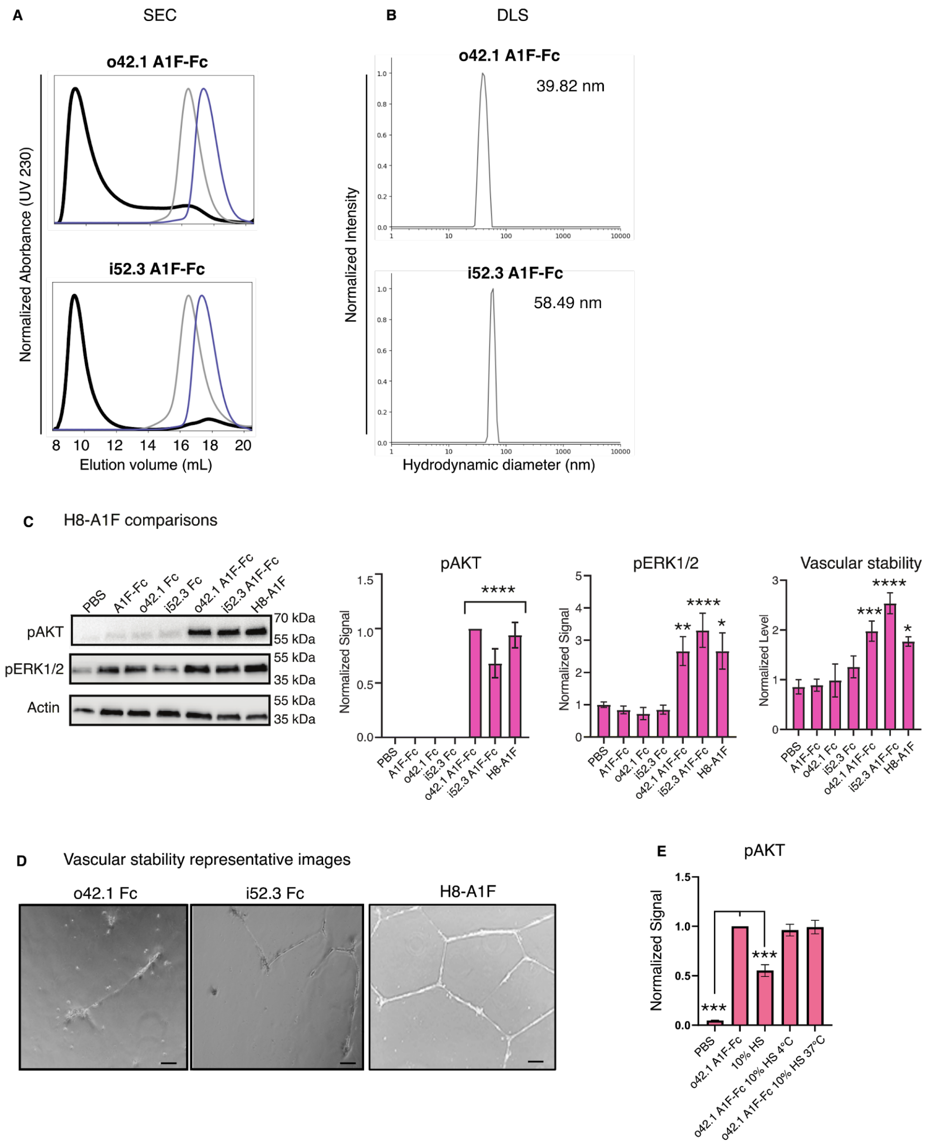


**Figure S9. Additional A1F-Fc AbC experiments. A-B,** o42.1 and i52.3 AbCs formed with A1F-Fc are monodisperse and of the expected size per SEC on a Superose 6 column (**A**) and DLS (**B**). SEC shows the assembly trace in black, the relevant AbC design component in grey, and the A1F-Fc in purple. **C**, A control assembly displaying 8 A1F ligands produced similar levels of pAKT and pERK1/2 activation along with a comparable increase in vascular stability. **D**, Representative images of o42.1 and i52.3 AbCs formed with Fc in the vascular stability assays. **E,** o42.1 A1F-Fc AbCs were incubated with 10% human serum (HS) for 24 hours at 4°C or 37°C and applied to HUVEC cells at 150 nM. pAKT signal showed no decrease from o42.1 A1F-Fc particles incubated with serum. Statistical analyses are reported in Table S9.

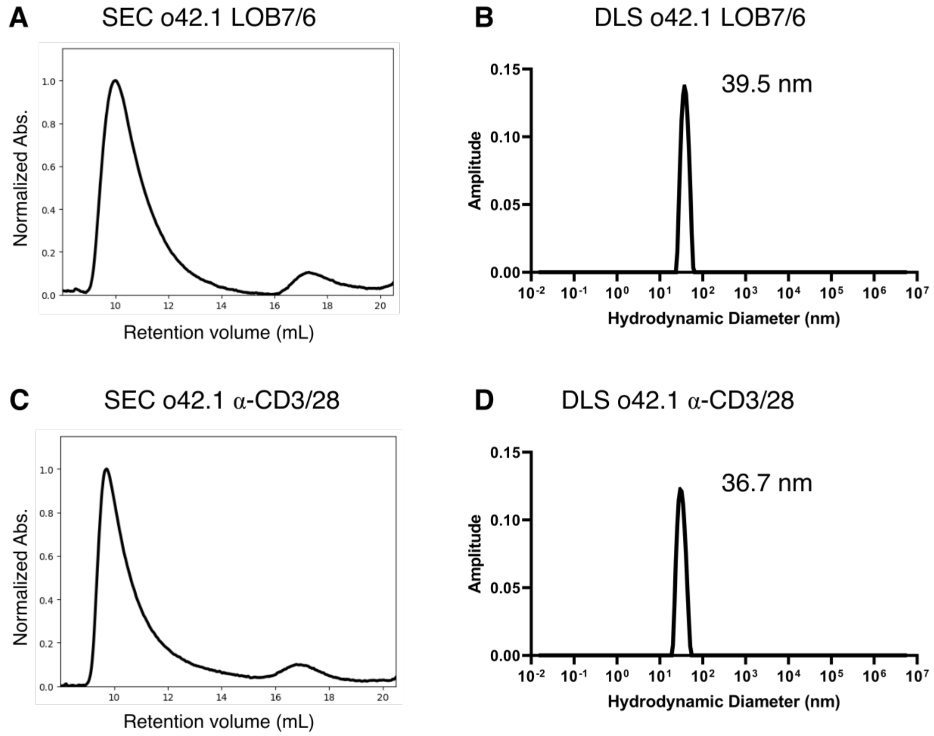


**Figure S10. Structural verification for immune stimulating AbCs (with α-CD40 or α-CD3/28). A-C,** Structural verification for α-CD40 AbCs formed with o42.1 and LOB7/6, using SEC on an Superose 6 column (**A**), DLS (**B**), and NS-EM (**C**). **D-E,** Structural verification for α-CD3/28 mosaic AbCs formed with o42.1, using SEC (**D**) and DLS (**E**), The excess peak corresponds on the right corresponds to unassembled component, likely o42.1 AbC-forming design (as excess was added).

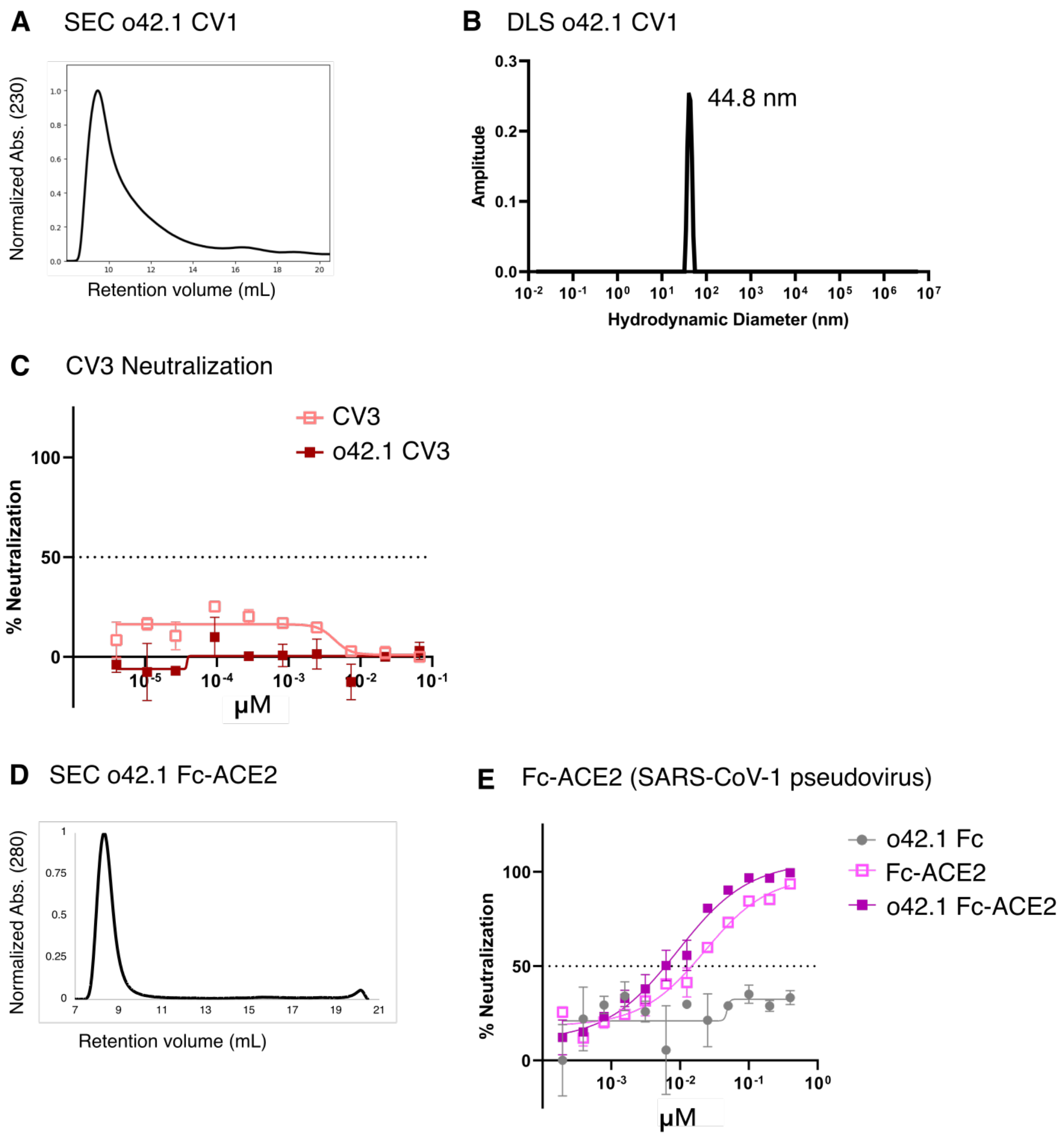


**Figure S11. Additional viral neutralization experiments. A-B,** Structural validation of o42.1 CV1 using SEC (**A**) and DLS (**B**). **C,** Neither o42.1 CV3 or free CV3 (α-CoV-2 S IgG, *31*) effectively neutralize SARS-CoV-2 pseudovirus. **D**, SEC characterization of o42.1 Fc-ACE2. **E**, o42.1 Fc-ACE2 is slightly more effective at neutralizing a SARS-CoV-1 pseudovirus compared to free Fc-ACE2.

### Supplemental Tables

**Table S1. Success rates of designed antibody-binding cage-forming oligomers.** Solubility (column 2) refers to the presence of protein in the post-lysis, post-centrifugation, pre-IMAC soluble fraction as read out by SDS gel. Good SEC component (column 3) refers to SEC traces with some peak corresponding to the approximate predicted size of the nanocage-forming design model. Data for cage formation with Fc are shown in Fig. 2 and 3.

| Geometry | # ordered | Soluble component | Good SEC component | Forms cage with Fc |
|---|---|---|---|---|
| D2 dihedron | 6 | 5 | 4 | 3 |
| T32 tetrahedron | 11 | 8 | 7 | 2 |
| O32 octahedron | 4 | 3 | 3 | 0 |
| O42 octahedron | 2 | 1 | 1 | 1 |
| I32 icosahedron | 14 | 14 | 10 | 0 |
| I52 icosahedron | 11 | 11 | 10 | 2 |
| Total | 48 | 42 | 35 | 8 |

**Table S2. Structural properties of designed models from SAXS analyses.** AbC design model predicted data (model) is compared against experimentally-derived SAXS data (exp) for radius of gyration (Rg) and $d_{max}$. The $q_{max}$ used for analysis is reported. All data was analyzed using Scatter3.

| Design | $R_g$ model (Å) | $R_g$ exp (Å) | $D_{max}$ model (Å) | $D_{max}$ exp (Å) | $q_{max}$ 1/nm |
|---|---|---|---|---|---|
| **d2.3** | 60.69 | 71.8 | 210 | 217 | 0.21 |
| **d2.4** | 60.67 | 63.88 | 210 | 214 | 0.25 |
| **d2.7** | 58.95 | 59.2 | 197 | 199 | 0.25 |
| **t32.4** | 107.27 | 112.02 | 280 | 282 | 0.25 |
| **t32.8** | 94.39 | 108.22 | 263 | 278 | 0.17 |
| **o42.1** | 126.48 | 135.1 | 320 | 331 | 0.16 |
| **i52.3** | 167.73 | 175.76 | 427 | 409 | 0.15 |
| **i52.6** | 187.32 | 188.9 | 454 | 443 | 0.15 |
| **d3.08** | 56.34 | 55.16 | 164 | 159 | 0.20 |
| **d3.36** | 60.38 | 59.32 | 168 | 175 | 0.15 |

**Table S3.** Details on EM data acquisition of different AbC samples.

| Sample name | Stain | Magnification | Pixel size (Å /pixiel) | # Micrographs |
|---|---|---|---|---|
| d2.3 Fc | UF | 67,000 | 1.6 | 300 |
| d2.4 Fc | UF | 67,000 | 1.6 | 234 |
| d2.7 Fc | nano-W | 67,000 | 1.6 | 251 |
| t32.4 Fc | nano-W | 67,000 | 1.6 | 359 |
| t32.8 Fc | nano-W | 67,000 | 1.6 | 739 |
| o42.1 Fc | cryo | 36,000 | 1.16 | 487 |
| i52.3 Fc | cryo | 36,000 | 1.16 | 460 |
| i52.6 Fc | nano-W | 52,000 | 2.07 | 342 |
| d2.3 hIgG1 | UF | 67,000 | 1.6 | 331 |
| d2.4 hIgG1 | UF | 67,000 | 1.6 | 160 |
| d2.7 hIgG1 | nano-W | 67,000 | 1.6 | 206 |
| t32.4 hIgG1 | nano-W | 67,000 | 1.6 | 346 |
| t32.8 hIgG1 | nano-W | 67,000 | 1.6 | 193 |
| o42.1 hIgG1 | UF | 67,000 | 1.6 | 525 |
| i52.3 hIgG1 | nano-W | 52,000 | 2.07 | 391 |
| i52.6 hIgG1 | nano-W | 52,000 | 2.07 | 282 |
| D3-08 Fc | nano-W | 57,000 | 2.52 | 88 |
| D3-36 Fc | nano-W | 57,000 | 2.52 | 93 |

**Table S4.** Details on EM data processing of different AbCs.

| Sample name | Particle picking | CTF estimation | 2D class averages | Ab initio reconstruction and symmetry applied | 3D refinement and symmetry applied | # particle in final 3D map/total picked particles | Estimated resolution of 3D map (Å) (*) |
|---|---|---|---|---|---|---|---|
| d2.3 Fc | cisTEM | cisTEM | cisTEM | cisTEM, D2 | cisTEM, D2 | 8295/11211 | - |
| d2.4 Fc | DoG picker | GCTF | cryoSPARC | cryoSPARC, C1 | cryoSPARC, C1 | 28562/46306 | - |
| d2.7 Fc | cisTEM | cisTEM | cisTEM | cisTEM, D2 | cisTEM, D2 | 17002/24441 | - |
| t32.4 Fc | cisTEM | cisTEM | cisTEM | cisTEM, T2 | cisTEM, T2 | 12416/16806 | - |
| t32.8 Fc | cisTEM | cisTEM | cisTEM | cisTEM, T2 | cisTEM, T2 | 7638/16147 | - |
| o42.1 Fc (cryo-EM) | cryoSPARC Template picking | CTFFIND4 within cryoSPARC | cryoSPARC | cryoSPARC, C1 | cryoSPARC, O | 4032/16611 | 11.14 |
| i52.3 Fc (cryo-EM) | cryoSPARC Template picking | CTFFIND4 within cryoSPARC | cryoSPARC | cryoSPARC, C1 | cryoSPARC, I | 3918/11076 | 12.18 |
| i52.6 Fc | cisTEM | cisTEM | cisTEM | cisTEM, I2 | cisTEM, I2 | 11801/26436 | - |
| d2.3 hIgG1 | cisTEM | cisTEM | cisTEM | - | - | - | - |
| d2.4 hIgG1 | cisTEM | cisTEM | cisTEM | - | - | - | - |
| d2.7 hIgG1 | cisTEM | cisTEM | cisTEM | - | - | - | - |
| t32.4 hIgG1 | cisTEM | cisTEM | cisTEM | - | - | - | - |
| t32.8 hIgG1 | cisTEM | cisTEM | cisTEM | - | - | - | - |
| o42.1 hIgG1 | DoG picker | GCTF | cryoSPARC | - | - | - | - |
| i52.3 hIgG1 | cisTEM | cisTEM | cisTEM | - | - | - | - |

| | | | | | | | |
|---|---|---|---|---|---|---|---|
| i52.6 hIgG1 | cisTEM | cisTEM | Relion | - | - | - | - |
| D3-08 Fc | cisTEM | cisTEM | cisTEM | cisTEM, D3 | cisTEM, D3 | 12322/80375 | - |
| D3-36 Fc | cisTEM | cisTEM | cisTEM | cisTEM, D3 | cisTEM, D3 | 16947/43301 | - |

(*) Negative stain reconstructions obtained had resolution of ~20Å.

**Table S5. Success rates of designed antibody-binding cage-forming oligomers using unvalidated building blocks** (see table S1 for descriptions of columns 3+4). Data for cage formation with Fc are shown in Fig. S5.

| **Geometry** | **# ordered** | **Soluble component** | **Good SEC component** | **Forms cage with Fc** |
|---|---|---|---|---|
| D3 dihedron | 14 | 9 | 3 | 2 |
| T32 tetrahedron | 11 | 10 | 2 | 0 |
| O32 octahedron | 15 | 12 | 3 | 0 |
| I32 icosahedron | 8 | 5 | 2 | 0 |
| Total | 48 | 36 | 10 | 2 |

**Table S6.** Details on dynamic light scattering data. Predicted diameters are estimated from computational models fit with appropriate ligands. Given the difficult-to-assess flexibility associated with Fc-fusions to either functional ligands (A1F) or Fab domains, the estimates of these cages may not be accurate.

| AbC description | Figure | Predicted diameter from design (nm) | Hydrodynamic diameter (nm) | Standard deviation (nm) |
|---|---|---|---|---|
| d2.3 Fc week 1 | S5a | 14.19 | 12.87 | 6.70 |
| d2.3 Fc week 2 | S5a | 14.19 | 13.75 | 7.72 |
| d2.3 Fc week 3 | S5a | 14.19 | 11.96 | 5.93 |
| d2.4 Fc week 1 | S5a | 14.30 | 13.08 | 5.79 |
| d2.4 Fc week 2 | S5a | 14.30 | 13.10 | 6.13 |
| d2.4 Fc week 3 | S5a | 14.30 | 12.87 | 5.58 |
| d2.7 Fc week 1 | S5a | 14.96 | 13.73 | 4.77 |
| d2.7 Fc week 2 | S5a | 14.96 | 13.65 | 4.74 |
| d2.7 Fc week 3 | S5a | 14.96 | 13.45 | 5.23 |
| d2.7 Fc week 4 | S5a | 14.96 | 13.89 | 5.14 |
| t32.4 Fc week 1 | S5a | 27.96 | 24.03 | 6.44 |
| t32.4 Fc week 2 | S5a | 27.96 | 24.79 | 6.98 |
| t32.4 Fc week 3 | S5a | 27.96 | 24.30 | 6.98 |
| t32.4 Fc week 4 | S5a | 27.96 | 25.77 | 7.49 |
| t32.8 Fc week 1 | S5a | 25.31 | 24.16 | 6.59 |
| t32.8 Fc week 2 | S5a | 25.31 | 24.68 | 8.68 |
| t32.8 Fc week 3 | S5a | 25.31 | 25.71 | 6.47 |
| t32.8 Fc week 4 | S5a | 25.31 | 25.80 | 10.46 |
| o42.1 Fc week 1 | S5a | 31.50 | 30.65 | 5.71 |
| o42.1 Fc week 2 | S5a | 31.50 | 30.98 | 7.01 |
| o42.1 Fc week 3 | S5a | 31.50 | 30.77 | 3.40 |

| | | | | |
|---|---|---|---|---|
| o42.1 Fc week 4 | S5a | 31.50 | 31.52 | 9.06 |
| i52.3 Fc week 1 | S5a | 42.12 | 43.84 | 9.61 |
| i52.3 Fc week 2 | S5a | 42.12 | 42.75 | 8.96 |
| i52.3 Fc week 3 | S5a | 42.12 | 43.00 | 5.49 |
| i52.3 Fc week 4 | S5a | 42.12 | 43.85 | 4.98 |
| i52.6 Fc week 1 | S5a | 44.96 | 51.33 | 9.19 |
| i52.6 Fc week 2 | S5a | 44.96 | 50.32 | 9.83 |
| i52.6 Fc week 3 | S5a | 44.96 | 50.53 | 13.88 |
| i52.6 Fc week 4 | S5a | 44.96 | 50.37 | 14.21 |
| o42.1 A1F-Fc | S8b | 38.25 | 39.82 | 25.24 |
| i52.3 A1F-Fc | S8b | 49.43 | 58.49 | 32.17 |
| o42.1 LOB7/6 (α-CD40) | S9b | 40.40 | 79.52 | 20.21 |
| o42.1 α-CD3/28 | S9d | 40.40 | 36.66 | 3.66 |
| o42.1 CV1 (α-CoV-2 S) | S10b | 40.40 | 44.76 | 23.67 |

**Table S7. Statistical information for exchange experiments.**

| Experiment (Fig.) | Condition | n | Test | Mean compared to | α | Summary | Adjusted P value |
|---|---|---|---|---|---|---|---|
| Cage fraction 485 ex./ 510 em. | o42.1 GFP-Fc | 3 | 1way ANOVA with post-hoc Dunnett | N/A | N/A | N/A | N/A |
| | RFP-Fc | 3 | 1way ANOVA with post-hoc Dunnett | o42.1 GFP-Fc | 0.05 | **** | <0.0001 |
| | o42.1 pre-exchanged | 3 | 1way ANOVA with post-hoc Dunnett | o42.1 GFP-Fc | 0.05 | **** | <0.0001 |
| | 5 min o42.1 GFP-Fc + 25x RFP-Fc | 3 | 1way ANOVA with post-hoc Dunnett | o42.1 GFP-Fc | 0.05 | **** | <0.0001 |
| | 2 hr o42.1 GFP-Fc + 25x RFP-Fc | 3 | 1way ANOVA with post-hoc Dunnett | o42.1 GFP-Fc | 0.05 | ns | 0.9891 |
| | 4 hr o42.1 GFP-Fc + 25x RFP-Fc | 3 | 1way ANOVA with post-hoc Dunnett | o42.1 GFP-Fc | 0.05 | ns | 0.5186 |
| | 24 hr o42.1 GFP-Fc + 25x RFP-Fc | 3 | 1way ANOVA with post-hoc Dunnett | o42.1 GFP-Fc | 0.05 | ns | 0.0773 |
| Cage fraction 558 ex./ 605 em. | o42.1 GFP-Fc | 3 | 1way ANOVA with post-hoc Dunnett | N/A | N/A | N/A | N/A |
| | RFP-Fc | 3 | 1way ANOVA with post-hoc Dunnett | o42.1 GFP-Fc | 0.05 | ns | 0.7647 |
| | o42.1 pre-exchanged | 3 | 1way ANOVA with post-hoc Dunnett | o42.1 GFP-Fc | 0.05 | **** | <0.0001 |
| | 5 min o42.1 GFP-Fc + 25x RFP-Fc | 3 | 1way ANOVA with post-hoc Dunnett | o42.1 GFP-Fc | 0.05 | *** | 0.0004 |
| | 2 hr o42.1 GFP-Fc + 25x RFP-Fc | 3 | 1way ANOVA with post-hoc Dunnett | o42.1 GFP-Fc | 0.05 | **** | <0.0001 |

| | | | | | | | |
|---|---|---|---|---|---|---|---|
| | 4 hr o42.1 GFP-Fc + 25x RFP-Fc | 3 | 1way ANOVA with post-hoc Dunnett | o42.1 GFP-Fc | 0.05 | **** | <0.0001 |
| | 24 hr o42.1 GFP-Fc + 25x RFP-Fc | 3 | 1way ANOVA with post-hoc Dunnett | o42.1 GFP-Fc | 0.05 | **** | <0.0001 |

**Table S8. Statistical information for DR5 experiments.** All analyses were performed using Graphpad Prism Software.

| Experiment (Fig.) | Condition | Concentration | n | Test | Mean compared to | α | Summary | Adjusted P value |
|---|---|---|---|---|---|---|---|---|
| Caspase-3,7 RCC4 (4b) | PBS | 15 pM | 6 | 2way ANOVA with post-hoc Dunnett | N/A | N/A | N/A | N/A |
| | | 1.5 nM | 6 | 2way ANOVA with post-hoc Dunnett | 15 pM PBS | 0.05 | ns | 0.9996 |
| | | 150 nM | 6 | 2way ANOVA with post-hoc Dunnett | 15 pM PBS | 0.05 | ns | 0.9994 |
| | TRAIL | 15 pM | 6 | 2way ANOVA with post-hoc Dunnett | 15 pM PBS | 0.05 | ns | 0.9996 |
| | | 1.5 nM | 6 | 2way ANOVA with post-hoc Dunnett | 15 pM PBS | 0.05 | ns | 0.9996 |
| | | 150 nM | 6 | 2way ANOVA with post-hoc Dunnett | 15 pM PBS | 0.05 | ns | 0.9668 |
| | α-DR5 | 15 pM | 6 | 2way ANOVA with post-hoc Dunnett | 15 pM PBS | 0.05 | ns | 0.9997 |
| | | 1.5 nM | 6 | 2way ANOVA with post-hoc Dunnett | 15 pM PBS | 0.05 | ns | >0.9999 |
| | | 150 nM | 6 | 2way ANOVA with post-hoc Dunnett | 15 pM PBS | 0.05 | ns | 0.2005 |
| | d2.4 α-DR5 | 15 pM | 6 | 2way ANOVA with post-hoc Dunnett | 15 pM PBS | 0.05 | ns | 0.9995 |

| | | | | | | | |
|---|---|---|---|---|---|---|---|
| | 1.5 nM | 6 | 2way ANOVA with post-hoc Dunnett | 15 pM PBS | 0.05 | ns | 0.9988 |
| | 150 nM | 6 | 2way ANOVA with post-hoc Dunnett | 15 pM PBS | 0.05 | **** | <0.0001 |
| t32.4 α-DR5 | 15 pM | 6 | 2way ANOVA with post-hoc Dunnett | 15 pM PBS | 0.05 | ns | 0.9998 |
| | 1.5 nM | 6 | 2way ANOVA with post-hoc Dunnett | 15 pM PBS | 0.05 | **** | <0.0001 |
| | 150 nM | 6 | 2way ANOVA with post-hoc Dunnett | 15 pM PBS | 0.05 | **** | <0.0001 |
| t32.8 α-DR5 | 15 pM | 3 | 2way ANOVA with post-hoc Dunnett | 15 pM PBS | 0.05 | ns | 0.984 |
| | 1.5 nM | 3 | 2way ANOVA with post-hoc Dunnett | 15 pM PBS | 0.05 | ns | 0.6177 |
| | 150 nM | 3 | 2way ANOVA with post-hoc Dunnett | 15 pM PBS | 0.05 | **** | <0.0001 |
| o42.1 α-DR5 | 15 pM | 6 | 2way ANOVA with post-hoc Dunnett | 15 pM PBS | 0.05 | ns | 0.9991 |
| | 1.5 nM | 6 | 2way ANOVA with post-hoc Dunnett | 15 pM PBS | 0.05 | **** | <0.0001 |
| | 150 nM | 6 | 2way ANOVA with post-hoc Dunnett | 15 pM PBS | 0.05 | **** | <0.0001 |

| | | | | | | | | |
|---|---|---|---|---|---|---|---|---|
| | i52.3 α-DR5 | 15 pM | 3 | 2way ANOVA with post-hoc Dunnett | 15 pM PBS | 0.05 | ns | 0.9442 |
| | i52.3 α-DR5 | 1.5 nM | 3 | 2way ANOVA with post-hoc Dunnett | 15 pM PBS | 0.05 | ns | 0.8227 |
| | i52.3 α-DR5 | 150 nM | 3 | 2way ANOVA with post-hoc Dunnett | 15 pM PBS | 0.05 | **** | <0.0001 |
| Viability 4d RCC4 (4c) | PBS | 150 nM | 3 | 1way ANOVA with post-hoc Dunnett | N/A | N/A | N/A | N/A |
| | TRAIL | 150 nM | 3 | 1way ANOVA with post-hoc Dunnett | PBS | 0.05 | ns | 0.5207 |
| | α-DR5 | 150 nM | 3 | 1way ANOVA with post-hoc Dunnett | PBS | 0.05 | ns | 0.9996 |
| | d2.4 α-DR5 | 150 nM | 3 | 1way ANOVA with post-hoc Dunnett | PBS | 0.05 | * | 0.0238 |
| | t32.4 α-DR5 | 150 nM | 3 | 1way ANOVA with post-hoc Dunnett | PBS | 0.05 | **** | <0.0001 |
| | t32.8 α-DR5 | 150 nM | 3 | 1way ANOVA with post-hoc Dunnett | PBS | 0.05 | **** | <0.0001 |
| | o42.1 α-DR5 | 150 nM | 3 | 1way ANOVA with post-hoc Dunnett | PBS | 0.05 | **** | <0.0001 |
| | i52.3 α-DR5 | 150 nM | 3 | 1way ANOVA with post-hoc Dunnett | PBS | 0.05 | ** | 0.0079 |

| | | | | | | | | |
|---|---|---|---|---|---|---|---|---|
| | PBS | 150 nM | 3 | 1way ANOVA with post-hoc Dunnett | N/A | N/A | N/A | N/A |
| Viability 4d RCC4 Fc cages (4d) | d2.4 Fc | 150 nM | 3 | 1way ANOVA with post-hoc Dunnett | PBS | 0.05 | ns | 0.7157 |
| | t32.4 Fc | 150 nM | 3 | 1way ANOVA with post-hoc Dunnett | PBS | 0.05 | ns | 0.9976 |
| | t32.8 Fc | 150 nM | 3 | 1way ANOVA with post-hoc Dunnett | PBS | 0.05 | ns | 0.8556 |
| | o42.1 Fc | 150 nM | 3 | 1way ANOVA with post-hoc Dunnett | PBS | 0.05 | ns | 0.2309 |
| | i52.3 Fc | 150 nM | 3 | 1way ANOVA with post-hoc Dunnett | PBS | 0.05 | ns | 0.9302 |
| | PBS | 150 nM | 6 | 1way ANOVA with post-hoc Dunnett | N/A | N/A | N/A | N/A |
| Viability 6d RCC4 (4e) | TRAIL | 150 nM | 6 | 1way ANOVA with post-hoc Dunnett | PBS | 0.05 | ns | 0.9996 |
| | α-DR5 | 150 nM | 6 | 1way ANOVA with post-hoc Dunnett | PBS | 0.05 | ns | >0.9999 |
| | t32.4 Fc | 150 nM | 3 | 1way ANOVA with post-hoc Dunnett | PBS | 0.05 | ns | 0.9591 |
| | o42.1 Fc | 150 nM | 3 | 1way ANOVA with post-hoc Dunnett | PBS | 0.05 | ns | 0.9593 |

| | | | | | | | | |
|---|---|---|---|---|---|---|---|---|
| | t32.4 α-DR5 | 150 nM | 6 | 1way ANOVA with post-hoc Dunnett | PBS | 0.05 | **** | <0.0001 |
| | o42.1 α-DR5 | 150 nM | 6 | 1way ANOVA with post-hoc Dunnett | PBS | 0.05 | *** | 0.0001 |
| c-PARP quant. (4g) | PBS | 150 nM | 4 | 1way ANOVA with post-hoc Dunnett | N/A | N/A | N/A | N/A |
| | TRAIL | 150 nM | 4 | 1way ANOVA with post-hoc Dunnett | PBS | 0.05 | ns | 0.0845 |
| | α-DR5 | 150 nM | 3 | 1way ANOVA with post-hoc Dunnett | PBS | 0.05 | ns | 0.4746 |
| | o42.1 Fc | 150 nM | 3 | 1way ANOVA with post-hoc Dunnett | PBS | 0.05 | ns | 0.9979 |
| | o42.1 α-DR5 | 150 nM | 3 | 1way ANOVA with post-hoc Dunnett | PBS | 0.05 | **** | <0.0001 |
| Caspase-3,7 Colo205 (S7a) | PBS | 15 pM | 2 | 2way ANOVA with post-hoc Dunnett | N/A | N/A | N/A | N/A |
| | | 1.5 nM | 2 | 2way ANOVA with post-hoc Dunnett | 15 pM PBS | 0.05 | ns | >0.9999 |
| | | 150 nM | 2 | 2way ANOVA with post-hoc Dunnett | 15 pM PBS | 0.05 | ns | >0.9999 |
| | TRAIL | 15 pM | 2 | 2way ANOVA with post-hoc Dunnett | 15 pM PBS | 0.05 | ns | >0.9999 |

| | | | | | | | |
|---|---|---|---|---|---|---|---|
| | 1.5 nM | 2 | 2way ANOVA with post-hoc Dunnett | 15 pM PBS | 0.05 | *** | 0.0006 |
| | 150 nM | 2 | 2way ANOVA with post-hoc Dunnett | 15 pM PBS | 0.05 | **** | <0.0001 |
| α-DR5 | 15 pM | 2 | 2way ANOVA with post-hoc Dunnett | 15 pM PBS | 0.05 | ns | 0.9997 |
| | 1.5 nM | 2 | 2way ANOVA with post-hoc Dunnett | 15 pM PBS | 0.05 | ns | >0.9999 |
| | 150 nM | 2 | 2way ANOVA with post-hoc Dunnett | 15 pM PBS | 0.05 | ns | 0.9996 |
| d2.4 α-DR5 | 15 pM | 2 | 2way ANOVA with post-hoc Dunnett | 15 pM PBS | 0.05 | ns | 0.9996 |
| | 1.5 nM | 2 | 2way ANOVA with post-hoc Dunnett | 15 pM PBS | 0.05 | **** | <0.0001 |
| | 150 nM | 2 | 2way ANOVA with post-hoc Dunnett | 15 pM PBS | 0.05 | **** | <0.0001 |
| t32.4 α-DR5 | 15 pM | 2 | 2way ANOVA with post-hoc Dunnett | 15 pM PBS | 0.05 | ns | 0.1074 |
| | 1.5 nM | 2 | 2way ANOVA with post-hoc Dunnett | 15 pM PBS | 0.05 | **** | <0.0001 |
| | 150 nM | 2 | 2way ANOVA with post-hoc Dunnett | 15 pM PBS | 0.05 | **** | <0.0001 |

| | | | | | | | | |
|---|---|---|---|---|---|---|---|---|
| | t32.8 α-DR5 | 15 pM | 2 | 2way ANOVA with post-hoc Dunnett | 15 pM PBS | 0.05 | ns | >0.9999 |
| | | 1.5 nM | 2 | 2way ANOVA with post-hoc Dunnett | 15 pM PBS | 0.05 | **** | <0.0001 |
| | | 150 nM | 2 | 2way ANOVA with post-hoc Dunnett | 15 pM PBS | 0.05 | **** | <0.0001 |
| | o42.1 α-DR5 | 15 pM | 2 | 2way ANOVA with post-hoc Dunnett | 15 pM PBS | 0.05 | ns | 0.6538 |
| | | 1.5 nM | 2 | 2way ANOVA with post-hoc Dunnett | 15 pM PBS | 0.05 | **** | <0.0001 |
| | | 150 nM | 2 | 2way ANOVA with post-hoc Dunnett | 15 pM PBS | 0.05 | **** | <0.0001 |
| | i52.3 α-DR5 | 15 pM | 2 | 2way ANOVA with post-hoc Dunnett | 15 pM PBS | 0.05 | ns | >0.9999 |
| | | 1.5 nM | 2 | 2way ANOVA with post-hoc Dunnett | 15 pM PBS | 0.05 | **** | <0.0001 |
| | | 150 nM | 2 | 2way ANOVA with post-hoc Dunnett | 15 pM PBS | 0.05 | **** | <0.0001 |
| Caspase-8 RCC4 12 h (S7b) | PBS | 1.5 nM | 6 | 2way ANOVA with post-hoc Dunnett | 1.5 nM PBS | 0.05 | N/A | N/A |
| | | 150 nM | 6 | 2way ANOVA with post-hoc Dunnett | 1.5 nM PBS | 0.05 | ns | 0.9909 |

| | | | | | | | |
|---|---|---|---|---|---|---|---|
| TRAIL | 1.5 nM | 6 | 2way ANOVA with post-hoc Dunnett | 1.5 nM PBS | 0.05 | ns | 0.999 |
| | 150 nM | 6 | 2way ANOVA with post-hoc Dunnett | 1.5 nM PBS | 0.05 | **** | <0.0001 |
| α-DR5 | 1.5 nM | 6 | 2way ANOVA with post-hoc Dunnett | 1.5 nM PBS | 0.05 | ns | 0.9918 |
| | 150 nM | 6 | 2way ANOVA with post-hoc Dunnett | 1.5 nM PBS | 0.05 | ns | 0.9955 |
| t32.4 Fc | 1.5 nM | 6 | 2way ANOVA with post-hoc Dunnett | 1.5 nM PBS | 0.05 | ns | >0.9999 |
| | 150 nM | 6 | 2way ANOVA with post-hoc Dunnett | 1.5 nM PBS | 0.05 | ns | 0.9048 |
| o42.1 Fc | 1.5 nM | 6 | 2way ANOVA with post-hoc Dunnett | 1.5 nM PBS | 0.05 | ns | >0.9999 |
| | 150 nM | 6 | 2way ANOVA with post-hoc Dunnett | 1.5 nM PBS | 0.05 | ns | 0.9997 |
| t32.4 α-DR5 | 1.5 nM | 6 | 2way ANOVA with post-hoc Dunnett | 1.5 nM PBS | 0.05 | **** | <0.0001 |
| | 150 nM | 6 | 2way ANOVA with post-hoc Dunnett | 1.5 nM PBS | 0.05 | **** | <0.0001 |
| o42.1 α-DR5 | 1.5 nM | 6 | 2way ANOVA with post-hoc Dunnett | 1.5 nM PBS | 0.05 | **** | <0.0001 |

| | | | | | | | | |
|---|---|---|---|---|---|---|---|---|
| | | 150 nM | 6 | 2way ANOVA with post-hoc Dunnett | 1.5 nM PBS | 0.05 | **** | <0.0001 |
| Caspase-3,7 RCC4 Fc cages (S7d) | PBS | 150 nM | 3 | 1way ANOVA with post-hoc Dunnett | N/A | N/A | N/A | N/A |
| | d2.4 Fc | 150 nM | 6 | 1way ANOVA with post-hoc Dunnett | PBS | 0.05 | * | 0.0129 |
| | t32.4 Fc | 150 nM | 6 | 1way ANOVA with post-hoc Dunnett | PBS | 0.05 | * | 0.046 |
| | t32.8 Fc | 150 nM | 3 | 1way ANOVA with post-hoc Dunnett | PBS | 0.05 | ns | 0.9198 |
| | o42.1 Fc | 150 nM | 6 | 1way ANOVA with post-hoc Dunnett | PBS | 0.05 | ns | 0.2112 |
| | i52.3 Fc | 150 nM | 3 | 1way ANOVA with post-hoc Dunnett | PBS | 0.05 | ns | 0.9996 |
| RAM009 Caspase-3,7 (S7f) | PBS | 1.5 nM | 6 | 2way ANOVA with post-hoc Dunnett | PBS | 0.05 | N/A | N/A |
| | | 150 nM | 6 | 2way ANOVA with post-hoc Dunnett | PBS | 0.05 | ns | 0.3848 |
| | TRAIL | 1.5 nM | 6 | 2way ANOVA with post-hoc Dunnett | PBS | 0.05 | ns | 0.9726 |
| | | 150 nM | 6 | 2way ANOVA with post-hoc Dunnett | PBS | 0.05 | ns | 0.0525 |

| | | | | | | | |
|---|---|---|---|---|---|---|---|
| α-DR5 | 1.5 nM | 6 | 2way ANOVA with post-hoc Dunnett | PBS | 0.05 | ns | 0.9566 |
| | 150 nM | 6 | 2way ANOVA with post-hoc Dunnett | PBS | 0.05 | ns | 0.2752 |
| d2.4 α-DR5 | 1.5 nM | 6 | 2way ANOVA with post-hoc Dunnett | PBS | 0.05 | ns | 0.9677 |
| | 150 nM | 6 | 2way ANOVA with post-hoc Dunnett | PBS | 0.05 | ** | 0.0076 |
| t32.4 α-DR5 | 1.5 nM | 6 | 2way ANOVA with post-hoc Dunnett | PBS | 0.05 | ns | 0.9703 |
| | 150 nM | 6 | 2way ANOVA with post-hoc Dunnett | PBS | 0.05 | ** | 0.0028 |
| t32.8 α-DR5 | 1.5 nM | 6 | 2way ANOVA with post-hoc Dunnett | PBS | 0.05 | ns | 0.9996 |
| | 150 nM | 6 | 2way ANOVA with post-hoc Dunnett | PBS | 0.05 | ** | 0.0067 |
| o42.1 α-DR5 | 1.5 nM | 6 | 2way ANOVA with post-hoc Dunnett | PBS | 0.05 | ns | 0.9991 |
| | 150 nM | 6 | 2way ANOVA with post-hoc Dunnett | PBS | 0.05 | ** | 0.0038 |
| i52.3 α-DR5 | 1.5 nM | 6 | 2way ANOVA with post-hoc Dunnett | PBS | 0.05 | ns | 0.9994 |

| | | | | | | | | |
|---|---|---|---|---|---|---|---|---|
| | | 150 nM | 6 | 2way ANOVA with post-hoc Dunnett | PBS | 0.05 | *** | 0.0006 |
| | d2.4 Fc | 150 nM | 6 | 2way ANOVA with post-hoc Dunnett | PBS | 0.05 | ns | 0.9966 |
| | t32.4 Fc | 150 nM | 6 | 2way ANOVA with post-hoc Dunnett | PBS | 0.05 | ns | 0.9997 |
| | t32.8 Fc | 150 nM | 6 | 2way ANOVA with post-hoc Dunnett | PBS | 0.05 | ns | 0.9992 |
| | o42.1 Fc | 150 nM | 6 | 2way ANOVA with post-hoc Dunnett | PBS | 0.05 | ns | 0.9994 |
| | i52.3 Fc | 150 nM | 6 | 2way ANOVA with post-hoc Dunnett | PBS | 0.05 | ns | 0.9995 |
| RAM009 Viability (S7g) | PBS | 150 nM | 6 | 2way ANOVA with post-hoc Dunnett | PBS | 0.05 | | |
| | TRAIL | 150 nM | 3 | 2way ANOVA with post-hoc Dunnett | PBS | 0.05 | ns | 0.9901 |
| | α-DR5 | 150 nM | 3 | 2way ANOVA with post-hoc Dunnett | PBS | 0.05 | ns | 0.9995 |
| | d2.4 α-DR5 | 150 nM | 3 | 2way ANOVA with post-hoc Dunnett | PBS | 0.05 | ns | 0.9996 |
| | t32.4 α-DR5 | 150 nM | 3 | 2way ANOVA with post-hoc Dunnett | PBS | 0.05 | ns | 0.9212 |

| | | | | | | | | |
|---|---|---|---|---|---|---|---|---|
| | t32.8 α-DR5 | 150 nM | 3 | 2way ANOVA with post-hoc Dunnett | PBS | 0.05 | ns | 0.7875 |
| | o42.1 α-DR5 | 150 nM | 3 | 2way ANOVA with post-hoc Dunnett | PBS | 0.05 | ns | 0.7485 |
| | i52.3 α-DR5 | 150 nM | 3 | 2way ANOVA with post-hoc Dunnett | PBS | 0.05 | ns | 0.1419 |
| | d2.4 Fc | 150 nM | 3 | 2way ANOVA with post-hoc Dunnett | PBS | 0.05 | ns | 0.9996 |
| | t32.4 Fc | 150 nM | 3 | 2way ANOVA with post-hoc Dunnett | PBS | 0.05 | ns | 0.9718 |
| | t32.8 Fc | 150 nM | 3 | 2way ANOVA with post-hoc Dunnett | PBS | 0.05 | ns | 0.999 |
| | o42.1 Fc | 150 nM | 3 | 2way ANOVA with post-hoc Dunnett | PBS | 0.05 | ns | 0.9913 |
| | i52.3 Fc | 150 nM | 3 | 2way ANOVA with post-hoc Dunnett | PBS | 0.05 | ns | 0.9837 |

**Table S9. Statistical information for A1F-Fc experiments.** All analyses were performed using Graphpad Prism Software.

| Experiment (Fig.) | Condition | n | Test | Mean compared to | α | Summary | Adjusted P value |
|---|---|---|---|---|---|---|---|
| pAKT (4j, S8c (H8)) | PBS | 13 | 1way ANOVA with post-hoc Dunnett | N/A | N/A | N/A | N/A |
| | A1F-Fc | 3 | 1way ANOVA with post-hoc Dunnett | PBS | 0.05 | ns | >0.9999 |
| | o42.1 Fc | 4 | 1way ANOVA with post-hoc Dunnett | PBS | 0.05 | ns | >0.9999 |
| | i52.3 Fc | 3 | 1way ANOVA with post-hoc Dunnett | PBS | 0.05 | ns | >0.9999 |
| | o42.1 A1F-Fc | 9 | 1way ANOVA with post-hoc Dunnett | PBS | 0.05 | **** | <0.0001 |
| | i52.3 A1F-Fc | 8 | 1way ANOVA with post-hoc Dunnett | PBS | 0.05 | **** | <0.0001 |
| | H8-A1F | 4 | 1way ANOVA with post-hoc Dunnett | PBS | 0.05 | **** | <0.0001 |
| pERK1-2 (4j; S8c (H8)) | PBS | 13 | 1way ANOVA with post-hoc Dunnett | N/A | N/A | N/A | N/A |
| | A1F-Fc | 3 | 1way ANOVA with post-hoc Dunnett | PBS | 0.05 | ns | 0.9997 |
| | o42.1 Fc | 4 | 1way ANOVA with post-hoc Dunnett | PBS | 0.05 | ns | 0.9955 |
| | i52.3 Fc | 3 | 1way ANOVA with post-hoc Dunnett | PBS | 0.05 | ns | 0.9997 |
| | o42.1 A1F-Fc | 9 | 1way ANOVA with post-hoc Dunnett | PBS | 0.05 | ** | 0.0015 |
| | i52.3 A1F-Fc | 8 | 1way ANOVA with post-hoc Dunnett | PBS | 0.05 | *** | 0.0004 |
| | H8-A1F | 6 | 1way ANOVA with post-hoc Dunnett | PBS | 0.05 | ns | 0.4766 |
| Vascular stability (4k; S8c (H8)) | PBS | 7 | 1way ANOVA with post-hoc Dunnett | N/A | N/A | N/A | N/A |
| | A1F-Fc | 6 | 1way ANOVA with post-hoc Dunnett | PBS | 0.05 | ns | 0.9998 |
| | o42.1 Fc | 4 | 1way ANOVA with post-hoc Dunnett | PBS | 0.05 | ns | 0.9946 |

| | | | | | | | |
|---|---|---|---|---|---|---|---|
| | i52.3 Fc | 3 | 1way ANOVA with post-hoc Dunnett | PBS | 0.05 | ns | 0.6157 |
| | o42.1 A1F-Fc | 6 | 1way ANOVA with post-hoc Dunnett | PBS | 0.05 | *** | 0.0004 |
| | i52.3 A1F-Fc | 5 | 1way ANOVA with post-hoc Dunnett | PBS | 0.05 | **** | <0.0001 |
| | H8-A1F | 4 | 1way ANOVA with post-hoc Dunnett | PBS | 0.05 | * | 0.011 |
| pAKT (10% HS exp.; S8e) | PBS | 3 | 1way ANOVA with post-hoc Dunnett | o42.1 A1F-Fc | 0.05 | *** | <0.0001 |
| | o42.1 A1F-Fc | 3 | 1way ANOVA with post-hoc Dunnett | o42.1 A1F-Fc | 0.05 | N/A | N/A |
| | 10% HS | 3 | 1way ANOVA with post-hoc Dunnett | o42.1 A1F-Fc | 0.05 | *** | 0.0002 |
| | o42.1 A1F-Fc 10% HS 4°C | 3 | 1way ANOVA with post-hoc Dunnett | o42.1 A1F-Fc | 0.05 | ns | 0.9431 |
| | o42.1 A1F-Fc 10% HS 37°C | 3 | 1way ANOVA with post-hoc Dunnett | o42.1 A1F-Fc | 0.05 | ns | 0.9998 |

**Table S10. EC50s from CD40 activation experiments.** EC50 values were interpolated from the response curves determined using the log(agonist) vs. response -- Variable slope (four parameters) fit using Graphpad Prism Software

| | IC50 log(uM) | 95% CI log(uM) |
|---|---|---|
| o42.1 IgG control | ~ -1.455 | (Very wide) |
| α-CD40 | 1.466 | 1.247 to 1.833 |
| LOB7/6 | ~ -1.650 | (Very wide) |
| o42.1 LOB7/6 | 0.1134 | -0.001058 to ??? |

**Table S11. IC50s from SARS-CoV neutralization experiments.** IC50 values were interpolated from the neutralization curves determined using the log(inhibitor) vs. response -- Variable slope (four parameters) fit using Graphpad Prism Software

| **SARS-CoV-2 pseudovirus neutralization** | | |
|---|---|---|
| | **IC50 log(uM)** | **95% CI log(uM)** |
| CV1 | -1.201 | -1.935 to ??? |
| o42.1 CV1 | -3.566 | ??? to -3.044 |
| CV30 | -3.301 | -3.464 to -3.141 |
| o42.1 CV30 | -3.716 | -3.914 to -3.556 |
| CV3 | ~ -2.591 | (Very wide) |
| o42.1 CV3 | -4.449 | ??? |
| o42.1 Fc | -2.068 | ??? |
| Fc-ACE2 | -1.768 | -2.016 to -1.483 |
| o42.1 Fc-ACE2 | -2.655 | -2.740 to -2.589 |
| **SARS-CoV-1 pseudovirus neutralization** | | |
| | **IC50 log(uM)** | **95% CI log(uM)** |
| o42.1 Fc | ~ 1.631e+015 | ~ 1.631e+015 |
| Fc-ACE2 | -1.642 | -1.642 |
| o42.1 Fc-ACE2 | -2.039 | -2.039 |

**Table S12. List of antibodies formed into cages as verified by at minimum size exclusion chromatography.** Successfully formed cages (by SEC) listed by the antibody target reactivity, antibody species and isotype, and designs used

| Ab reactivity | Ab subclass | Designs (validated by SEC at minimum) | Comments |
|---|---|---|---|
| α-CD3 | mIgG2a | t32.4, o42.1 | OKT3 |
| α-CD4 | mIgG2b | o42.1 | OKT4 |
| α-CD28 | hIgG1 | t32.4, o42.1 | CD28.6 |
| α-CD40 | mIgG2a or mIgG2b | o42.1 | LOB7/6 or 82111 (respectively) |
| α-CoV2 S | hIgG1 | o42.1 | CV1, CV3, CV30 |
| α-DR5 (human) | hIgG1 | d2.3, d2.4, d2.7, t32.4, t32.8, o42.1, i52.3, i52.6 | conatumumab |
| α-DR5 (mouse) | Armenian hamster IgG | t32.4, o42.1 | MD5-1 |
| α-EGFR | hIgG1 | mIgG2b | cetuximab |
| α-LRP6 | hIgG1 | t32.4, o42.1 | YW210.09 |
| α-RSV F | hIgG1 | d2.3, d2.4, d2.7, t32.4, t32.8, o42.1, i52.3, i52.6 | mpe8 |
| Non-specific | Rabbit IgG | d2.4, o42.1 | Rabbit serum IgG |

**Table S13. List of Fc-fusions formed into cages as verified by at minimum size exclusion chromatography.** Successfully formed cages (by SEC) listed by the ligand that was fused to Fc, the Fc sequence species and isotype, and designs used

| Fc-fusion ligand | Fc subclass | Designs (validated by SEC at minimum) | Comments |
|---|---|---|---|
| Angiopoietin-1 F-domain | hIgG1 | d2.4, t32.4, t32.8, o42.1, i52.3 | |
| Angiotensin-converting enzyme 2 (ACE2) | hIgG1 | o42.1 | |
| CD80 | hIgG1 | o42.1 | |
| mRuby2 | hIgG1 | d2.4, t32.4, t32.8, o42.1, i52.3 | |
| sfGFP | hIgG1 | d2.4, t32.4, t32.8, o42.1, i52.3 | |
| VEGF-a | hIgG1 | t32.4, o42.1 | |
| VEGF-c | hIgG1 | t32.4, o42.1 | |

**Table S14.** Amino acid sequences of all successful AbC-forming designs.

| Name | Sequence |
|---|---|
| d2.3 | MSDEEERNELIKRIREAAQRAREAAERTGDPRVRELARELARIAQIAFYLVLHDPSSSEVNEALKAVVKAIELAVRALEEAEK<br>TGDPEVRELAREVVRLAVEVATATAAGENDTLRKVAERALRLAKEAAKRGDAKAAKQAAKIAKLAAANAGDEDVLKKVELV<br>RLAIELVEIVVENAKRKGDDDKEAAEAALAAFRIVLAAAQLAGIASLEVLELALRLIKEVVENAQREGYDIAVAAIAAAVAFAVV<br>AVAAAAADITSSEVLELAIRLIKEVVENAQREGYVILLAALAAAAAFVVVAAAAKRAGITSSETLKRAIEEIRKRVEEAQREGND<br>ISEAARQAAEEFRKKAEELKGSLEHHHHHH |
| d2.4 | MSDEEERNELIKRIREAAQRAREAAERTGDPRVRELARELARIAQIAFYLVLHDPSSSEVNEALKAVVKAIELAVRALEAAEK<br>TGDPRVRELAREVVKAAVDVAEAAQAGLNDKLREVAEKALRLAKEALKEGDSTAAELAAEIARLAAKLAGDEDVLKKVKLVL<br>EAIKLVKIVVENAKRKGDDSKEAAEAAVAAFLIVLAAAKLAGIASEEVLELAARLIKEVVENAQREGYDIAVAAIAAAVAFAVVA<br>VAAAAADITSSEVLELAIRLIKEVVENAQREGYVILLAALAAAAAFVVVAAAAKRAGITSSETLKRAIEEIRKRVEEAQREGNDI<br>SEAARQAAEEFRKKAEELKGSLEHHHHHH |
| d2.7 | MSDEEERNELIKRIREAAQRAREAAERTGDPRVRELARELAKLAQIAFYLVLHDPSAKEVNLALELIVKAIELAVRALEEAEKT<br>GDPHARELAREIVRLAVELARAVAEAAEEAKKQGNSELAEQVARAAQVALEVIKAAITAAKQGDRKAFRAALELVLEVIKAIE<br>EAVKQGNPKKVAEVALKAELIRIVVQNAANKGDDADEAVEAARAAFEIVLAAAQLAGIDSEEVLELAARLIKEVVENAQREGY<br>DIAVAAIAAAVAFAVVAVAAAAADITSSEVLELAIRLIKEVVENAVREGYVILLAALAAAAAFVVVAAAAKRAGITSSETLKRAIE<br>EIRKRVEEAQREGNDISEAARQAAEEFRKKAEELKGSLEHHHHHH |
| t32.4 | MFNKSQQSAFYLILNMPNLNEAQRNGFIQSLKDDPSKSEVVAGEAAIEAARNALKKGSPETAREAVRLALELVQEAERQAR<br>KTGSTERLIAAAKLAIEVARVALKVGSPETAREAVRTALELVQELIRQARKTGSKEVLEEAAKLALEVAKVAAEVGSPETAAR<br>AVATAVEALKEAGASEDEIAEIVARVISEVIRILKESGSEYKVICRAVARIVAEIVEALKRSGTSEDEIAEIVARVISEVIRTLKES<br>GSDYLIICVCVAIIVAEIVEALKRSGTSEDEIAEIVARVISEVIRTLKESGSSYEVIKECVQIIVLAIILALMKSGTEVEEILLILLRVK<br>TEVRRTLKESGSLEHHHHHH |
| t32.8 | MFNKDQQSAFYEVLNMPNLNEAQRNGFIQSLKDDPSQSLKILIKAAAGGDSELEEVAKRIVKELAEQGRSEKEAAKEAAELI<br>ERITRAAGGNSDLIELAVRIVKILEEQGRSPSEAAKEAVEAIEAIVRAAGGDSEAIKVAAEIAKTIITQKESGSEYKEICRTVARI<br>VAEIVEKLKRNGASEDEIAEIVAAIIAAVILTLKLSGSDYLIICVCVAIIVAEIVEALKRSGTSEDEIAEIVARVISAVIRVLKESGSS<br>YEVIKECVQIIVLAIILALMKSGTEVEEILLILLRVKTEVRRTLKESGSLEHHHHHH |
| o42.1 | MFNKDQQSAFYEILNMPNLNEALRNGFIQLLKDDPSKSTVILTAAKVAAELSEKIRTLKESGSSYEQIAETVAKAVAKLVEKLK<br>RNGVSEDEIALAVALIISAVIQTLKESGSSYEVIAEIVARIVAEIVEALKRSGTSEDEIAEIVARVISEVIRTLKESGSSYEVIAEIV<br>ARIVAEIVEALKRSGTSEDEIAKIVARVIAEVLRTLKESGSSEEVIKEIVARIITEIKEALKRSGTSEDEIELITLMIEAALEIAKLKS<br>SGSEYEEICEDVARRIAELVEKLKRDGTSAVEIAKIVAAIISAVIAMLKASGSSYEVICECVARIVAEIVEALKRSGTSAAIIALIVA<br>LVISEVIRTLKESGSSFEVILECVIRIVLEIIEALKRSGTSEQDVMLIVMAVLLVVLATLQLSGSLEHHHHHH |
| i52.3 | MSDEEERNELIKRIREAAQRAREAAERTGDPRVRELARELARLAQRAFYLVLHDPSSSDVNEALKLIVEAIEAAVRALEAAE<br>RAGDPELREDAREAVRLAVEAAEEVQRNPSSSTANLLLKAIVALAEALAAAANGDKEKFKKAAESALEIAKRVVEVASKEGD<br>PEAVLEAAKVALRVAELAAKNGDKEVFKKAAESALEVAKRLVEVASKEGDPELVLEAAKVALRVAELAAKNGDKEVFQKAA<br>ASAVEVALRLTEVASKEGDSELETEAAKVITRVRELASKQGDAAVAILAETAEVKLEIEESKKRPQSESAKNLILIMQLLINQIR<br>LLVLQIRMLDEQRQEGSLEHHHHHH |
| i52.6 | MSDEEERNELIKRIREAAQRAREAAERTGDPRVRELARELARLAQRAFYLVLHDPSSSDVNEALKLIVEAIEAAVRALEAAE<br>RTGDPKVREEARELVRRAVEAAEEVQRNPSSSEVNEKLKAIVVEIEVKVASLEAKEVTDPDKALKIAKKVIELALEAVKENPS<br>TEALRAVLEAVRLASEVAKRVTDPDKALKIAKLVIELALEAVKEDPSTDALRAVLEAVRLASEVAKRVTDPDKALKIAKLVLEL<br>AAEAVKEDPSTDALRAAKEAERLATEVAKRVTDPKKAREIEMLVLKLQMEAILAETEEVKKEIEESKKRPQSESAKNLILIMQ<br>LLINQIRLLALQIRMLALQLQEGSLEHHHHHH |
| D3-08 | MSDEEERNELIKRIREAAQRAREAAERTGDPRVRELARELARLAQIMFYLVLHDPSAKFVNEALKVVVEMIEMAVRALEKAE<br>RIGDPEMREMARELVRAAVEMADLMTRAAEEARRDPDSSDVNEALKLIREAIEAAKRALEAAERTGDPEVLRLAILLMELAV<br>LAARLVQLDPSASDANEALKKIVEAIERAVRALEKAERTGDPEEREKARQKVAEAVVEAALILAEAALRVAEKAAKNGDKEL<br>FKKAAELALKVARLLVEVASKAGAPEFVLAAAEIAIAVLELAVKQGDRDVALLAAATALFVLVMAARVLFEAGGWLEHHHHH<br>H |
| D3-36 | MFNKDQQSAFYEILNLPKLTEEFRNGFIQALKTAPLASEAILGAAKMAAKATDEEVRRVLLEVVRELARLFTEAERSNDDEC<br>RRLAELAIKAVSLLMKAAEIATDEEEIRRLAEEARELIRLAQEACRSNDDDELTKAAMFVAEMIAKAARETGDDKVLAEALRL<br>EARLIVELAEKACKRGNSEAAERASELAQRVLEKARKVSEEAREQGDDEVLALALIAIALAVLALAEVACCRGNKEEAERAY<br>KDAQRVLLEAILVALKALLQGDEEVARLAQEAAELAQEALDHVQECRGGWLSVLEHHHHHH |

**Table S15.** Amino acid sequences of Fc and Fc-fusions.

| Name | Sequence |
|---|---|
| Fc | METDTLLLWVLLLWVPGSTGHHHHHHGGSENLYFQGGSEPKSSDKTHTCPPCPAPELLGGPSVFLFPPKPKDTLMISRT PEVTCVVVDVSHEDPEVKFNWYVDGVEVHNAKTKPREEQYNSTYRVVSVLTVLHQDWLNGKEYKCKVSNKALPAPIEKT ISKAKGQPREPQVYTLPPSRDELTKNQVSLTCLVKGFYPSDIAVEWESNGQPENNYKTTPPVLDSDGSFFLYSKLTVDKS RWQQGNVFSCSVMHEALHNHYTQKSLSLSPGK |
| GFP-Fc (sfGFP) | SRATMETDTLLLWVLLLWVPGSTGHHHHHHGGSENLYFQGGSSKGEELFTGVVPILVELDGDVNGHKFSVRGEGEGDA TNGKLTLKFICTTGKLPVPWPTLVTTLTYGVQCFSRYPDHMKRHDFFKSAMPEGYVQERTISFKDDGTYKTRAEVKFEGD TLVNRIELKGIDFKEDGNILGHKLEYNFNSHNVYITADKQKNGIKANFKIRHNVEDGSVQLADHYQQNTPIGDGPVLLPDNH YLSTQSVLSKDPNEKRDHMVLLEFVTAAGITHGMDELYKGGSGSEPKSSDKTHTCPPCPAPELLGGPSVFLFPPKPKDTL MISRTPEVTCVVVDVSHEDPEVKFNWYVDGVEVHNAKTKPREEQYNSTYRVVSVLTVLHQDWLNGKEYKCKVSNKALP APIEKTISKAKGQPREPQVYTLPPSRDELTKNQVSLTCLVKGFYPSDIAVEWESNGQPENNYKTTPPVLDSDGSFFLYSKL TVDKSRWQQGNVFSCSVMHEALHNHYTQKSLSLSPGK |
| RFP-Fc (mRuby 2) | SRATMETDTLLLWVLLLWVPGSTGHHHHHHGGSENLYFQGGSVSKGEELIKENMRMKVVMEGSVNGHQFKCTGEGEG NPYMGTQTMRIKVIEGGPLPFAFDILATSFMYGSRTFIKYPKGIPDFFKQSFPEGFTWERVTRYEDGGVVTVMQDTSLED GCLVYHVQVRGVNFPSNGPVMQKKTKGWEPNTEMMYPADGGLRGYTHMALKVDGGGHLSCSFVTTYRSKKTVGNIKM PGIHAVDHRLERLEESDNEMFVVQREHAVAKFAGLGGGMDELYKGGSGSEPKSSDKTHTCPPCPAPELLGGPSVFLFPP KPKDTLMISRTPEVTCVVVDVSHEDPEVKFNWYVDGVEVHNAKTKPREEQYNSTYRVVSVLTVLHQDWLNGKEYKCKVS NKALPAPIEKTISKAKGQPREPQVYTLPPSRDELTKNQVSLTCLVKGFYPSDIAVEWESNGQPENNYKTTPPVLDSDGSFF LYSKLTVDKSRWQQGNVFSCSVMHEALHNHYTQKSLSLSPGK |
| A1F-Fc | METDTLLLWVLLLWVPGSTGKAELASEKPFRDCADVYQAGFNKSGIYTIYINNMPEPKKVFCNMDVNGGGWTVIQHRED GSLDFQRGWKEYKMGFGNPSGEYWLGNEFIFAITSQRQYMLRIELMDWEGNRAYSQYDRFHIGNEKQNYRLYLKGHTG TAGKQSSLILHGADFSTKDADNDNCMCKCALMLTGGWWFDACGPSNLNGMFYTAGQNHGKLNGIKWHYFKGPSYSLR STTMMIRPLDFGGSGGSEPKSSDKTHTCPPCPAPELLGGPSVFLFPPKPKDTLMISRTPEVTCVVVDVSHEDPEVKFNWY VDGVEVHNAKTKPREEQYNSTYRVVSVLTVLHQDWLNGKEYKCKVSNKALPAPIEKTISKAKGQPREPQVYTLPPSRDEL TKNQVSLTCLVKGFYPSDIAVEWESNGQPENNYKTTPPVLDSDGSFFLYSKLTVDKSRWQQGNVFSCSVMHEALHNHYT QKSLSLSPGKGGSHHHHHH |

**Supplementary Materials 1. Geometry specification in the .config files used in fusion protocol (using the WORMS protocol at https://github.com/willsheffler/worms).**

**D2 Dihedron:**
[('fc_binder',orient(None,'C')),('Monomer',orient('N','C')),('C2_N',orient('N',None))]
D2(c2=0, c2b=-1)

**D3 Dihedron:**
[('fc_binder',orient(None,'C')),('Monomer',orient('N','C')),('C3_N',orient('N',None))]
D3(c2=0, c2b=-1)

**T32 Tetrahedron:**
[('fc_binder',orient(None,'C')),('Monomer',orient('N','C')),('C3_N',orient('N',None))]
Tetrahedral(c2=0, c3=-1)

**O32 Octahedron:**
[('fc_binder',orient(None,'C')),('Monomer',orient('N','C')),('C3_N',orient('N',None))]
Octahedral(c2=0, c3=-1)

**O42 Octahedron:**
[('fc_binder',orient(None,'C')),('Monomer',orient('N','C')),('C4_N',orient('N',None))]
Octahedral(c2=0, c4=-1)

**I32 Icosahedron:**
[('fc_binder',orient(None,'C')),('Monomer',orient('N','C')),('C3_N',orient('N',None))]
Icosahedral(c2=0, c3=-1)

**I52 Icosahedron:**
[('fc_binder',orient(None,'C')),('Monomer',orient('N','C')),('C5_N',orient('N',None))]
Icosahedral(c2=0, c5=-1)

**Supplementary Materials 2. Example .json file database entry for each building block used in the helical fusion protocol (using the WORMS protocol at https://github.com/willsheffler/worms).**


```
[
{"file": "/path/to/fc_binder/file1.pdb",
 "name": "protein_a_d_domain" ,
 "class": ["fc_binder"],
 "type": "fc_binder" ,
 "connections": [
          {"chain": 1, "direction": "C", "residues":["-17:"]}
 ]
},

{"file": "/path/to/monomer/file1.pdb",
 "name": "dhr10" ,
 "class": ["monomer"],
 "type": "monomer" ,
 "connections": [
          {"chain": 1, "direction": "N", "residues":[":50"]},
          {"chain": 1, "direction": "C", "residues":["-150:"]}
  ]
},

{"file": "/path/to/cyclic_oligomer/file1.pdb",
 "name": "example_c2" ,
 "class": ["C2_N"],
 "type": "C2_N" ,
 "connections": [
          {"chain": 1, "direction": "N", "residues":[":50"]},
  ]
},

{"file": "/path/to/cyclic_oligomer/file2.pdb",
 "name": "example_c3" ,
 "class": ["C3_N"],
 "type": "C3_N" ,
 "connections": [
          {"chain": 1, "direction": "N", "residues":[":50"]},
  ]
}
]
```

**Supplementary Materials 3. Example command line command used to launch AbC fusion generation job (using the WORMS protocol at https://github.com/willsheffler/worms).**

PYTHONPATH="/home/rdd48/worms" python /path/to/generate_chains.py --config_file /path/to/config_file/see_fig_s2 --err_cutoff 0.5 --clash_cutoff 1.0 --database_files /path/to/database_files/see_fig_s1

**Supplementary Materials 4. Example .xml file used during post-helical fusion residue design.** Paths to designable residue files (resfiles) and symdef (symmetry definition) files provided on the command-line during the run.

```
<ROSETTASCRIPTS>
	<SCOREFXNS>
		<ScoreFunction name="sfx_hard_symm" weights="beta.wts" symmetric="1"  >
			<Reweight scoretype="res_type_constraint" weight="1.0" />
			<Reweight scoretype="aa_composition" weight="1.0" />
			<Reweight scoretype="coordinate_constraint" weight="1.00" />
		</ScoreFunction>
	</SCOREFXNS>
	<TASKOPERATIONS>
		<InitializeFromCommandline name="init" />
		<IncludeCurrent name="ic" />
		<RestrictIdentities name="nomutate_VIRTUAL" identities="XXX" prevent_repacking="1" />
		<LimitAromaChi2 name="limitaro" chi2max="110" chi2min="70" />
		<ReadResfile name="resfile_designable" filename="%%resfile%%" />
	</TASKOPERATIONS>
	<MOVERS>
	<SetupForSymmetry name="symmetry_setup" definition="%%symdef%%"></SetupForSymmetry>
		<SymPackRotamersMover name="design_rotamers_resfile" scorefxn="sfx_hard_symm"
task_operations="init,ic,limitaro,nomutate_VIRTUAL,resfile_designable"></SymPackRotamersMover>
	</MOVERS>
	<PROTOCOLS>
    <Add mover_name="symmetry_setup" />
		<Add mover_name="design_rotamers_resfile" />
	</PROTOCOLS>
</ROSETTASCRIPTS>
```

**Supplementary Materials 5. Example resfile (residue specification file) used to design AbC helical fusion outputs.** Designable residues were near the fusion junctions. Residues from the original building were occasionally restored by directly specifying them (e.g. residue 245 was a glutamate in the original building block).

```
NATRO
START
245 A PIKAA E
248 A PIKAA R
253 A APOLAR
265 A PIKAA D
272 A PIKAA AVIL
277 A PIKAA VIL
294 A PIKAA E
298 A PIKAA S
306 A APOLAR
307 A PIKAA AVIL
309 A APOLAR
311 A PIKAA ST
```

### Full References

15. Y. Hsia, R. Mout, W. Sheffler, N. I. Edman, I. Vulovic, Y.-J. Park, R. L. Redler, M. J. Bick, A. K. Bera, A. Courbet, A. Kang, T. J. Brunette, U. Nattermann, E. Tsai, A. Saleem, C. M. Chow, D. Ekiert, G. Bhabha, D. Veesler, D. Baker, Hierarchical design of multi-scale protein complexes by combinatorial assembly of oligomeric helical bundle and repeat protein building blocks (2020).

40. M. A. Tortorici, M. Beltramello, F. A. Lempp, D. Pinto, H. V. Dang, L. E. Rosen, M. McCallum, J. Bowen, A. Minola, S. Jaconi, F. Zatta, A. De Marco, B. Guarino, S. Bianchi, E. J. Lauron, H. Tucker, J. Zhou, A. Peter, C. Havenar-Daughton, J. A. Wojcechowskyj, J. B. Case, R. E. Chen, H. Kaiser, M. Montiel-Ruiz, M. Meury, N. Czudnochowski, R. Spreafico, J. Dillen, C. Ng, N. Sprugasci, K. Culap, F. Benigni, R. Abdelnabi, S.-Y. C. Foo, M. A. Schmid, E. Cameroni, A. Riva, A. Gabrieli, M. Galli, M. S. Pizzuto, J. Neyts, M. S. Diamond, H. W. Virgin, G. Snell, D. Corti, K. Fink, D. Veesler, Ultrapotent human antibodies protect against SARS-CoV-2 challenge via multiple mechanisms. *Science* (2020), doi:10.1126/science.abe3354.

41. L. Piccoli, Y.-J. Park, M. A. Tortorici, N. Czudnochowski, A. C. Walls, M. Beltramello, C. Silacci-Fregni, D. Pinto, L. E. Rosen, J. E. Bowen, O. J. Acton, S. Jaconi, B. Guarino, A. Minola, F. Zatta, N. Sprugasci, J. Bassi, A. Peter, A. De Marco, J. C. Nix, F. Mele, S. Jovic, B. F. Rodriguez, S. V. Gupta, F. Jin, G. Piumatti, G. Lo Presti, A. F. Pellanda, M. Biggiogero, M. Tarkowski, M. S. Pizzuto, E. Cameroni, C. Havenar-Daughton, M. Smithey, D. Hong, V. Lepori, E. Albanese, A. Ceschi, E. Bernasconi, L. Elzi, P. Ferrari, C. Garzoni, A. Riva, G. Snell, F. Sallusto, K. Fink, H. W. Virgin, A. Lanzavecchia, D. Corti, D. Veesler, Mapping Neutralizing and Immunodominant Sites on the SARS-CoV-2 Spike Receptor-Binding Domain by Structure-Guided High-Resolution Serology. *Cell*. **183**, 1024–1042.e21 (2020).

49. J. K. Leman, B. D. Weitzner, S. M. Lewis, J. Adolf-Bryfogle, N. Alam, R. F. Alford, M. Aprahamian, D. Baker, K. A. Barlow, P. Barth, B. Basanta, B. J. Bender, K. Blacklock, J. Bonet, S. E. Boyken, P. Bradley, C. Bystroff, P. Conway, S. Cooper, B. E. Correia, B. Coventry, R. Das, R. M. De Jong, F. DiMaio, L. Dsilva, R. Dunbrack, A. S. Ford, B. Frenz, D. Y. Fu, C. Geniesse, L. Goldschmidt, R. Gowthaman, J. J. Gray, D. Gront, S. Guffy, S. Horowitz, P.-S. Huang, T. Huber, T. M. Jacobs, J. R. Jeliazkov, D. K. Johnson, K. Kappel, J. Karanicolas, H. Khakzad, K. R. Khar, S. D. Khare, F. Khatib, A. Khramushin, I. C. King, R. Kleffner, B. Koepnick, T. Kortemme, G. Kuenze, B. Kuhlman, D. Kuroda, J. W. Labonte, J. K. Lai, G. Lapidoth, A. Leaver-Fay, S. Lindert, T. Linsky, N. London, J. H. Lubin, S. Lyskov, J. Maguire, L. Malmström, E. Marcos, O. Marcu, N. A. Marze, J. Meiler, R. Moretti, V. K. Mulligan, S. Nerli, C. Norn, S. Ó'Conchúir, N. Ollikainen, S. Ovchinnikov, M. S. Pacella, X. Pan, H. Park, R. E. Pavlovicz, M. Pethe, B. G. Pierce, K. B. Pilla, B. Raveh, P. Douglas Renfrew, S. S. Roy Burman, A. Rubenstein, M. F. Sauer, A. Scheck, W. Schief, O. Schueler-Furman, Y. Sedan, A. M. Sevy, N. G. Sgourakis, L. Shi, J. B. Siegel, D.-A. Silva, S. Smith, Y. Song, A. Stein, M. Szegedy, F. D. Teets, S. B. Thyme, R. Y.-R. Wang, A. Watkins, L. Zimmerman, R. Bonneau, Macromolecular modeling and design in Rosetta: recent methods and frameworks. *Nat. Methods*. **17**, 665–680 (2020).

55. K. Zhang, Gctf: Real-time CTF determination and correction. *J. Struct. Biol.* **193**, 1–12

(2016).

61. K. L. DeCicco-Skinner, G. H. Henry, C. Cataisson, T. Tabib, J. C. Gwilliam, N. J. Watson, E. M. Bullwinkle, L. Falkenburg, R. C. O'Neill, A. Morin, J. S. Wiest, Endothelial cell tube formation assay for the in vitro study of angiogenesis. *J. Vis. Exp.*, e51312 (2014).

62. K. H. D. Crawford, R. Eguia, A. S. Dingens, A. N. Loes, K. D. Malone, C. R. Wolf, H. Y. Chu, M. A. Tortorici, D. Veesler, M. Murphy, D. Pettie, N. P. King, A. B. Balazs, J. D. Bloom, Protocol and Reagents for Pseudotyping Lentiviral Particles with SARS-CoV-2 Spike Protein for Neutralization Assays. *Viruses*. **12** (2020), doi:10.3390/v12050513.
